# Cytoplasmic crowding acts as a porous medium reducing macromolecule diffusion

**DOI:** 10.1101/2025.07.28.667145

**Authors:** Olivier Destrian, Nicolas Moisan, René-Marc Mege, Benoit Ladoux, Benoit Goyeau, Morgan Chabanon

## Abstract

Intracellular transport of macromolecules is crucial for the proper functioning of most cellular processes. Although intracellular crowding is known to strongly alter macromolecule mobility, how cytoplasmic structures physically modulate diffusion remains largely unexplored. Here we investigated the mechanisms by which cytoplasmic crowding controls diffusivity using live-cell experiments and porous media modeling approaches. Confocal microscopy combined with fluorescence recovery after photobleaching (FRAP) and fluorescence correlation spectroscopy (FCS) measurements revealed an anti-correlation between free-GFP diffusivity and the heterogeneous cytoplasmic structure abundance in live mammalian cells. This motivated the development of a multiscale model, where the cytoplasm is treated as a hierarchical porous medium with nanometric and micrometric obstacles. Numerically solving the model allowed us to predict the effective cytoplasmic diffusion coefficient for various obstacle volume fractions, and to identify tortuous and porous hydrodynamic hindrances as key diffusion reduction mechanisms. Comparison with our experimental results highlighted the importance of hydrodynamic interactions between diffusing molecules and nanometric obstacles. Importantly, we found that the effective cytoplasmic diffusivity was not dependent on specific intracellular regions but rather on the local intracellular obstacle volume fraction. Finally, the model was extended to predict the diffusivity of larger macromolecules, showing excellent agreement with literature data for several macromolecules and cell lines. This study provides new insights into the physical mechanisms impeding intracellular diffusion, demonstrating the potential of porous media modeling approaches to predict transport mechanisms in dynamic or heterogeneous intracellular structures, as in cell motility, blebbing, and apoptosis.

## INTRODUCTION

The intracellular space is packed with various membranous and proteinic structures that occupy up to 40% of the cytoplasm [1, 2, 3]. The presence of these structures reduces the accessible volume for dilute macromolecules and hinders their mobility, effectively inducing crowding. Yet, numerous cellular processes rely on macromolecule passive transport, including protein synthesis [4], actin turnover in cell migration [5], and caspase dynamics during apoptosis [6].

Although intracellular crowding is known to drastically alter intracellular signaling and biogenesis [7, 8], how cytoplasmic structures impact globular protein diffusion remains unclear. Diverse organelle and polymeric molecular complexes are thought to hamper cytoplasmic diffusion, including the cytoskeleton [9, 10, 11], the endoplasmic reticulum [12], and densely packed ribosomes [13, 14]. Additionally, the intracellular organization is highly dynamic and responds to biochemical cues [13, 5], and mechanical stimuli such as osmotic shocks [15, 16, 17] and substrate stiffness [18, 19], potentially modulating crowding and diffusion dynamics.

Macromolecule transport mechanisms in the cytoplasm include convection, active transport on cytoskeletal filaments, and diffusion [20]. As diffusion does not consume energy it has been proposed that cells optimize this transport mechanism [21]. However, in contrast to diffusion in diluted solutions which follow the linear Stokes-Einstein law, macromolecular cytoplasmic diffusion exhibits complex non-linear behaviors: (i) Compared to water, it is strongly reduced and dependent on macromolecule size [22, 23, 24, 25, 26] (ii) It is non-uniform inside the cytoplasm [27, 28, 29] (iii) It is non-Brownian and displays a characteristic transient-subdiffusive behavior [30, 31, 32].

Understanding the mechanisms behind cytoplasmic diffusion remains a topic of active research. While electrostatic, hydrophobic, and specific interactions are elements of answers for charged and reactive particles [33, 34, 35], they cannot explain these complex behaviors for inert macromolecules such as dextrans and GFP-oligomers [30, 23, 26, 25]. Instead, diffusion of inert particles is likely determined by steric interactions with structures such as the cytoskeleton [36, 37] and arrays of ribosomes [13]. Some attempts have been made to link the diffusive behavior of inert particles to immobile intracellular structures [29] and to crowders [38]. Moreover, static cytoplasmic structures have been reported as a probable source of transient-subdiffusion [30, 23]. Some modeling works have focused on the cytoplasmic diffusion of infinitely small particles [34], or have neglected hydrodynamic effects [36]. Yet, others found that the effect of hydrodynamic interactions on finite-size particles in simple prokaryotic cells could be important [38]. To our knowledge, no physical model has yet been proposed that quantitatively links eukaryotic intracellular structures to the effective cytoplasmic diffusivity of macromolecules.

In this work, we propose to formalize the relationship between cytoplasmic macromolecular transport and intracellular structures abundance using an upscaling framework initially developed for transport phenomena in porous media [39]. This framework is well suited to determine average transport equations and effective transport coefficients – such as permeability or effective diffusivity – based on the knowledge of the transport mechanisms at the pore scale. This approach has been transposed successfully in soft matter physics. Notably, the study of hydrodynamic interactions between a hydrogel matrix and diffusing macromolecules [40, 41] was originally based on the seminal work of Brinkman on flows in porous media [42]. Additionally, recent studies demonstrated the potential of porous media modeling approaches to describe cell biophysical processes. Cytoplasm was shown to behave as a poroelastic material as described by Biot’s theory [43]. During blebbing, the cytosolic flow through the cortex was modeled using Darcy’s transport equation of viscous flow in porous media [44]. Cytoplasmic transient-subdiffusion was also recovered using a porous media description [23, 36]. Altogether, these studies point to the potential of porous media models in intracellular biophysics.

Here, we present a multiscale model of intracellular macromolecule diffusion informed by live-cell fluorescence experiments to quantify diffusivity and structures abundance in the cytoplasm. Cells expressing free-GFP were imaged using confocal microscopy to map the abundance of structures in several cytoplasmic regions, revealing a spatially heterogeneous level of crowding. GFP diffusivity was measured in each region and for several medium osmolarities, revealing an anti-correlation between diffusivity and the local abundance of nanometric structures. Following this, a multiscale diffusion model based on a porous media formulation was proposed. Comparison between model predictions and experimental data highlighted that the cytoplasm behaves as a nano-porous medium, inducing a heterogeneous and scale-dependent diffusivity reduction through two mechanisms: a tortuous hindrance and a hydrodynamic drag increase. Finally, the model generality was demonstrated by validating its predictions of diffusivity for various macromolecules sizes against independent measurements from the literature.

Taken together, this work suggests that (i) the diffusivity of macromolecules in cells follows a universal law that depends on the volume fraction of nanometric structures and on macromolecule size, and (ii) the cytoplasmic transport properties can be accurately predicted by porous media models.

## RESULTS

### Intracellular structures induce a heterogeneous crowding

To study how intracellular structures influence macromolecule diffusion, we first characterized the distribution of accessible and excluded volume for macromolecules in MDCK cells, used as a mammalian cell model. All experiments were carried out at low confluence, on large edge cells with a projected area typically between 2500 and 10000 *µm*^2^ (Fig. S1). Intracellular organization of cells was examined using differential interference contrast microscopy (Fig. 1A), and confocal microscopy of immunostained cells (Fig. 1B), showing that structures are heterogeneously distributed in the cytoplasm.

**Figure 1.**
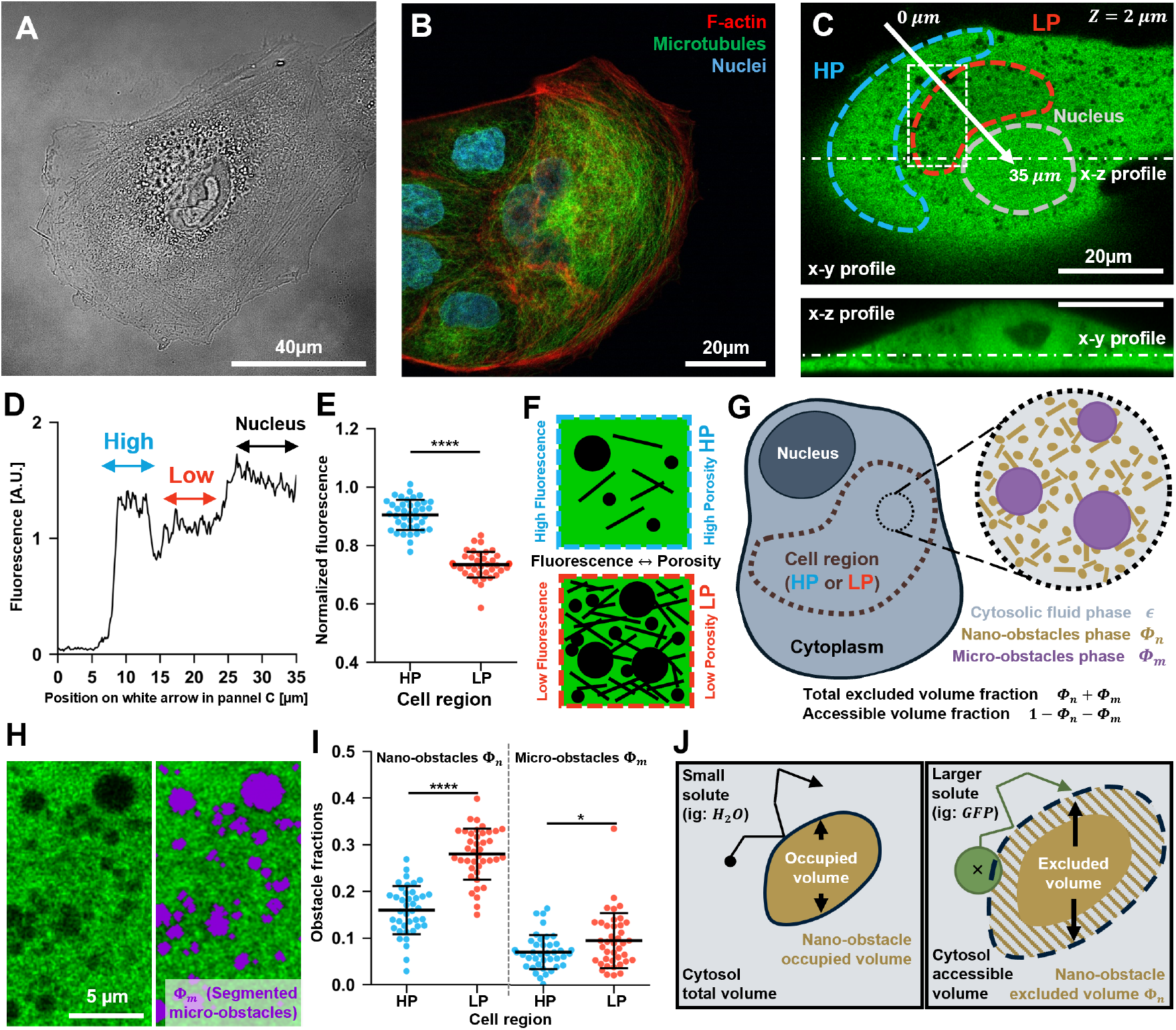
Characterization of MDCK intracellular architecture using optical microscopy. The cytoplasm features heterogeneous regions with different obstacle density, akin to porous media. (A) Differential interference contrast image of a cell, with perinuclear accumulation of heterogeneities. (B) Confocal image of a fixed cell stained for F-actin (red), microtubules (green) and nuclei (blue). (C) Confocal image of free-GFP fluorescence, which is highly heterogeneous in the cell. (D) GFP signal on the white arrow present in C. (E) GFP signal in the HP and LP regions of cells normalized by nucleoplasmic signal. (F) Schematic of how GFP fluorescence signal in HP and LP regions is linked to a higher or lower accessible volume for GFP. (G) Schematic of the cytoplasm crowded by nanometric and micrometric sized obstacles. (H) Zoom on the white rectangle present in C. Large dark spheroids are present (left). Their volume is estimated by image segmentation (right). (I) Micro- and nano-obstacle fractions inferred from normalized fluorescence and segmentation data. (J) Schematic showing that the volume nano-obstacles exclude for GFP is higher than the volume they occupy due to the non-negligible size of GFP. (E and I) N cells=40, horizontal bars are mean and *±*STD. Welch’s t-test: \**p* < 5*e* − 2 ; \*\**p* < 1*e* − 2; \*\*\**p* < 1*e* − 3 ; \*\*\*\**p* < 1*e* − 4. somes 12.5*nm*), Φ_*n*_ is larger than the volume fraction that the nano-obstacles physically occupy (Fig. 1J). The total excluded volume fraction for GFP is Φ = Φ_*m*_ + Φ_*n*_, so the volume fraction accessible for GFP is 1 − Φ (Fig. 1G).

To quantify the overall structure distribution, we used free-GFP fluorescence as a probe of accessible volume for macromolecule diffusion. Free-GFP is inert and highly mobile [45] and its distribution in the fluid phase of the cytoplasm – the cytosol – is assumed homogeneous. Additionally, weighting 27kDa, GFP can freely diffuse in the nucleus [46] (Fig. S2) and has a size close to mid-sized proteins such as G-actin (42kDa) or caspases (25-60kDa). Thus, in our experiments GFP fluorescence intensity indicates the distribution of the accessible volume fraction to many globular proteins [47].

Cytoplasmic structures with negligible mobility compared to GFP reduce the volume accessible to the mass-center of globular proteins such as GFP. These structures, refered here as obstacles to diffusion, notably include organelles, membranous networks, cytoskeletal fibers, and macromolecular complexes such as ribosomes. Importantly, molecules that diffuse much faster than GFP are not considered as obstacles but as viscogens that increase the cytosol viscosity [7]. Due to their great mobility, we assume that the overall viscogens distribution in the cytosol is homogeneous and thus does not impact GFP fluorescence signal distribution. We infected MDCK cells to stably express TurboGFP and conducted Z-stack confocal imaging of the cells to characterize the intracellular organization. We observed that GFP fluorescence is highly heterogeneous within cells (Fig. 1C, Movie S1), showing dark spots of a few microns and important fluorescence fluctuations outside of these spots (Fig. 1H left). In Fig. 1D, GFP signal was plotted as a function of the position along the white arrow across the cell presented in Fig. 1C. We observed several fluorescence plateaus: the nucleoplasm with the highest intensity and two cytoplasmic regions of high and low intensity respectively.

From these observations, we studied cytoplasmic regions with either high fluorescence thus large accessible volume to GFP, or lower fluorescence thus smaller accessible volume, that we refer to as high porosity (HP) and low porosity (LP) regions respectively (Fig. 1C-F). To ease comparison between cells, the HP and LP fluorescence from each cell was normalized by the nucleoplasm fluorescence, which is high and homogeneous outside of the nucleolus [47, 46]. We found HP and LP normalized fluorescence values about 0.9 and 0.75 (Fig. 1I).

Next, we quantified the abundance of obstacles in the HP and LP regions. As seen in Fig. 1H, the overall fluorescence level is impacted not only by micron-size dark spheroids (>100nm) that are well resolved by confocal microscopy, but also by the fluorescence signal being lower in the more homogeneous subregions surrounding these objects. These subregions correspond to locations where smaller and unresolved obstacles (*<*100nm) are abundant. These two types of obstacles are interpreted as micro- and nano-obstacles respectively.

To quantify the volume fraction excluded by micro-obstacles Φ_*m*_, we used a custom segmentation procedure (Fig. 1H and SI Appendix 1E). Given that the accessible nucleoplasmic volume fraction for GFP is about 85% [48], Φ_*n*_ could then be inferred from Φ_*m*_ and fluorescence data (Fig. 1E). Importantly, as GFP radius *R*_GFP_ = 2.3*nm* [49] is non-negligible compared to typical nano-obstacles radii (F-actin 4*nm*, ribo-

With these definitions, the total excluded volume fraction Φ = Φ_*n*_ + Φ_*m*_ is significantly higher in LP than in HP regions (Fig. 1I). Nano-obstacles Φ_*n*_ are responsible for most of the excluded volume in both HP and LP regions, while micro-obstacles Φ_*m*_ are scarce. The volume fraction occupied by the nano-obstacles – which does not depend on GFP size – was evaluated to be about half of Φ_*n*_ (Fig. S3). These results suggest that the cytoplasm is highly and heterogeneously crowded by structures at different scales, bearing similarities with hierarchical porous media [50].

### Volume fraction accessible to macromolecules can be tuned with medium osmolarity

Next, we sought to modulate the accessible volume for GFP particles inside cells. Hyper-osmotic shocks induce a cell volume reduction due to water leakage, increasing macromolecular concentration and thus the abundance of cytoplasmic obstacles (Fig. 2A) [15]. To increase the osmolarity above the physiological level (300mOsm), sucrose was added to reach osmolarities of 450, 600, and 900mOsm (SI Appendix 1C). Confocal z-stack images of GFP fluorescence before and after sucrose addition show a reduction in cell height with a conserved projected area, resulting in a net cell volume reduction (Fig. 2B).

**Figure 2.**
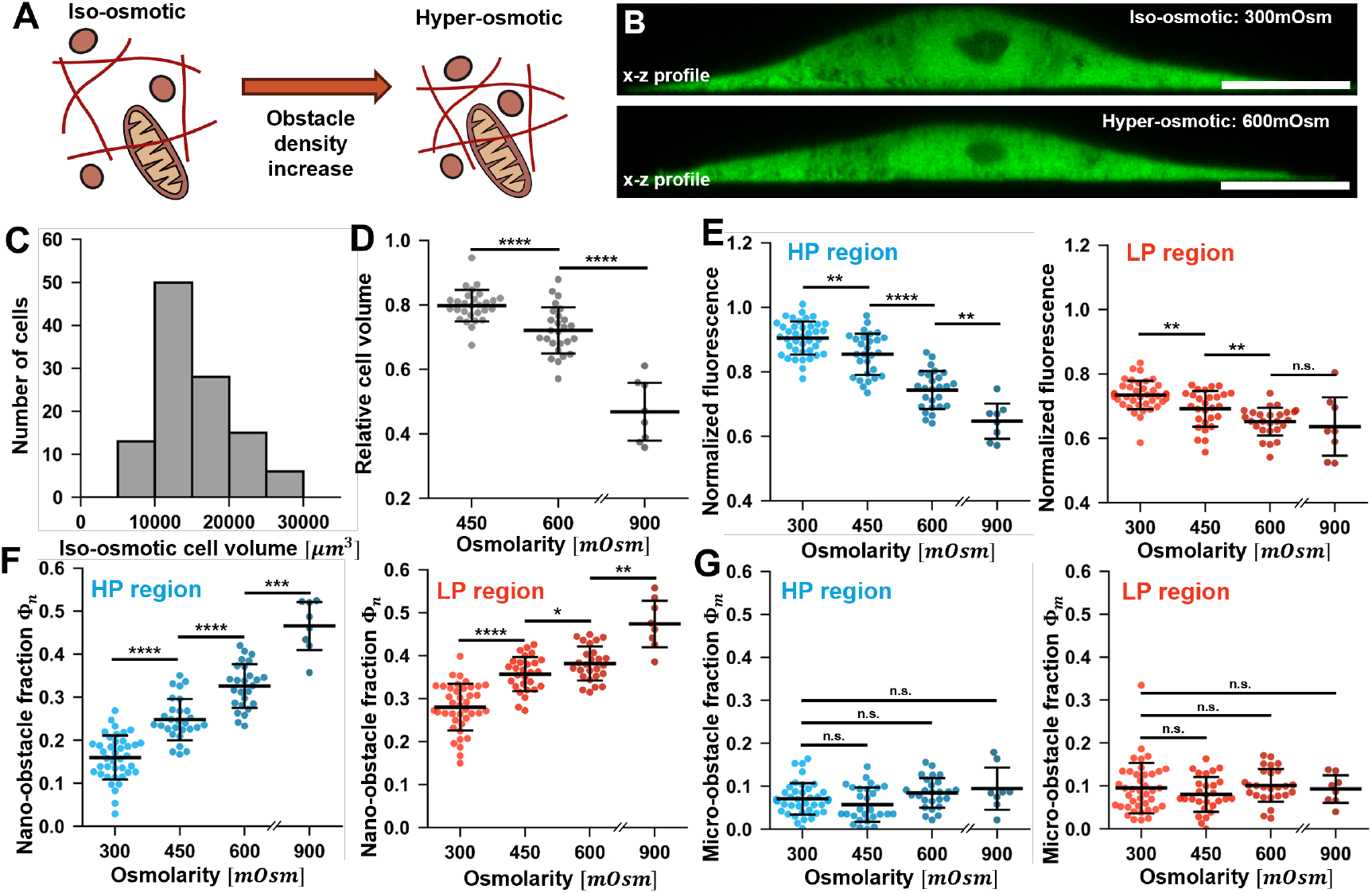
Quantification of cell volume and cytoplasmic obstacles volume fractions for several osmotic conditions. Obstacle abundance is greater in LP than in HP regions and is increased upon hyper-osmotic shock. (A) Schematic showing how hyperosmolarity increases the concentration of cytoplasmic obstacles. (B) Confocal fluorescence images of free-GFP in a living cell (same cell as in Fig. 1C) in iso-osmotic condition (300 mOsm) and after immersion in a hyper-osmotic (600mOsm) medium. (C) Measured cell volume in iso-osmotic condition. (D) Relative cell volume after/before osmotic shock. (E) Cytoplasmic fluorescence (normalized by nucleoplasm fluorescence) in HP regions (left) and LP regions (right). (F) Inferred values for nano-obstacles excluded volume fractions in HP regions (left) and LP regions (right). (G) Inferred values for micro-obstacles excluded volume fractions, in HP regions (left) and LP regions (right). (D-G) Each dot corresponds to a cell, with N cells=40 for 300mOsm, 28 for 450mOsm, 26 for 600mOsm and 8 for 900mOsm. Horizontal bars represent mean and *±*STD. Welch’s t-test: \**p* < 5*e* − 2 ; \*\**p* < 1*e* − 2; \*\*\**p* < 1*e* − 3 ; \*\*\*\**p* < 1*e* − 4.

Cell volume before and after osmotic shock was estimated using a custom segmentation procedure on Z-stack data (SI Appendix 1F, Fig. S4). Iso-osmotic cell volume was typically between 10000*µm*^3^ and 20000*µm*^3^ (Fig. 2C). After shock, cell volume was reduced by about 20% *±* 5% for 450 mOsm, about 28% *±* 7% for 600 mOsm, and about 53% *±* 9% for 900 mOsm (Fig. 2D), consistently with previous results [15].

The effect of osmolarity on GFP fluorescence was also estimated using Z-stack images. Though absolute fluorescence is increased after shock due to the increase of GFP concentration (Fig. S5), the cytoplasm fluorescence normalized by nucleoplasm fluorescence decreases with osmolarity in both HP and LP regions (Fig. 2E). Then, the excluded volume fractions Φ_*n*_ and Φ_*m*_ were inferred from the GFP fluorescence intensity. As in the previous subsection, the nucleus was taken as a reference for fluorescence and accessible volume fraction. The nucleoplasmic accessible volume fraction in iso-osmotic condition was estimated around 85% and adjusted for each osmotic condition by assuming a conserved nucleoplasmic chromatin content (SI Appendix 1E and Fig. S6). As seen in Fig. 2F, the excluded volume fraction due to nano-obstacles Φ_*n*_ almost doubles between 300 and 900mOsm, both in HP and LP regions. On the contrary, the effect of micro-obstacles on the excluded volume fraction is nearly unchanged with osmotic stress, suggesting that these structures are sensitive to osmotic stress (Fig. 2G). While Φ_*m*_ is almost constant within the cell, Φ_*n*_ is significantly larger in LP than in HP regions up 600mOsm. At 900mOsm it homogenizes, suggesting that HP regions are more strongly affected by osmotic stress.

These results indicate that the previously defined HP and LP regions are still distinguishable after mild hyper-osmotic shocks. Additionally, they show that hyper-osmotic shocks can tune the volume fraction excluded by cytoplasmic structures and particularly by nano-obstacles, providing us with a tool to probe a broader range of obstacle densities.

### Cytoplasmic diffusivity of free-GFP is anti-correlated with nano-obstacles abundance

The above results suggest that intracellular crowding is spatially heterogeneous and is increased in hyperosmotic conditions. We thus sought to examine consequences on cytoplasmic diffusion of inert macromolecules.

To measure GFP fast diffusion – assumed representative of mid-size globular proteins – we developed a specific FRAP protocol. Briefly, after applying a Gaussian bleaching spot at the center of a LP or HP region, fluorescence recovery was imaged at 50Hz over a 25*µm ×* 1*µm* rectangle. Rectangle lines were merged into a 1D chronograph, which was then used to fit a Gaussian diffusion model, yielding an estimate for GFP effective diffusivity *D*_exp_ (Fig. 3A, SI Appendix 1G).

**Figure 3.**
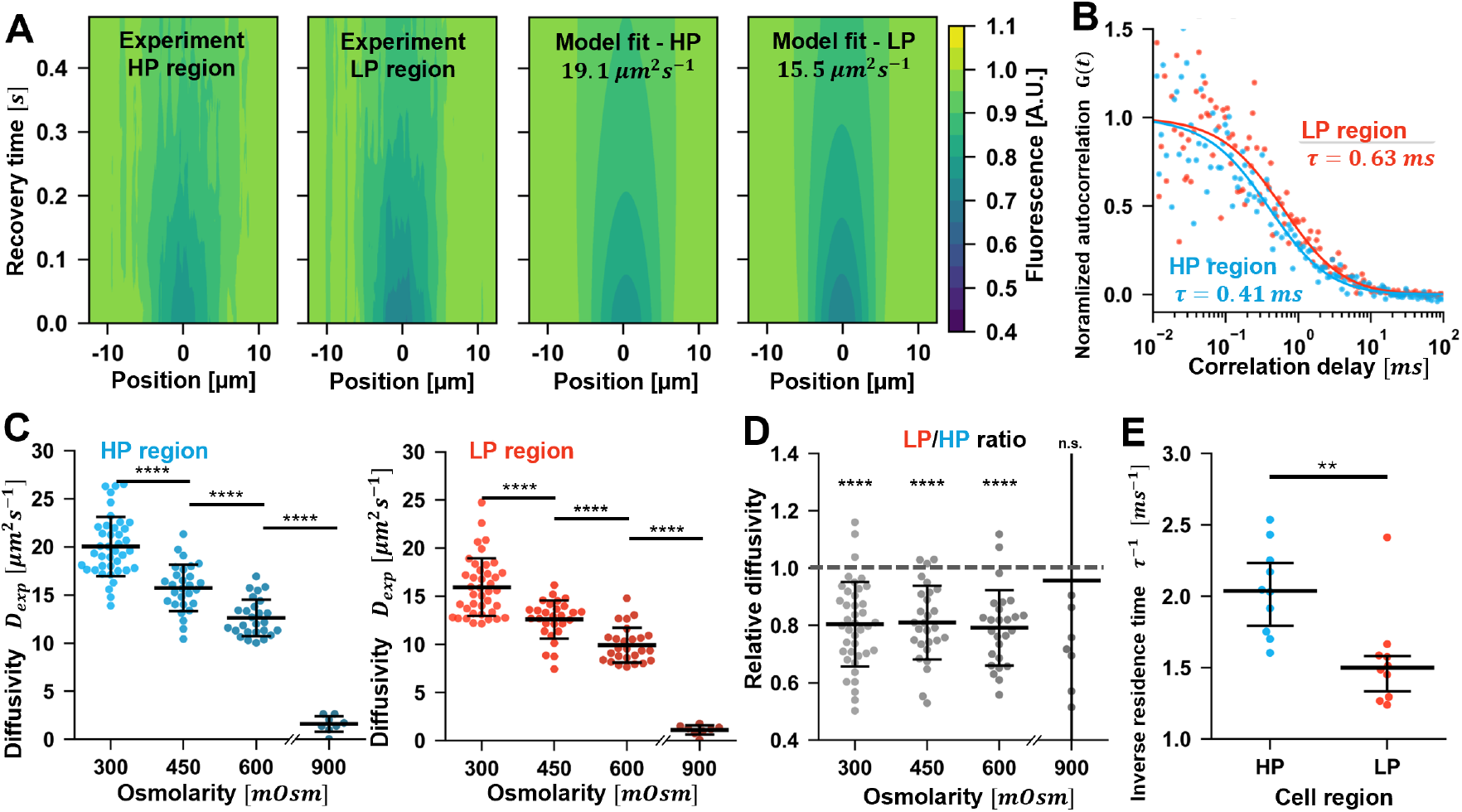
Estimation of free-GFP diffusivity with FRAP and FCS techniques in HP and LP regions for several osmotic conditions. GFP diffusivity correlates with the local accessible volume fraction of the cytoplasm. (A) Normalized fluorescence chronographs of FRAP recovery in the HP and LP regions of a cell (left) and associated FRAP diffusion model fits (right). (B) Experimental FCS autocorrelation curves in the HP and LP regions of a cell (dots) and normalized autocorrelation curves from FCS diffusion model fit (curves). (C) GFP diffusivity *D*_exp_ estimated from FRAP for HP (left) and LP regions (right). (D) GFP relative LP/HP diffusivity for each cell, from FRAP results in panel C. (E) GFP inverse residence time from FCS for HP and LP regions in iso-osmotic condition. The inverse residence time is proportional to diffusivity for a given setup. (C-E) Each dot corresponds to a cell, with N=40 for 300mOsm FRAP, 28 for 450mOsm, 26 for 600mOsm and 8 for 900mOsm and N=10 for 300mOsm FCS. Horizontal bars are mean and *±*STD. Welch’s t-test (for C and E) and One-sample t-test with test value=1.0 (for D): \**p* < 5*e* − 2 ; \*\**p* < 1*e* − 2; \*\*\**p* < 1*e* − 3 ; \*\*\*\**p* < 1*e* − 4.

Fig. 3C shows GFP diffusivity measured by FRAP in HP and LP regions, as a function of osmolarity. *D*_exp_ values were significantly below 87*µm*^2^*s*^−1^, the diffusivity of GFP in water [23, 51, 37], and diffusivity in both regions was strongly reduced with increasing osmolarity. At 900mOsm, *D*_exp_ drops suddenly in both regions, suggesting a transition in the diffusion regime above a crowding threshold.

Interestingly, cytoplasmic diffusivity is lower in LP than in HP regions. The difference is very clear when comparing regions in each cell (Fig. 3D), with about 20% difference in *D*_exp_ for several osmolarities. To confirm this observation, FCS measurements were also conducted in iso-osmotic conditions (Fig. 3B, SI Appendix 1H). As seen in Fig. 3E, results obtained for GFP inverse residence time – which is proportional to diffusivity for a given microscope point spread function – confirm the approximately 20% diffusivity reduction in LP regions compared to HP.

To investigate if structures such as cytoskeletal filaments were major sources of diffusional hindrance, we studied cells treated with cytoskeleton depolymerizing drugs (SI Appendix 2, Fig. S7-S13). Though important morphological changes were observed, F-actin and microtubules disruption did not drastically alter GFP diffusivity. This suggests that in the bulk cytoplasm regions studied, most of the diffusional hindrance arises from the presence of numerous other nano-obstacles such as membranous networks and ribosomes.

Together with previous sections data, these results show that the diffusivity of macromolecules is anticorrelated with nano-obstacles abundance and thus with the level of crowding.

### Multiscale modeling of macromolecule cytoplasmic diffusion

Our experimental results show that GFP diffusivity *D*_exp_ is correlated with the accessible volume fraction 1 − Φ, in several cytoplasmic regions and osmotic conditions. To investigate the mechanisms underlying this correlation, we derived a multiscale model of macromolecule diffusion in the cytoplasm based on a porous media modeling framework (SI Appendix 3 for detailed derivation).

Based on our observations (Fig. 1G-I), we decomposed the intracellular medium into three spatial scales (Fig. 4A): the cytoplasm-scale (∼ 10*µm*), the micro-scale (∼1*µm*), and the nano-scale (∼ 100*nm*).

**Figure 4.**
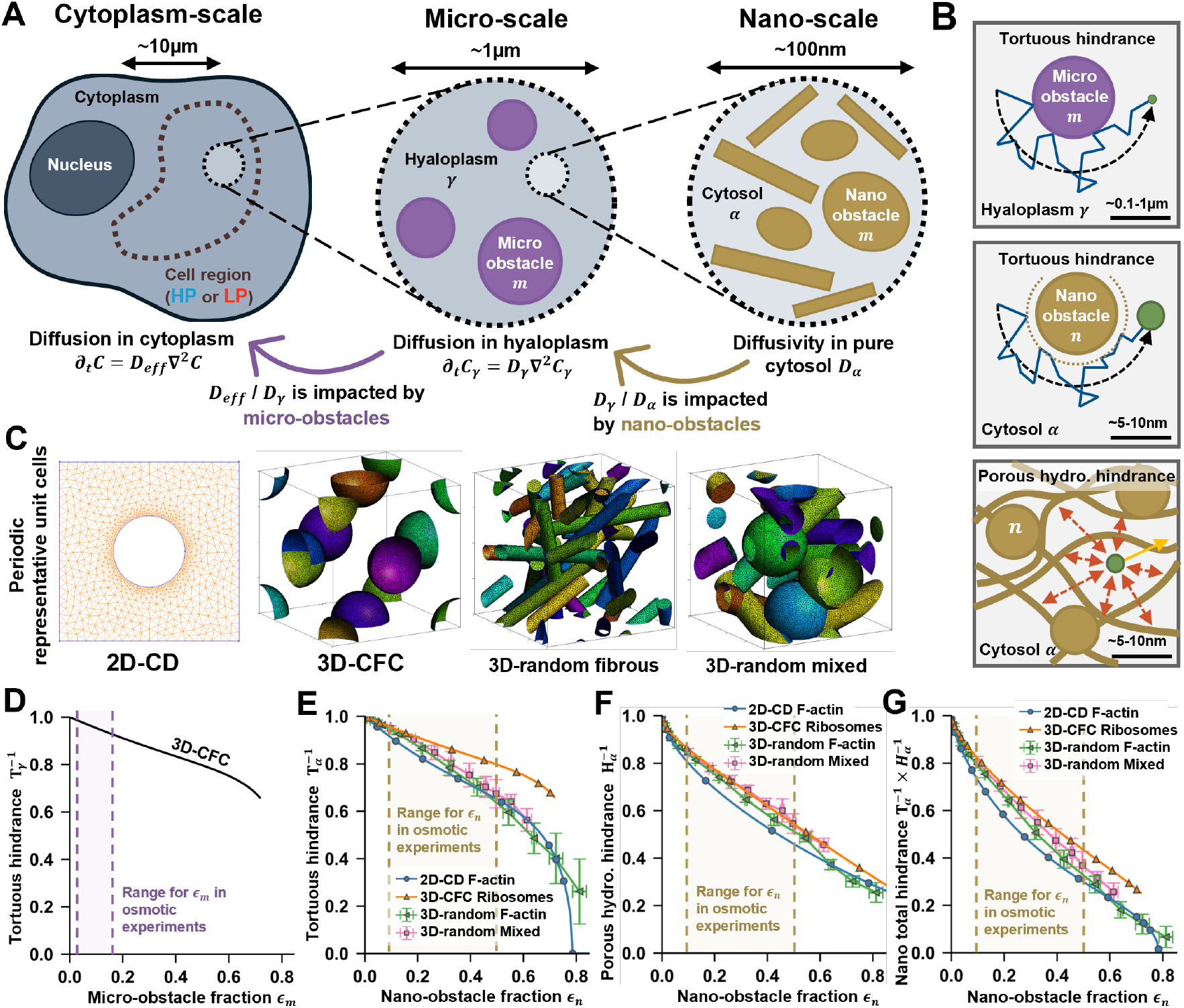
Presentation of the multiscale model for the cytoplasmic diffusion of free-GFP. Nano-obstacles result in tortuous and porous hydrodynamic hindrances that can strongly reduce the diffusivity of particles with GFP size. (A) Schematic of the cytoplasm decomposition into three scales. (B) Schematic showing the sources of diffusional hindrance arising from micro-obstacles (in purple) and nano-obstacles (in brown). (C) Examples of periodic meshes generated for the Representative Elementary Volumes (REV) used for numerical resolution of closure problems. (D) Dependence of 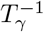 on *ε*_*m*_. (E) Dependence of 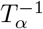 on *ε*_*n*_. (F) Dependence of 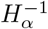 on *ε*_*n*_. (G) Dependence of 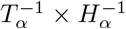 on *ε*_*n*_. (E-G) Errorbars are mean*±*STD for N*≥*3 random geometries.

At the cytoplasm-scale, the cytoplasm is considered an equivalent continuous medium where GFP effective diffusivity *D*_eff_ is impacted by obstacles at smaller scales. *D*_eff_ is the model counterpart of the experimentally measured FRAP diffusivity *D*_exp_ (Fig. 3C).

At the micro-scale, micron size objects surrounded by the hyaloplasm are considered. These obstacles represent large organelles and vesicles, and are considered non-deformable, impermeable, and immobile. We define *ε*_*m*_ the volume fraction excluded by micro-obstacles in the cytoplasm. The hyaloplasm is itself an equivalent continuous medium in which GFP has an effective diffusivity *D*_*γ*_ driven by sub-micrometric physics. The hyaloplasmic volume fraction in the cytoplasm is 1 − *ε*_*m*_.

At the nano-scale, nanometric obstacles (such as cytoskeletal filaments, ribosomes, and membranous networks) surrounded by the liquid cytosol are considered. Nano-obstacles are considered non-deformable, impermeable, and immobile on GFP diffusion timescales (SI Appendix 3B). These obstacles are surrounded by the cytosol, considered as a continuous medium constituted of water and fast-diffusing solutes. In pure cytosol, GFP has a local diffusivity *D*_*α*_. At this scale the volume fraction excluded by nano-obstacles for GFP is noted *ε*_*n*_, so the accessible cytosol volume fraction is 1 − *ε*_*n*_. With this framework, *D*_eff_ and *D*_*α*_ are linked by

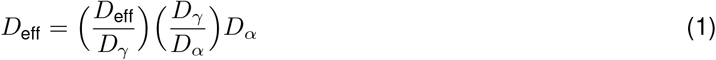

where diffusional hindrances induced by micro-obstacles, nano-obstacles, and viscogens are present in *D*_eff_*/D*_*γ*_, *D*_*γ*_ */D*_*α*_ and *D*_*α*_ respectively.

Micro-obstacles are much larger than GFP, thus GFP transport at the micro-scale can be modeled as a classic diffusion process in a rigid, impermeable porous medium. This is a well-known porous media problem, of which a classical result is that the effective diffusivity is reduced due to a tortuous hindrance *T*_*γ*_ at the micro-scale (Fig. 4B, top panel)

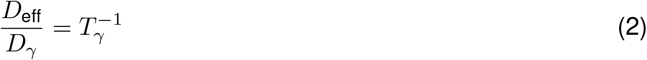

Using the volume averaging method [39], *T*_*γ*_ can be computed by solving a system of partial differential equations – the closure problem – on a representative elementary volume of the micro-scale (SI Appendix 3E).

Conversely, nano-obstacles – including F-actin and ribosomes – are less than an order of magnitude larger than GFP *R*_GFP_ = 2.3*nm*. Thus diffusion is not only impacted by tortuosity but also by hydrodynamic interactions between GFP and nearby nano-obstacles (Fig. 4B, middle and bottom panels, SI Appendix 3C). In line with previous gel inspired models used to study cytoplasmic properties [43, 52], we turned to solute diffusion models in hydrogels where the effective diffusivity *D*_*γ*_ is reduced by the multiplicative contributions of nano-scale tortuosity *T*_*α*_ and porous hydrodynamic hindrance *H*_*α*_ [53, 54]

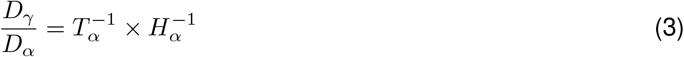

*T*_*α*_ is estimated by solving a closure problem – similarly to *T*_*γ*_ – on a nano-scale REV. However, as nano-obstacles exclude a larger volume than they occupy, solving is not conducted on the cytosol physical volume but on the accessible fraction for GFP mass-center 1 − *ε*_*n*_ (Fig. 1J and SI Appendix 3D).

The porous hydrodynamic hindrance *H*_*α*_ corresponds to a hydrogel-induced fluid friction increase. When a particle diffuses in a porous medium such as a gel, the porous structure hinders the embedded fluid flow, increasing the resulting viscous force exerted on the particle and reducing its diffusivity. The porous hydrodynamic drag was originally determined by upscaling Stokes flow in a porous medium [42, 55], and then adapted to solute diffusion in hydrogels [41, 56, 57]. The expression for 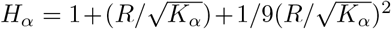 involves the ratio between the particle radius *R* and the square root of the medium permeability 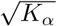. Importantly 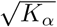 is also a common estimate of the pore size, and *H*_*α*_ → 1 when *R* becomes negligible compared to pore size, as for GFP at the micro-scale. We evaluated *K*_*α*_ with the volume averaging method by upscaling Stokes flow at the nano-scale. As water particles are very small compared to nano-obstacles, the obtained closure problem is solved on the cytosol physical volume (Fig. 1J).

Closure problems for *T*_*γ*_, *T*_*α*_ and *H*_*α*_ were solved for various obstacle fractions using the finite element method with FEniCS on periodic unit cells meshed using Gmsh (Fig. 4C, S14-S17 and SI Appendix 3F). For *H*_*α*_, results were expressed as a function of *ε*_*n*_ using simple geometrical models (SI Appendix 3C). The results (Fig. 4D-G) indeed show that higher obstacles volume fractions *ε*_*n*_ and *ε*_*m*_ reduce effective diffusivity *D*_eff_. Within the range of studied osmotic conditions, micro-obstacles reduce GFP diffusivity by less than 10%, while nano-obstacles – generating porous hydrodynamic hindrance *H*_*α*_ and higher exclusion volumes – reduce it by up to 70%. Interestingly, the choice of geometry marginally affects the diffusivity reduction.

### The model captures the heterogeneous and scale-dependent diffusivity reduction induced by cytoplasmic structures

We then sought to compare model predictions for cytoplasmic diffusivity *D*_eff_ against results from osmotic experiments in HP and LP regions.

While experimental and model micro-obstacle volume fractions are equivalent Φ_*m*_ = *ε*_*m*_, model nano-obstacle excluded volume fraction in hyaloplasm *ε*_*n*_ and experimental nano-obstacle excluded fraction in cytoplasm Φ_*n*_ are linked through Φ_*n*_ = (1 − *ε*_*m*_)*ε*_*n*_. Since nano-obstacles are responsible for most of the diffusional hindrance (Fig. 4D, G), we fixed Φ_*m*_ = 0.08 for all cell regions and osmolarities and focused on variations of GFP effective diffusivity with Φ_*n*_ (Fig. 2G).

Fig. 5A-C shows the comparison between model predictions for GFP effective diffusivity *D*_eff_ (Fig. S18) and GFP diffusivity measured by FRAP *D*_exp_. 3D-CFC geometry results from Fig. 4C-G were considered (Fig. S15). For comparison, the full model predictions were systematically compared to results with only the porous hydrodynamic hindrance considered (*T*_*α*_ = *T*_*γ*_ = 1), and with only the tortuosity considered (*H*_*α*_ = 1). For each osmolarity, the model’s cytosolic diffusivity *D*_*α*_ was fitted based on the measured cytoplasmic diffusivity in HP and LP regions. In all cases the full model recovers well the measured diffusivities, with root mean square errors (RMSE) below the ones obtained with only the hydrodynamic or tortuous hindrances.

**Figure 5.**
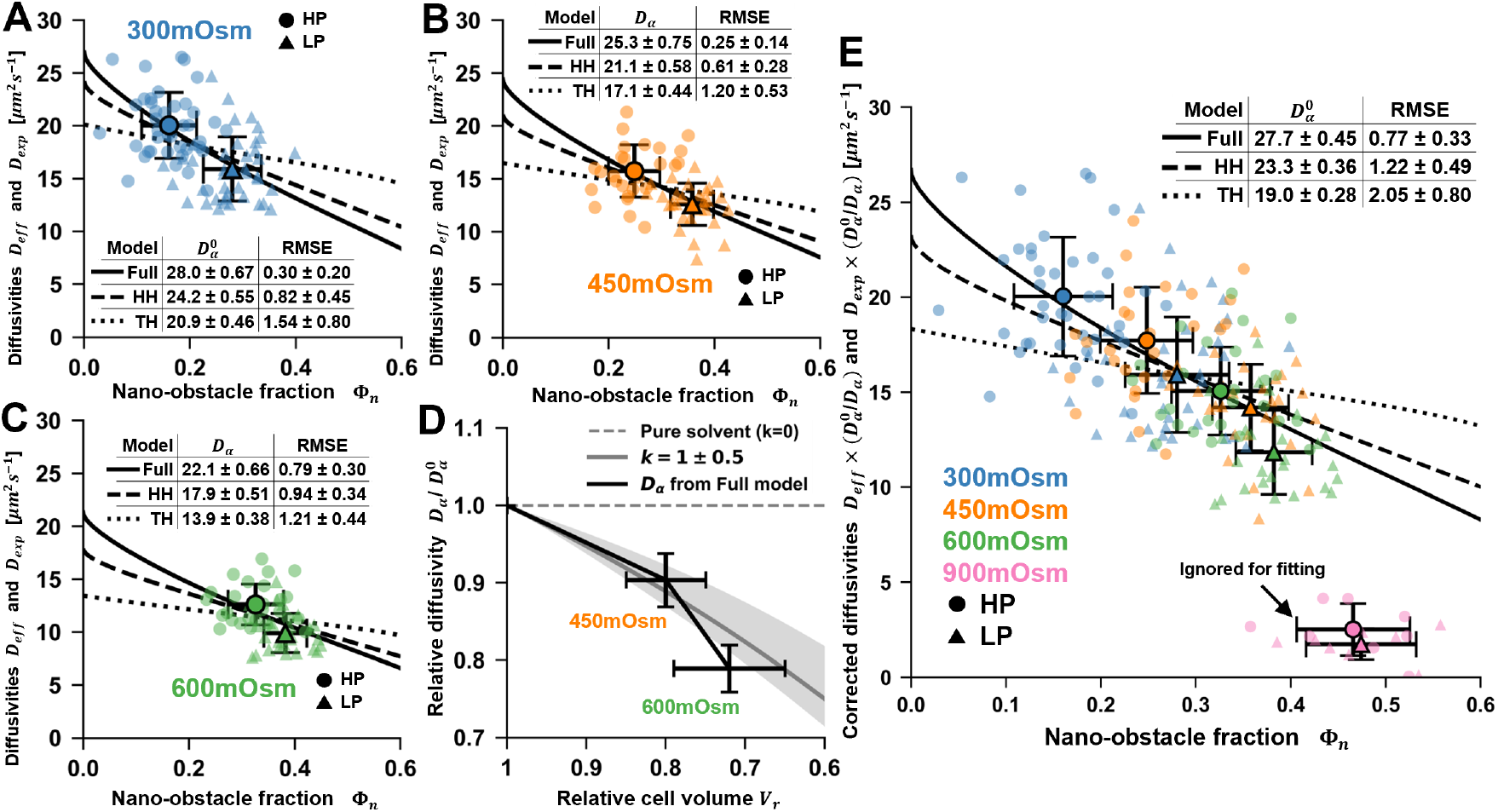
Comparison between the multiscale model and osmotic experiments. Model predictions recovers the 300, 450, 600mOsm results, indicating that cytoplasmic obstacles significantly reduce GFP diffusion through both tortuous and porous hydrodynamic hindrances. (A-C) Model fit with respectively 300mOsm (A), 450mOsm (B), and 600mOsm (C) experimental data. (D) Comparison between full model values for 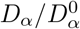 from panels (A-C) and Eq. 5 predictions with *k*^*′*^ = 1 *±* 0.5. (E) Model fit with 300mOsm, 450mOsm, and 600mOsm experimental data altogether. 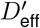 and 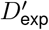 are corrected using Eq. 5 and *k*^*′*^ = 1. 900mOsm data is ignored for the fit, as it likely corresponds to a sudden change in GFP diffusion physics. (A-C, E) HH and TH stand respectively for hydrodynamic hindrance only (with *T*_*γ*_ = *T*_*α*_ = 1) and tortuous hindrance only (with *H*_*α*_ = 1). 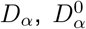, and RMSE are in *µm*^2^*s*^−1^ *±*STD from bootstraping (SI Appendix). Data point with errorbars are experimental average *±*STD. (D) Horizontal errorbar is experimental average *±*STD, vertical errorbar is model fit*±*STD from bootstraping.

Importantly, the *D*_*α*_ values fitted decrease with increasing osmolarity, as reported for the full model in Fig. 5D. Indeed, as water leaks out of the cells in hyper-osmotic conditions, the viscogens concentration – and thus cytosolic viscosity – increases, reducing macromolecular diffusivity. To model this effect, we rewrote Eq. 1 to express the effective diffusivity *D*_eff_ reduction with respect to iso-osmotic cytosolic diffusivity 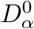

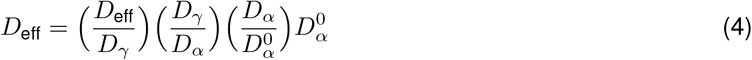

where 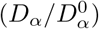 incorporates the influence of osmolarity on the cytosolic diffusivity. Using the ideal solvent model and Stokes-Einstein’s law, cytosolic diffusivity variation with osmotic condition can be expressed as (SI Appendix 4A)

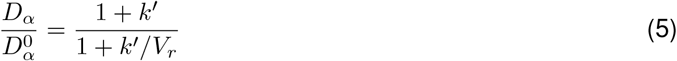

where the constant *k*^*′*^ accounts for cytosol increased viscosity (compared to water) in iso-osmotic condition, and *V*_*r*_ is the cell relative volume after osmotic shock, plotted in Fig. 2D. Based on experimental reports of GFP rotational and short-range translational mobility measurements [30, 22, 58], we estimated *k*^*′*^ ∈ [0.5, 1.5], yielding consistent results with full model predictions for *D*_*α*_ in Fig. 5D.

To compare the model and experimental cytoplasmic diffusivities for all osmotic conditions at once, *D*_eff_ and *D*_exp_ were corrected for cytosolic diffusivity variation with osmotic condition by normalizating them with 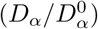 and Eq. 5. As shown in Fig. 5E for *k*^*′*^ = 1, we find that all corrected diffusivity datapoints collapse on a single curve as a function of the nano-obstacle fraction Φ_*n*_ at 300, 450, and 600mOsm. For 900mOsm, a sudden change in diffusion physics is suspected. Importantly, datapoints for HP and LP regions in different osmotic conditions overlap, highlighting the relevance of Φ_*n*_ as a key controlling parameter for cytoplasmic diffusivity.

Moreover, the effective corrected cytoplasmic diffusivity predicted by the full model successfully describes the 300, 450, and 600mOsm corrected data altogether, with lowest RMSE values. As for Fig. 5A-C, this highlights that both tortuous and porous hydrodynamic hindrances are crucial to accurately capture the dependence of effective diffusivity on the nano-obstacle volume fraction.

For completeness, we repeated the analysis with different geometries, values for *k*^*′*^ and for the iso-osmotic nucleoplasm accessible volume fraction (Table S1, S2) and found consistent results. Notably, iso-osmotic cytosolic diffusivity 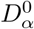 remained between 25 and 40 *µm*^2^*s*^−1^, well above cytoplasmic diffusivity (∼18 *µm*^2^*s*^−1^) and well below the diffusivity in pure water (∼87 *µm*^2^*s*^−1^). This confirms that both the cytoplasmic obstacles and the cytosol viscosity significantly reduce cytoplasmic diffusivity.

Altogether, these results suggest that cytoplasmic structures induce both tortuous and hydrodynamic effects, resulting in a spatially heterogeneous and scale-dependent diffusivity reduction which is stronger in LP regions and at an effective-cytoplasm scale. These findings confirm the relevance of porous media modeling for intracellular diffusion.

### Non-linear particle-size dependence for cytoplasmic diffusivity is captured by the multiscale model

Several experimental reports pointed to a non-linear relation between particle cytoplasmic diffusivity and particle size, departing from Stokes-Einstein’s law [23, 26, 25, 13]. We therefore tested whether the multiscale model could capture this behavior (SI Appendix 4C for detailed methodology).

In the model, the diffusive particle radius *R* determines the nano-obstacles excluded volume fraction and thus the tortuosity *T*_*α*_, and it drives the porous hydrodynamic hindrance *H*_*α*_ (Fig. S19). As both these mechanisms occur at the nano-scale, we neglect the contribution of the micro-obstacles in this section.

To compare the model to literature experimental results, we assumed a homogeneous cytoplasm structure with uniform nano- and micro-obstacle abundances close to our results for MDCK cells. 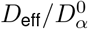 was then estimated for *R <* 10nm using a 3D-CFC ribosome nano-scale geometry (Fig. S20). We observe a strong decrease of cytoplasmic diffusivity with particle radius (up to 75% for *R* = 10nm), which is largely due to the porous hydrodynamic hindrance.

Finally, we compared our model predictions to literature FRAP and FCS diffusivity measurements [23, 26, 25]. As these studies were conducted on different cell lines – with different cytosolic viscosities – we defined the normalized model and experimental cytoplasmic diffusivities 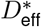 and 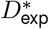 as

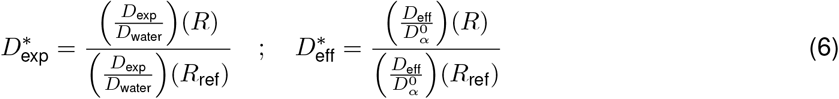

where (*D*_exp_*/D*_water_) is the relative cytoplasmic to pure water diffusivity and *R*_ref_ is an arbitrary particle radius. We set *R*_ref_ = 3.5*nm* so that (*D*_exp_*/D*_water_)(*R*_ref_) could be estimated for the three literature datasets using interpolation.

Fig. 6C compares 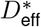 and 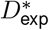 for *R <* 10nm. The full model captures well the non-linear decrease in cytoplasmic diffusivity with increasing particle radius for most particles and cell lines. This confirms that nanometric obstacles in the cytoplasm are responsible for the strong particle-size dependence of diffusivity observed in the literature, and that the porous hydrodynamic hindrance plays a key role inside living cells.

**Figure 6.**
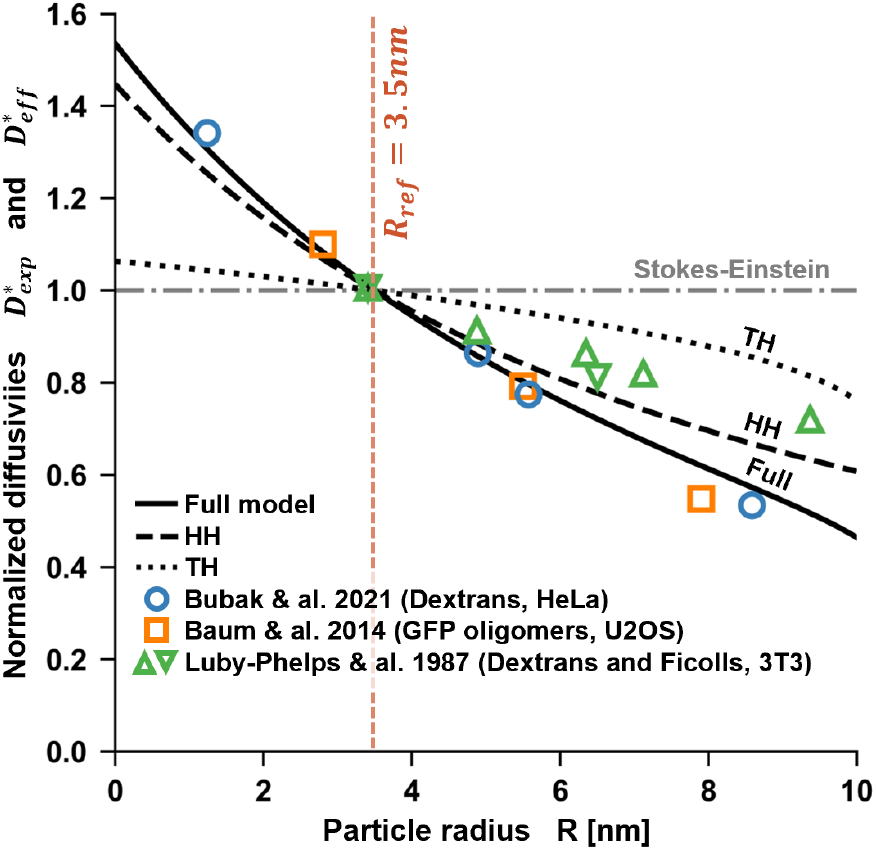
Comparison between model predictions and literature experimental results for different diffusive particle sizes. The model recovers the strong dependence of cytoplasmic diffusivity on particle size observed for several particle types and cell lines. *R*_ref_ = 3.5*nm* is used for normalization in 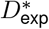 and in 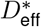. HH and TH stand respectively for hydrodynamic hindrance only (with *T*_*γ*_ = *T*_*α*_ = 1) and tortuous hindrance only (with *H*_*α*_ = 1).

## DISCUSSION

In this work, we proposed an approach based on porous media theory and fluorescence microscopy techniques to study how the cytoplasm structure impacts the diffusion of inert macromolecules. Using confocal microscopy and free-GFP fluorescence to image MDCK cells, we confirmed that the cytoplasm present several regions with significantly different porosities [59, 29, 47]. Further image analysis allowed us to quantify the abundance of structures that act as micrometric or nanometric obstacles. We conducted FRAP and FCS measurements in the identified cytoplasm regions, showing that GFP diffusivity is lower by 20% in regions of lower porosity. To screen a broader porosity range, we conducted experiments in hyper-osmotic conditions, increasing obstacle (and viscogen) concentrations and reducing diffusivity [60, 61, 17]. We find that GFP diffusivity is anti-correlated with the local abundance of obstacles, in particular nano-obstacles.

To identify the physical origin of this anti-correlation, we developed a multiscale model for cytoplasmic diffusion based on a porous media approach. The model, which accounts for cytosolic viscosity – increased by viscogens – and two obstacle-induced diffusional hindrances, recovers our experimental results for GFP diffusion obtained for nano-obstacle excluded volume fractions up to 40%.

While tortuosity that arises from both nano- and micro-obstacles was already shown to reduce diffusion for finite-size particles [36], here we show that porous hydrodynamic hindrance is the dominant porous effect for all obstacle volume fractions and diffusive particle radii investigated, consistently with previous work on bacteria cytoplasm [38]. This further supports the relevance of a framework inspired by porous media and hydrogels to study cytoplasmic diffusion. As only nanometric obstacles induce a porous hydrodynamic hindrance, this highlights the greater importance of nano-obstacles at reducing cytoplasmic diffusivity compared to micro-obstacles. Thus, smaller obstacles — difficult to detect with diffraction-limited microscopy — impede diffusion more effectively than larger, easily observable organelles.

To better identify the nature of the nano-obstacles, we investigated the effects of cytoskeletal filaments depolymerization on GFP diffusion. In the literature contrasted results were obtained, either supporting [37, 23, 28] or tempering the role of F-actin on diffusion [31]. Here, in the considered bulk cytoplasmic regions, we found no clear impact of F-actin and microtubules on GFP diffusion. As studies on hydrogels showed that polymer fibers fraction as low as a few percent could impact macromolecular diffusion [62, 54], we hypothesize that microtubules and F-actin are too scarce in bulk cytoplasm to impact diffusion. Though F-actin may alter diffusion in the cell periphery where it is more abundant, in the regions we studied the most abundant nano-obstacles are likely membranous networks such as endoplasmic reticulum [63] and golgi apparatus, and ribosomes [13].

Importantly, the scale-dependent diffusivity predicted in our model — with a cytosolic diffusivity 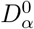 higher than the effective cytoplasmic diffusivity *D*_eff_ – is in line with previous studies [30, 58, 22, 64]. Thus our work confirms that the transient-subdiffusive behavior reported previously is notably due to the presence of structures in the cytoplasm [30, 65], and in particular of nanometric obstacles.

Moreover, the model was able – without further adjustments – to recover the non-linear dependence of diffusivity on particle-size observed previously, for several particles types and in several cell lines [13, 23, 26, 25]. While previous studies accounted for particle-size diffusivity dependence using phenomenological models [26, 13], here we captured this behavior by deriving a mechanistic model based on transport theory in porous media and in hydrogels. This model shows that deviation from Stokes-Einstein’s law for inert particles in the cytoplasm is primarily driven by the combination of tortuosity and porous hydrodynamic hindrance caused by nanometric obstacles. Interestingly, a similar relation between globular protein size and diffusivity was observed in the nucleoplasm [66, 26, 23], suggesting that our approach might be suitable to investigate passive macromolecule transport in the nucleoplasm.

Altogether, these findings support the relevance of porous media modeling approaches to better understand cytoplasmic diffusion of inert particles in eukaryotic cells. The cytoplasmic structures, in particular nanometric obstacles such as ribosomes and membranous networks, hinder globular protein diffusion through tortuous and porous hydrodynamic hindrances, resulting in a spatially heterogeneous, scale-dependent, and particle-size dependent diffusivity reduction. Importantly, our approach suggests the possibility of a universal link between the abundance of nanometric obstacles and diffusion properties in the cytoplasm, regardless of the cell line, cytoplasmic region, or osmotic condition.

This work provides a multiscale physical framework to improve our understanding of dynamic intracellular processes for which space- and size-dependent diffusion is central, such as caspase signaling cascade during apoptosis and actin treadmilling during cell migration.

## Data availability

All data are available in the manuscript and in the supplementary materials.

## Acknowledgment

We thank C. Leduc, M. Delarue, H. Duval and H. Auradou for insightful discussions. We thank CAM lab members for introduction to cell culture techniques.

