## Supplementary Information for "Cytoplasmic crowding acts as a porous medium reducing macromolecule diffusion"

**Supporting Information for**
### Cytoplasmic crowding acts as a porous medium reducing macromolecule diffusion.

Olivier Destrian, Nicolas Moisan, René-Marc Mège, Benoît Ladoux, Benoît Goyeau, Morgan Chabanon

Corresponding author: Morgan Chabanon


**This PDF file includes:**

SI Appendix 1 to 4
Figures S1 to S20, cited in the main manuscript
Figures S21 to S28, cited only in SI Appendices
Tables S1 to S2
Legend for Movie S1
SI References

**Other supporting materials for this manuscript include the following:**

Movie S1

#### Table of Contents

1. **Supplementary experimental material and methods** **3**

a. *MDCK cell culture* *3*

b. *Confocal microscopy setup* *3*

c. *Hyper-osmotic shocks and cytoskeletal depolymerizing drug treatments* *3*

d. *Fixation and immunostaining procedures* *4*

e. *Inference of micro- and nano-obstacle fractions* $\Phi m$ *and* $\Phi n$ *from confocal microscopy data* *4*

f. *Computation of cell volume from confocal microscopy data* *6*

g. *FRAP protocol* *6*

h. *FCS protocol* *7*

2. **Investigation of the role of cytoskeletal filaments using depolymerizing drugs** **9**

3. **Multi-scale model derivation and resolution** **10**

a. *Introduction to the method of volume averaging* *10*

b. *Nano-scale hypotheses justification* *10*

c. *Geometrical and physical analysis of GFP diffusion in the nano-scale medium* *11*

d. *Model derivation: estimation of nano-obstacles induced diffusional hindrances* *13*

e. *Micro-scale physical analysis and model derivation: estimation of micro-obstacle induced tortuous hindrance* *15*

f. *Numerical resolution of the closure problems* *16*

4. **Supplementary developments for model comparison with experiments** **19**

a. *Cytosolic diffusivity dependence on osmolarity and correction* *19*

b. *Bootstrapping for assessment of uncertainty in model-experiment fitting procedures* *21*

c. *Model application to particle with a radius* $R < 10nm$ *21*

5. **Supporting Figures** **24**

**SI References** **56**

### 1. Supplementary experimental material and methods

#### a. MDCK cell culture

MDCK type II cells (ATCC CRL-2936) were cultured in Dulbecco's modified eagle medium (DMEM) with red-phenol, GlutaMax, 4.5g/L D-Glucose, Pyruvate (Gibco Ref. 31966-021), 10% fetal bovine serum (Biowest, S1810-500), and 1% penicillin-streptomycin (Gibco Ref. 15140-122). Cells were incubated at 37°C, 5% CO2 and passaged two times a week using 0.05% Trypsin-EDTA (Gibco, Ref. 25300-054). Cells were tested negative for mycoplasma (MycoSPY Master Mix, Biontex). Cells were transduced to express free-GFP using MISSION® TRC3 ORF GFP Lentivirus Control (Sigma) and were then sorted using FACS for their GFP expression levels. Sorted cells with high fluorescence level were used for fluorescence Z-stacks images and FRAP measurements. Sorted cells with low fluorescence level were used for FCS measurements.

MDCK live-cell samples were prepared from 48h to 72h before experiments. Cells were seeded in fluorodishes (FD35-100, World Precision Instruments) coated with fibronectin (Sigma Aldrich, 1003624379). On the day of experiments, cells were washed with PBS 1X and then immersed in DMEM without red-phenol, with 4.5g/L D-Glucose, L-Glutamine, 25mM HEPES, sodium pyruvate (Gibco Ref. 21063-029), 10% fetal bovine serum (Biowest, S1810-500), and 1% penicillin-streptomycin.

On the day of experiments, cells confluency was typically between 10% and 30% so that cells formed small islands (Fig. S1). Large and spread cells present at the border of the islands were used for experiments. The same samples were used for FRAP measurements and confocal Z-stacks, and in a given sample, each cell used for acquiring Z-stack images was also used for acquisition of FRAP data. FCS experiments were conducted on separate samples.

#### b. Confocal microscopy setup

All optical microscopy experiments were performed using a Zeiss LSM980 – Airyscan2 microscope with incubated enclosure at 37°C, 5%CO2, with ZenBlue software. For Z-stacks and FRAP measurements, a Plan Apochromat 63X/1.4-NA oil DIC objective was used. For FCS measurements, a C-APO 40X/1.2-NA Water Corr UV-VIS-IR objective was used, with PicoQuant FCS upgrade kit for LSM (integrating two PMA cooled hybrid photomultipliers plugged on a MultiHarp 150 Multichannel time correlated single photon counter, and laser source at 480 nm generating nanosecond pulses at a repetition rate of 40MHz) along with Symphotime 64 software.

Z-stacks were acquired using Airyscan multiplex SR4Y mode, and raw Z-stacks data was processed using ZenBlue 2D Airyscan post-processing. The Airscan detector was calibrated each day prior to the experiments. FRAP recovery images were acquired using confocal mode with 1UA pinhole aperture. FCS data was acquired with 1 UA pinhole aperture.

#### c. Hyper-osmotic shocks and cytoskeletal depolymerizing drug treatments

Hyper-osmotic shocks and cytoskeletal depolymerizing drugs experiments were carried out by quickly replacing the medium on the fluorodish in the incubated enclosure of the microscope with the same medium containing sucrose or drugs, with great care applied to avoid moving the sample. The medium was prepared before experiments and heated at 37°C just before the experiments. Hyper-osmolar media (450, 600, 900mOsm) were prepared by adding sucrose (Sigma-Aldrich, 102500489) at respective concentrations of 50g/L, 100g/L, and 200g/L. Drugs containing media were prepared by adding Latrunculin-A (Sigma, L5163), Cytochalasin-D (Calbiochem, 250255), Nocodazole (Thermofischer, 358240100) dissolved in DMSO at the final concentrations indicated for each experiment. For osmotic experiments, hyper-osmotic sample were used between 10mn and 60mn after medium replacement, and drug treated samples were used between 30 and 90mn after medium replacement.

##### d. Fixation and immunostaining procedures

Cells were seeded on coverslips in a 12-well plate with culture medium between 48h to 72h before fixation and kept at 37°C 5%CO2. On the day of fixation, cells were fixed with PBS+4% formaldehyde (Thermo Scientific, Ref 28908) for 10mn. Then, cells were rinsed several times using PBS. Cells were permeabilized using Triton 0.5% (Sigma-Aldrich, T8787) for 10mn, and were rinsed again using PBS. Cells were incubated in PBS+BSA 1% (Roche, Ref 10 735 094 001) for 1h, alpha-tubulin primary rat antibody (Bio-Rad, MCA77G) was added overnight. The next day, cells were washed using PBS, and the secondary antibody (anti-rat Alexa647, Life Technologies, A21247) was added along with Phalloidin-Alexa568 (Life Technologies, A12380) for 2h. Cells were rinsed again and immersed in PBS containing Hoechst 33342 (Life Technologies, H3570). Cells were rinsed again and the coverslips were mounted on a coverslide with Mowiol (Sigma-Aldrich, 101904315).

##### e. Inference of micro- and nano-obstacle fractions $\Phi_m$ and $\Phi_n$ from confocal microscopy data

Free-GFP fluorescence Z-stack data was first processed using a custom Python code to estimate average GFP fluorescence in the low porosity (LP) and high porosity (HP) regions of each cell. To estimate the fluorescence of the different intracellular regions, the HP, LP, and nucleoplasmic regions were manually tagged using a basic image analysis software such as Paint on a copy of the Z-stack file. The original and tagged Z-stack files were then loaded in python. The absolute fluorescence of the HP, LP, and nucleoplasmic tagged regions were computed from the original Z-stack file. The tagged nucleoplasm region never comprised the nucleoli nor the nuclear borders. For each cell, the nucleoplasm fluorescence value was used for HP and LP fluorescence normalization.

Assuming the effective porosity (in accessible volume fraction) for GFP is proportional to GFP fluorescence, the *relative* porosity of the HP and LP regions is known from normalized fluorescence data. To estimate the *absolute* effective porosity in HP and LP regions $\varepsilon = 1 - \Phi_n - \Phi_m$ , we needed to estimate to what effective porosity $\varepsilon_{nucleus}$ the nucleoplasm fluorescence corresponds. We assumed GFP concentration in the cytosol $C_\alpha$ to be uniform in the cell. Since GFP can move in and out of the nucleus on the timescale of ≈ 1−5mn (Fig. S2, and consistently with [1]), we also assumed that GFP concentration in the nucleosol is equal to $C_\alpha$. In a given intracellular region, we thus have in the cytosolic $\alpha$-phase

$$\langle C_\alpha \rangle = \varepsilon C_\alpha$$

Where $\langle C_\alpha \rangle$ is the superficial average of free-GFP cytosolic concentration in a given cytoplasmic region, and is defined as

$$\langle C_\alpha \rangle = \frac{1}{V} \int_{V_\alpha} C_\alpha \ \ dV$$

Where $V_\alpha$ is the cytosolic/nucleosolic domain in a HP, LP or nucleoplasmic region. The superficial average of free-GFP concentration in the HP/LP/nucleus regions can thus be written as

$$\langle C_\alpha \rangle_{HP} = \varepsilon_{HP} \times C_\alpha$$
$$\langle C_\alpha \rangle_{LP} = \varepsilon_{LP} \times C_\alpha$$
$$\langle C_\alpha \rangle_{nucleus} = \varepsilon_{nucleus} \times C_\alpha$$

In a given cell, the GFP fluorescence intensity $I$ in a region (either HP, LP or nucleoplasmic) is assumed proportional to the region superficial free-GFP concentration, thus we have

$$\frac{I_{HP}}{I_{nucleus}} = \frac{\varepsilon_{HP}}{\varepsilon_{nucleus}}$$
$$\frac{I_{LP}}{I_{nucleus}} = \frac{\varepsilon_{LP}}{\varepsilon_{nucleus}}$$

$I_{HP}/I_{nucleus}$ and $I_{LP}/I_{nucleus}$ are known from the results presented in Fig. 1E, but we still need to estimate $\epsilon_{nucleus}$ so that the absolutes effective porosities for HP and LP regions can be estimated using

$$\varepsilon_{HP} = \varepsilon_{nucleus} \times \frac{I_{HP}}{I_{nucleus}}$$
$$\varepsilon_{LP} = \varepsilon_{nucleus} \times \frac{I_{LP}}{I_{nucleus}}$$

It is thus crucial to find an estimate for $\varepsilon_{nucleus}$. Consistently with [2], we observed that the nucleoplasm fluorescence (outside of nucleoli) is higher and more homogeneous than the cytoplasm fluorescence, so the nucleoplasm region selected has a limited impact on $I_{nucleus}$. We assumed the nucleoplasm structures to be primarily composed of chromatin, which act as a nano-obstacle. Thus $\varepsilon_{nucleus}$ can be obtained by estimating the nucleoplasmic excluded fraction for free-GFP by chromatin

$$\varepsilon_{nucleus} \approx 1 - \Phi_{chromatin}$$

We suppose that the high nucleoplasm fluorescence corresponds to euchromatin domains and that heterochromatin domains are mainly located in the perinucleolar region and near the nuclear membrane [3]. We thus focus on euchromatin domains. In iso-osmotic condition, it is reasonable to assume that the nucleoplasmic chromatin occupied fraction in these domains is $\Phi^{\text{occupied}}{}_{chromatin} \approx 0.1 \pm 0.05$ [4]. Assuming that the volumes excluded by each chromatin fiber do not intersect (as for nano-obstacles, see SI Appendix 3.c)

$$\frac{\Phi_{chromatin}}{\Phi^{\text{occupied}}{}_{chromatin}} = \left(\frac{R_{GFP} + R_{chromatin}}{R_{chromatin}}\right)^2 = 2.13$$

Which implies that $\varepsilon_{nucleus} \in [0.7, 0.9]$ in iso-osmotic condition. All experimental data in iso-osmotic conditions presented in the main text (Figs. 1, 2, 5 and 6) use $\varepsilon_{nucleus} = 0.85$.

With hyper-osmotic shocks, the nucleus is expected to shrink, thus we needed to estimate $\varepsilon_{nucleus}$ dependence on the nucleus volume to have a porosity reference for each experimentally tested osmotic condition. Following a hyper-osmotic shock, both the cytoplasm and the nucleoplasm shrink. Thus, both the cytoplasmic and nucleoplasmic porosities are reduced. Let us focus on a cell submitted to an arbitrary osmotic shock. We define $V_{nucleus}$, $V_{chromatin}$, $V_{nucleosol}$ as respectively the nucleus volume, the nucleoplasmic volume excluded for free-GFP by chromatin, and the nucleosolic fluid volume accessible for free-GFP. For any variable $A$, we note $A_0$ its value in iso-osmotic condition, before the hyper-osmotic shock. We have before and after the hyper-osmotic shock

$$V^0_{\text{nucleus}} = V^0_{\text{chromatin}} + V^0_{\text{nucleosol}}$$
$$V_{\text{nucleus}} = V_{\text{chromatin}} + V_{\text{nucleosol}}$$

We assume the volume occupied by chromatin is not changed by the osmotic shock. With the no-intersection hypothesis for the volume excluded by each fiber, we have $V_{chromatin} = V^0_{chromatin}$. From the definition of the effective porosity of the nucleus $\epsilon_{nucleus} = V_{nucleosol}/V_{nucleus}$ we obtained

$$\epsilon_{\text{nucleus}} = \frac{V_{\text{nucleus}} - V_{\text{chromatin}}}{V_{\text{nucleus}}} = 1 - \frac{V^0_{\text{chromatin}}}{V^0_{\text{nucleus}}} \times \frac{V^0_{\text{nucleus}}}{V_{\text{nucleus}}}$$

Consistently with the literature [5], [6], [7], we assumed that the nucleus volume is approximately impacted in the same proportion as the cell volume after a hyper-osmotic shock, and thus

$$\frac{V_{\text{nucleus}}}{V^0_{\text{nucleus}}} \approx V_r$$

Where $V_r$ is the cell volume ratio after/before the hyper-osmotic shock. This quantity was estimated experimentally in Fig. 2D. We finally obtained a relation for the dependence of the porosity reference on the osmotic condition

$$\epsilon_{\text{nucleus}} = 1 - \left(1 - \epsilon^0_{\text{nucleus}}\right) \times V_r^{-1}$$

The results for $\epsilon_{\text{nucleus}}$ for different $V^r$ and $\epsilon^0_{\text{nucleus}}$ values are shown in Fig. S6. At this point, we could estimate $\varepsilon = 1 - \Phi_n - \Phi_m$ for HP and LP regions and for all osmotic conditions. To estimate $\Phi_n$ and $\Phi_m$, we needed to distinguish these obstacles in our Z-stack data. We used a custom segmentation procedure (see github for code) to process the Z-stack data as presented in Fig. 1H. Using thresholding and morphological operations, the program returns a mask $M_m$ that indicates which pixels of a selected region belong to the micro-obstacle phase. The micro-obstacle fraction in a HP or LP region (delimited by the $M_{region}$ mask) was then estimated by

$$\Phi_m = \frac{M_m}{M_{\text{region}}}$$

In other words, the micro-obstacle fraction was simply estimated as the proportion of the selected region that has been identified as part of the micro-obstacle phase. The nano-obstacle fraction in the selected region was then deduced using the relation $\Phi_n = 1 - \epsilon - \Phi_m$.

##### f. Computation of cell volume from confocal microscopy data

To estimate the absolute volume before and after osmotic shock, we used the Z-stacks taken before and after the shock for each of the cells. The two Z-stacks were manually processed using basic image processing software to remove the other cells. A fluorescence threshold relative to the nucleoplasmic region fluorescence was then applied to convert the Z-stack data into a binary image (Fig. S4). After a few morphological operations (see github for code), the 1-valued pixels (high fluorescence) were then summed and converted into an absolute volume. The relative cell volume $V_r$ was obtained by dividing the absolute volume after shock by the volume before shock.

##### g. FRAP protocol

During the pre-bleach and post-bleach imaging phases, the confocal 2D scanning mode of the microscope was used. Pinhole aperture was set to 1 A.U. and the 488nm laser power was set to a low value, 1%. A 25µm × 1µm rectangle was imaged continuously at a frequency of 50Hz. The point of using a thin 1µm rectangle was to increase the imaging speed because free-GFP diffuses quickly and the FRAP recovery is generally fast ≤ 500ms. The rectangle is highly elongated with a length of 25µm, to observe the full recovery spatial profile and for the rectangle borders to reach cytoplasmic regions that are nearly unaffected by the bleach.

During the bleaching phase, the confocal 0D spot mode of the microscope was used. The 488nm laser power was set to 100% (with a measured 5.2 mW laser power upstream of the microscope objective) with a point bleach duration of 25ms. The reason for bleaching a single spot was that it allowed for fast and sufficiently deep bleaching, which was essential for the quality of the FRAP measurements. The size of the bleached region was not imposed and was physically controlled by the bleaching point spread function of the microscope and by the diffusion of free-GFP during the bleaching phase. The bleached radius measured on the first post-bleach images was typically near ≈ 3 − 6μm. For each cell, the FRAP experiment was repeated three times in the HP and in the LP regions. In each region, a ≈ 20s delay was observed between each repetition to ensure a complete fluorescence recovery. The time between the last pre-bleach image and the first post-bleach image was close to 50ms and was slightly variable between experiments.

From the recovery phase images, we observed that the bleaching profile was gaussian at all times and that the fluorescence would fully recover everywhere in the imaged rectangle, meaning no significant GFP bound fraction was present. Thus, we used a gaussian diffusion model to infer GFP diffusivity from our FRAP experiments

$$\forall t' \geq 0, \frac{I(x,t')}{I_0} = 1 - \frac{K'}{4D_{GFP}\left(t' + t_{pb}\right)} \exp^{-\frac{x^2}{4D_{GFP}\left(t'+t_{pb}\right)}}$$

Where $I(x,t')$ is a 1D recovery fluorescence profile, $I_0(x)$ is the pre-bleach fluorescence profile, $K'$ is a constant proportional to the bleaching depth, $t_{pb}$ is a time-constant, and $D_{GFP}$ is GFP diffusivity. $(K', t_{pb}, D_{GFP})$ were fitted using data from the experimental chronographs, as presented in Fig. 3A. To obtain these experimental chronographs from raw confocal images, several processing steps were performed.

First, the data from each repetition of the FRAP experiment in a given region were individually processed. The rectangle lines of the 2D images were averaged into a 1D fluorescence profile. Then, the background noise was estimated from the average signal in a region outside of the cell and subtracted to the 1D profile. To correct for involuntary photobleaching during imaging phases, we applied a reference area correction (thought to be more reliable according to [8]) using the fluorescence of the extremities (2μm each side) of our imaged rectangle as presented in Fig. S21. To account for the spatially heterogeneous fluorescence inside the cytoplasm, and consistently with other works [9], [10], [11], we also normalized fluorescence signal by the pre-bleach fluorescence profile $I_0$. Thus, the experimental chronograph for a single FRAP repetition was obtained. Then, the three repetition chronographs were averaged to obtain an average experimental chronograph (for each region of each cell) which was used for fitting with the above equation. The fitting was applied to the chronograph data taken from 100ms to 600ms after photobleaching.

##### h. FCS protocol

FCS was used to measure GFP relative diffusivity in HP and LP regions of MDCK live cells. During FCS experiments, GFP fluorescence data was acquired five times for five minutes in each HP and LP regions, to account for cell movement during acquisition. Autocorrelation curves were computed for each individual acquisition using Symphotime software. The autocorrelation curves were then imported into a custom python program. An average autocorrelation curve was computed for each region for each cell, and then used to fit a FCS diffusion model. A simple FCS diffusion model was considered [12]

$$G(t) = \left(\rho\left[1 + \frac{t}{\tau_D}\right]^{-1}\left[1 + \frac{t}{\tau_D\kappa^2}\right]^{-\frac{1}{2}}\right) + \rho_\infty$$

Where $\rho$ is a multiplicative constant, $t$ is the time delay, $\tau_D$ is the characteristic residence time, and $\kappa$ is the axial/radial length ratio of the measurement volume. In this equation we added the constant $\rho_\infty$ to correct the autocorrelation offset at very large > 1000ms timescales, which was due to organelle movements in and out of the measurement volume. This equation was used for fitting procedures (with $\rho$, $\rho_\infty$ and $\tau_D$ as fitting parameters) with our experimental autocorrelation curves. This provided simple and satisfying data fitting, with an example present in Fig. S22. As the same microscope configuration was used for all FCS measurements, we finally observed that

$$D_{GFP} \propto \tau_D^{-1}$$

Which allowed us to compare relative diffusivity of GFP in HP and LP regions and in each cell without relying on the precise characterization of the FCS measurement volume that would be required for estimating absolute values for GFP diffusivity $D_{GFP}$. This way the FCS results were sufficient to confirm the difference of GFP diffusivity observed between LP and HP regions.

For plotting purposes (in Fig. 3B and S22), the experimental correlation data points as well as the fitted autocorrelation curves $G_{model}(t)$ were normalized as follows

$$G_{normalized}(t) = \frac{G(t) - G(+\infty)}{G(0) - G(+\infty)}$$

#### 2. Investigation of the role of cytoskeletal filaments using depolymerizing drugs

To gain insight on the nature of the nano-obstacles responsible for the cytoplasmic diffusional reduction, we tested the role of F-actin and microtubules fibers, two of the main cytoskeletal filaments. MDCK cells were treated with Latruculin-A and cytochalasin-D to disrupt F-actin, and with nocodazole to depolymerize microtubules. The effect of these drugs on the intracellular organization was examined on immunostained fixed cells (Fig. S7) confirming the effect of these drugs on the cytoskeleton. These drugs were also tested on live cells, that were imaged using confocal Z-stack microscopy of free-GFP fluorescence (Figs. S8-11).

Though all three drugs drastically affected cell morphology, the HP and LP regions could still be identified for limited drug concentrations, with a visible difference in GFP signal and in obstacles fractions between the two (Fig. S12). FRAP diffusivity measurements were conducted for GFP in HP and LP regions (Fig. S13). FRAP results showed that microtubule depolymerization do not significantly affect free-GFP diffusivity, while the disruption of F-actin results in a mild reduction of GFP diffusivity in HP and LP regions. These results suggest that F-actin and microtubules do not significantly hinder diffusion in the studied bulk cytoplasm regions (HP and LP regions). An explanation could be that microtubules are not abundant enough and that F-actin is mostly located in the cell cortex and lamellipodia, regions that were not studied here. Also, the effect of these cytoskeleton depolymerizing drugs may be more complex than previously though. Thus, as depolymerizing drugs induce a global morphological disruption of the cells, the key role of F-actin and microtubules on cell architecture is indeed confirmed, yet their direct effect on diffusion of mid-size inert macromolecules as free-GFP remains unsure.

#### 3. Multi-scale model derivation and resolution

##### a. Introduction to the method of volume averaging

The method of volume averaging [13] was used several times in the derivation of our multiscale model. The interested reader is directed to [13] (Chapter 1) for a detailed upscaling of heterogeneous reaction-diffusion in a homogeneous porous medium. Here, a very brief introduction to the homogenization procedure is presented and illustrated in Fig. S23.

The Method of Volume Averaging is an upscaling technique applicable to porous media in which the local transport equations, that are valid at the pore-scale (e.g., the solutal diffusion equation in the fluid phase) are known. Considering a medium constituted of a $\omega$ fluid-phase and a $\kappa$ solid-phase, upscaling procedure allows for the rigorous derivation of macroscopic transport equations (e.g., a macroscopic diffusion equation) along with effective transport parameters (e.g., an effective diffusivity). These macroscopic transport equations are valid at a macro-scale $L$ much larger than the averaging volume of length $l$, itself larger than the typical pore size $l_\omega$ and typical obstacle size $l_\kappa$. At the macro-scale, the porous medium is described as a continuous equivalent medium: it is a homogenized representation of the porous medium. The transition from the two-phase pore-scale medium to the homogenized medium is achieved by taking into account the effects of the pore-scale boundary conditions at the interface $A_{\omega\kappa}$ in a macroscopic continuous transport equation. For a porous medium constituted of a $\kappa$ solid-phase and a $\omega$ fluid-phase, the general procedure can be summarized as follows:

1. Definition of the pore scale transport equations: As an input of the method, the transport equations at the pore-scale and the boundary conditions at the fluid-solid interface need to be known.

2. Derivation of the unclosed spatially averaged equations: The pore-scale equations are averaged over a representative elementary volume significantly larger than the pore size, using the averaging theorems. After introducing Gray's decomposition, each remaining local variable field is separated into its spatial averaged value and its local spatial deviation. Spatially averaged transport equations featuring both the spatial deviation and spatially averaged variables are obtained.

3. Determination and resolution of the closure problems: The spatially averaged equation is subtracted from the local pore-scale equation. With additional assumptions, a boundary-value problem for the spatial deviation variable is obtained. Using a linear decomposition of the spatial deviation variable into closure variables, the closure problems are obtained. These closure problems do not depend on the spatially averaged variables and can be solved numerically.

4. Derivation of the closed macroscopic transport equations: The closure variables are injected into the unclosed spatially averaged transport equations, leading to the obtention of the effective transport equations. The macroscopic transport equations feature effective transport parameters that can be estimated from the solution of the closure problems.

##### b. Nano-scale hypotheses justification

Prior to quantify the effects of nano-obstacles on GFP diffusion, we analyzed them quantitatively. Many structures can be considered as nano-obstacles, and to simplify our approach, we assumed that most nano-obstacles were geometrically similar to F-actin filaments or ribosomes. F-actin structures are among the thinnest nano-obstacles present ($R_{act} = 4nm$), while ribosomes are likely closer to the average size of nano-obstacles ($R_{rib} = 12.5nm$). Studying these two configurations allowed us to study the dependence of our hypothesis and model results to the geometry of nano-obstacles. We posed several important hypotheses:

1. Free-GFP is a solute in the cytosolic fluid: With a radius $R_{GFP} = 2.3nm$, GFP can be considered a diffusive solute in the cytosolic fluid that is mostly composed of water $R_{H_2O} = 0.15nm$.

2. The nano-obstacles are immobile: The nano-obstacle velocity is expected to be negligible compared to free-GFP velocity. To justify this assumption, we used dimensional analysis. It is possible to roughly estimate a characteristic time $\tau_D$ for GFP diffusion around a F-actin fiber

$$\tau_D \approx \frac{R_{act}^2}{D_{GFP}} \approx \frac{4nm}{20 - 80\mu m^2 s^{-1}} \leq 0.2\mu s$$

Where $D_{GFP}$ is a very rough estimate of GFP diffusivity in pure cytosol. Thus, free-GFP movement around fibers is very fast compared to the time range for F-actin turnover [14]. Unattached ribosomes likely have a higher mobility than cytoskeletal fibers. However, since ribosome diffusivity is way lower than GFP diffusivity, the assumption that ribosomes are immobile on the timescales of GFP movement around them is also reasonable

$$D_{\mathrm{rib}} \approx 0.1 - 1\mu m^2 s^{-1} \ll D_{GFP} \approx 20 - 80\mu m^2 s^{-1}$$

Where $D_{\mathrm{rib}}$ is a rough estimate for ribosome diffusivity [15]. More generally, other structures with a characteristic size larger than free-GFP, such as membranous networks, are expected to move slowly compared to free-GFP. These structures were therefore modeled as nano-obstacles. On the other hand, compounds that are smaller than free-GFP and that are not bound were modeled as solutes in the cytosolic fluid phase and acted as viscogens [16].

3. The nano-obstacles are impermeable: While free-GFP cannot penetrate stable macromolecular assemblies such as ribosomes and F-actin fibers, it may permeate through certain membranous structures. However, in practice the timescales involved are expected to be very large compared to the characteristic time $\tau_D$ for GFP diffusion around them. The nano-obstacles are also assumed to be impermeable for solvent particles.

##### c. Geometrical and physical analysis of GFP diffusion in the nano-scale medium

With the aforementioned hypotheses, to highlight the specificities of free-GFP mobility at the nano-scale, we characterized the nano-scale medium geometrically. From the hypothesis presented in the last paragraph, we have that the radius of many nano-obstacles is typically in the range of 4 − 12.5nm. This implies that the free-GFP radius $R_{GFP} = 2.3nm$ is of the same order of magnitude as the nano-obstacle size. This has important implications since we relied on the framework of continuum mechanics that model solute particles as point-like. For infinitely small particles, the whole cytosolic volume is accessible for the particles' center of mass. For small but finite particles, such as water molecules $R_{H_2O} = 0.15nm$ << $R_{act} = 4nm$, this assertion is still reasonable. However, for larger particles such as GFP, the cytosolic volume accessible for the GFP center of mass is significantly smaller than the whole cytosolic volume, as illustrated in Fig. 1J.

We defined the cytosolic physical volume fraction $\epsilon_\alpha^* \in [0,1]$ as the volume fraction the cytosol occupies in the nano-scale porous medium. The nano-obstacle occupied volume fraction is its complementary with $\epsilon_n^* = 1 - \epsilon_\alpha^* \in [0,1]$. This description is adapted to characterize the nano-scale structure and to study the transport of small particles such as water molecules.

Alternatively, we defined the cytosolic accessible volume fraction (for free-GFP mass-center) $\epsilon_\alpha \in [0,1]$. The nano-obstacle excluded fraction (for free-GFP mass-center) is its complementary with $\epsilon_n = 1 - \epsilon_\alpha \in [0,1]$. We have $\epsilon_\alpha \leq \epsilon_\alpha^*$ and $\epsilon_n^* \leq \epsilon_n$. These two parameters $\epsilon_\alpha$ and $\epsilon_n$ refer to the effective porous medium perceived by the free-GFP particles considered as point-like. This description is adapted to characterize the nano-scale structure effect on free-GFP or other spherical particles with the same radius as free-GFP.

The two sets of variables $(\epsilon_\alpha, \epsilon_n)$ and $(\epsilon_\alpha^*, \epsilon_n^*)$ are both important since $(\epsilon_\alpha, \epsilon_n)$ is relevant to study the transport of GFP particles and $(\epsilon_\alpha^*, \epsilon_n^*)$ is relevant to study the transport of water and small solute particles, impacting the porous hydrodynamic hindrance. Establishing a relation between $\epsilon_n$ and $\epsilon_n^*$ is thus necessary.

We proposed a very simple approach to obtain such a relation for F-actin-like nano-obstacles or ribosome-like nano-obstacles. In a porous medium composed of several nano-obstacles immersed in cytosolic fluid, we assumed no intersections between the volume excluded by each obstacle. Then we have for infinitely long cylindrical F-actin fibers and for spherical ribosomes

$$\left(\frac{\epsilon_n}{\epsilon_n^*}\right)_{act} = \left(\frac{R_{GFP} + R_{act}}{R_{act}}\right)^2 = 2.48$$
$$\left(\frac{\epsilon_n}{\epsilon_n^*}\right)_{rib} = \left(\frac{R_{GFP} + R_{rib}}{R_{rib}}\right)^3 = 1.66$$

For evenly spaced obstacles, we expect the no-intersection hypothesis to be reasonable. For randomly arranged obstacles, this hypothesis is less obvious. To assess this assumption, we compared this model with the Swiss-cheese model [17]

$$\frac{\epsilon_n}{\epsilon_n^*} = \frac{1 - (1 - \epsilon_n)^{r^*}}{\epsilon_n}$$

Where $r^*$ is the $\frac{\epsilon_n}{\epsilon_n^*}$ ratio obtained when the excluded volumes of each obstacle do not intersect. The results are presented in Fig. S24 and indicated that the no-intersection hypothesis is satisfying for $\epsilon_n \leq 0.5$, in accordance with our experimental results for $\Phi_n = \epsilon_n(1 - \Phi_m)$ presented in Fig. 2F.

We then highlighted the specificities of GFP diffusion at the nano-scale by studying the nano-scale pore size. In the literature, the cytoplasmic pore size (taking all types of structures into account) was estimated to be close to 10 − 20nm [18].

Here, we estimated an approximative nano-scale pore size using geometrical considerations and the estimate $\epsilon_n \in [0.1, 0.5]$. We considered two simplified representative geometries (as used in REVs, see Fig. 4C): a centered disk (with $R = R_{act} = 4nm$) in a periodic 2D cytosolic domain mimicking regularly spaced F-actin fibers (2D-CD F-actin), and a cubic face-centered arrangement of spheres (with $R = R_{rib} = 12.5nm$) in a periodic 3D cytosolic domain mimicking a ribosome population (3D-CFC ribosomes).

We defined the pore size $L_p$ as the size of the cytosolic domain bottleneck for free-GFP movement. For biologically relevant nano-obstacle fractions $\epsilon_n \in [0.1, 0.5]$, with the 2D-CD F-actin geometry, we have $L_p \in [10, 25]nm$ and for the 3D-CFC ribosome geometry we have $L_p \in [15, 40]nm$. This means that depending on the nano-obstacle abundance level, we expect $L_p$ in the range of tens of nanometers. This showed that GFP particles should not be trapped in the nano-scale porous matrix, yet their radius is less than an order of magnitude smaller than the pore size and is not negligible. This has important implications on the physics of free-GFP mobility:

1. The applicability of continuum mechanics for studying GFP mobility at the nano-scale is an approximation. The Knudsen number $K_n$ compares the mean free path of a particle in a fluid λ to a characteristic length of the flow L. If $K_n \ll 10^{-2}$ then the continuum mechanics framework is perfectly valid. Here, the two particles of interest are GFP itself and the water molecules that are

moved by free-GFP. For water, $\lambda_{H_2O} \approx 2R_{H_2O} = 0.3nm$, thus $K_n \approx 10^{-2}$. For GFP particles it is more difficult to estimate the mean free-path for GFP movement. We assumed $\lambda_{GFP} \in [2R_{H_2O}, 2R_{GFP}]$, and obtain $K_n \in [10^{-2}, 0.3]$. Thus, describing GFP diffusion at the nano-scale using continuum mechanics would be an approximation. In our modeling approach, we instead assumed that the continuum mechanics framework could be used to approximately estimate the effects of the nano-obstacles on GFP mobility *at the larger micro-scale*, on a characteristic length ≈ 1000nm.

2. Two diffusional hindrance mechanisms visible at the micro-scale arise from the presence of nano-obstacles. For one thing, nano-obstacles, which have low mobility, results in the nano-scale medium being *tortuous*. For free-GFP particles to diffuse through it, they must avoid the nano-obstacles (Fig. 4B, middle panel), resulting in longer paths and increased transport times. For another thing, when free-GFP particles are moving in the nano-scale matrix, they create a cytosolic flow around them. This flow is hindered due to the presence of the nano-obstacles. This results in an increased hydrodynamic drag for the free-GFP particles (Fig. 4B, bottom panel), leading to what we referred to as *porous hydrodynamic hindrance* This effect, which has been largely studied in the field of solute diffusion in hydrogels is present when the particle radius cannot be neglected compared to the pore size [19], [20], [21], [22], [23].

#### d. Model derivation: estimation of nano-obstacles induced diffusional hindrances

To establish a link between the diffusivity of GFP in pure cytosol $D_\alpha$ (with no nano-obstacles) and at the micro-scale $D_\gamma$, we estimated separately the nano-obstacle tortuous hindrance $T_\alpha$ and porous hydrodynamic hindrance $H_\alpha$, in accordance with Eq. 3 [24], [25].

To determine nano-obstacles tortuous hindrance $T_\alpha$, we used the method of volume averaging upscaling procedure for solute diffusion [13]. Importantly, we did not conduct an upscaling from the nano-scale to the micro-scale, but instead used the procedure to estimate $T_\alpha$ by simply solving a closure problem. For this procedure, we wrote the continuum mechanics equation for GFP diffusion in the cytosol

$$\partial_t C_\alpha = D_\alpha \nabla^2 C_\alpha \text{ in } V_\alpha$$
$$\mathbf{n}_{\alpha n} \cdot D_\alpha \nabla C_\alpha = 0 \text{ on } \mathrm{A}_{\alpha \mathrm{n}}$$

Where $C_\alpha$ is GFP cytosolic concentration, $V_\alpha$ is cytosolic accessible volume domain for GFP mass center, $\mathrm{A}_{\alpha\mathrm{n}}$ is the surface domain delimiting $V_\alpha$ from the excluded volume domain, and $\mathbf{n}_{\alpha n}$ is a unit vector normal to $\mathrm{A}_{\alpha\mathrm{n}}$. Assuming the nano-scale medium is isotropic, the procedure yields

$$T_\alpha^{-1} = 1 + \frac{1}{\mathrm{V}_\alpha} \int_{\mathrm{A}_{\alpha\mathrm{n}}} \mathbf{n}_{\alpha n} \cdot \mathbf{e}_\mathbf{j} b_\alpha \; dA$$

Where $\mathbf{e}_\mathbf{j}$ is a base unit vector and $b_\alpha$ is the closure variable solution of the following scalar closure problem

$$\nabla^2 b_\alpha = 0 \text{ on } \mathrm{V}_\alpha$$
$$-\mathbf{n}_{\alpha n} \cdot \nabla b_\alpha = \mathbf{n}_{\alpha n} \cdot \mathbf{e}_\mathbf{j} \text{ on } \mathrm{A}_{\alpha\mathrm{n}}$$
$$b_\alpha(\mathbf{x} + \mathbf{l}_\mathbf{i}) = b_\alpha(\mathbf{x}) \text{ for } i \in [1,2,3]$$

This closure problem has to be solved on a representative elementary volume (REV). In this study, the REVs are simplified periodic unit cells presented in Fig. 4C. Numerical solving of this closure problem is explained in more detail in SI Appendix 3.f.

Then, to determine $H_\alpha$, we relied on both the theory of solutal diffusion in hydrogels that expresses the dependence of the porous hydrodynamic drag on the permeability of a porous medium and the upscaling procedure for Stokes single-phase flow [13] that allows to compute the permeability for a given porous structure. Based on Brinkman [26] and Howells [27], the force exerted on a mobile spherical particle by a static flow embedded in a porous medium has been expressed as [31], [32]

$$F_p = 6\pi R_p \mu v_p \times \left[1 + \frac{R_p}{\sqrt{K}} + \frac{1}{9}\left(\frac{R_p}{\sqrt{K}}\right)^2\right]$$

Where $F_p$ is the force on the particle, $R_p$ is its radius, $\mu$ is the fluid dynamic viscosity, $v_p$ is the particle velocity and $K$ (in $m^2$) is the intrinsic permeability of the porous medium. It is noteworthy that $\sqrt{K}$ is homogeneous to a length and is of the same order of magnitude as the pore size of the porous matrix. Using Stokes-Einstein relation, [28], [29] one gets the diffusion coefficient of the particle

$$D_p = \frac{k_B T}{6\pi\mu R_p}\left[1 + \frac{R_p}{\sqrt{K}} + \frac{1}{9}\left(\frac{R_p}{\sqrt{K}}\right)^2\right]^{-1}$$

When the particle size is negligible compared to the characteristic pore length scale $\sqrt{K}$, one recovers the well-known Stokes-Einstein relation for a dilute particle in a pure fluid $D_p = k_B T/6\pi\mu R_p$. We can therefore identify the porous hydrodynamic hindrance as the correction to the particle diffusivity induce by the presence of the porous medium [29], here applied for GFP at the nano-scale medium

$$H_\alpha = 1 + \left(\frac{R_{GFP}}{\sqrt{K_\alpha}}\right) + \frac{1}{9}\left(\frac{R_{GFP}}{\sqrt{K_\alpha}}\right)^2$$

Where $R_{GFP} = 2.3nm$ is GFP radius and $K_\alpha$ is the nano-scale medium intrinsic permeability.

In this study, we compute $K_\alpha$ based on the structure of the nano-scale medium using the method of volume averaging for single phase Stokes-flow. Consistently with the hypotheses previously formulated, we considered the continuum mechanics framework valid for studying the cytosolic fluid flow in the nano-scale medium. The cytosolic incompressible, low Reynolds viscous flow can be described with the equations

$$0 = -\nabla p_\alpha + \mu_\alpha \nabla^2 \mathbf{v}_\alpha \text{ on } V_\alpha^*$$
$$\nabla \cdot \mathbf{v}_\alpha = 0 \text{ on } V_\alpha^*$$
$$\mathbf{v}_\alpha = 0 \text{ on } A_{\alpha n}^*$$

Where $p_\alpha$ is the cytosol pressure, $\mathbf{v}_\alpha$ is cytosolic flow speed, $\mu_\alpha$ is cytosolic viscosity (increased due to the presence of viscogens), $V_\alpha^*$ is cytosolic physical volume domain, $A_{\alpha n}^*$ is the surface domain delimiting $V_\alpha^*$ from the nano-obstacles occupied volume domain, and $\mathbf{n}_{\alpha n}$ is a unit vector normal to $\mathrm{A}_{\alpha\mathrm{n}}^*$. The first equation is the momentum conservation equation, the second is the mass conservation equation, and the third is the no-slip boundary condition on the surface of the nano-obstacles. With the method of volume averaging [13], assuming an isotropic nano-scale medium, the permeability tensor $\mathbf{K}_\alpha$ is diagonal and we have

$$\mathbf{K}_\alpha = K_\alpha \mathbf{I} = \epsilon_\alpha^* \langle \mathbf{B}'_\alpha \rangle^\alpha$$
$$\langle \mathbf{B}'_\alpha \rangle^\alpha = \frac{1}{V_\alpha^*}\int_{V_\alpha^*} \mathbf{B}'_\alpha \quad dV$$

And then in the, we estimated the isotropic scalar permeability $K_\alpha$ (used in the rest of our study) by solving the following closure problem where $\mathbf{B}'_\alpha$ is an order one tensor variable field and $\mathbf{b}'_\alpha$ is a scalar closure variable

$$
\begin{aligned}
&0 = -\nabla \mathbf{b}'_\alpha + \nabla^2 \mathbf{B}'_\alpha + \mathbf{I} \quad \text{on } V_\alpha^* \\
&\nabla \cdot \mathbf{B}'_\alpha = 0 \quad \text{on } V_\alpha^* \\
&\mathbf{B}'_\alpha = 0 \quad \text{on } A_{\alpha n}^* \\
&\langle \mathbf{b}'_\alpha \rangle^\alpha = 0 \quad \text{on } V_\alpha^* \\
&\mathbf{b}'_\alpha(r + l_i) = \mathbf{b}'_\alpha(r)\,;\ \mathbf{B}'_\alpha(r + l_i) = \mathbf{B}'_\alpha(r) \ \text{ for i} \in [1,2,3]
\end{aligned}
$$

This closure problem must be solved on a REV. In this study, the REVs we considered are simplified periodic unit cells presented in Fig. 4C. Numerical solving of this closure problem is explained in more detail in SI Appendix 3.f.

#### e. Micro-scale physical analysis and model derivation: estimation of micro-obstacle induced tortuous hindrance

We investigated how micro-obstacles may impact GFP diffusion in the cytoplasm and estimated their tortuous hindrance $T_\gamma$. The physical analysis for micro-obstacles impact on GFP diffusion is much simpler than for nano-obstacles: with a size in the range 100 − 1000nm, the micro-obstacles are much larger than GFP, thus the volume excluded by micro-obstacles is also the volume they occupy. The micro-scale pore size is much larger than GFP size, thus it played no role in our study and we did not estimate it. Moreover, the micro-obstacles were assumed to be non-mobile, non-deformable, and non-permeable to free-GFP.

For a typical free-GFP protein concentration of $C_\gamma \approx 1\mu M$, the number of free-GFP particles in a micro-scale volume ≈ 1µm3 is expected to be about 300 on average. We thus proposed to describe the spatial distribution of free-GFP in the hyaloplasmic γ-phase using a continuous concentration field $C_\gamma$. The framework of continuum mechanics was thus applicable, and we described free-GFP transport in the γ-phase using a diffusion equation.

To link GFP diffusion in the hyaloplasm to GFP diffusion in the cytoplasm, we conducted a rigorous upscaling from the micro-scale to the cytoplasm scale. We assumed the micro-scale medium to be isotropic and uniform in each cytoplasm region (HP or LP). GFP diffusion in the hyaloplasm is driven by

$$
\begin{aligned}
\partial_t C_\gamma &= D_\gamma \nabla^2 C_\gamma \quad on\ V_\gamma \\
-\mathbf{n}_{\gamma m} \cdot D_\gamma \nabla C_\gamma &= 0 \quad on\ A_{\gamma m}
\end{aligned}
$$

Where $V_\gamma$ is hyaloplasm volume domain, $A_{\gamma m}$ is the surface domain delimiting $V_\gamma$ from the micro-obstacles volume domain $V_m$, and $\boldsymbol{n}_{\gamma m}$ is a unit vector normal to $A_{\gamma m}$.
Note that $D_\gamma$ is an effective diffusion coefficient that accounts for the transport mechanisms identified at the nano-scale. It is considered constant withing the micro-scale averaging region. The upscaling procedure yields

$$\partial_t \langle C_\gamma \rangle^\gamma = D_{eff} \nabla^2 \langle C_\gamma \rangle^\gamma$$

Where the effective diffusivity of GFP in the cytoplasm is given by

$$\frac{D_{eff}}{D_\gamma} = T_\gamma^{-1} = 1 + \frac{1}{V_\gamma} \int_{A_{\gamma m}} \mathbf{n}_{\gamma m} \cdot \boldsymbol{e_i} b_\gamma \quad dA$$

In which $b_\gamma$ is a closure variable that can be computed by solving the following closure problem

$$\nabla^2 b_\gamma = 0 \quad \text{on } \mathrm{V_\gamma}$$
$$-\mathbf{n}_{\gamma m} \cdot \nabla b_\gamma = \mathbf{n}_{\gamma m} \cdot \mathbf{e_j} \quad \text{on } \mathrm{A_{\gamma m}}$$
$$b_\gamma(\mathbf{x} + \mathbf{l_i}) = b_\gamma(\mathbf{x}) \quad \text{for } i \in [1,2,3]$$

This closure problem has to be resolved on a REV In this study, the REVs we considered are simplified periodic unit cells presented in Fig. 4C. Numerical solving of this closure problem is explained in more detail in SI Appendix 3.f.

##### f. Numerical resolution of the closure problems

The closure problems for $T_\gamma$, $T_\alpha$ and $H_\alpha$ obtained using the method of volume averaging were solved numerically on four types of periodic REVs (also summarized in Fig. 4C):

- 2D-CD-F-actin: a 2D periodic square cytosolic domain (α-phase) with a centered disk obstacle (n-phase) with a radius $R = R_{act} = 4nm$. Mimicks F-actin-like nano-obstacles for a low computational cost. Detailed mesh is presented in Fig. S14.

- 3D-CFC-ribosome: a 3D periodic cube cytosolic domain (α-phase) with spheric obstacles in a cubic-face-centered arrangement (n-phase). The spheres have a radius $R = R_{rib} = 12.5nm$. Mimicks ribosome-like nano-obstacles, at an intermediate computational cost. Detailed mesh is presented in Fig. S15.

- 3D-random-F-actin: a 3D periodic cube cytosolic domain (α-phase) with randomly placed and oriented elongated cylindrical fibers (n-phase). The fibers have a radius $R = R_{act} = 4nm$. This geometry mimics F-actin-like nano-obstacles more realistically than the 2D-CD geometry but at a great computational cost. Detailed mesh is presented in Fig. S16.

- 3D-random-mixed: a 3D periodic cube cytosolic domain (α-phase) with randomly placed and oriented elongated cylindrical fibers and spheres (n-phase). The fibers have a radius $R_{act} = 4nm$, and the spheres have a radius $R_{rib} = 12.5nm$. This geometry mimics a mixture of F-actin-like and ribosome-like nano-obstacles at a great computational cost. Detailed mesh is presented in Fig. S17.

- 3D-CFC-micro: This configuration is geometrically identical to the 3D-CFC-ribosome but with a radius R representative of microscale obstacles. Detailed mesh is presented in Fig. S15.

To investigate the different effects of nano-obstacles abundance, we varied $\epsilon_n$ and $\epsilon_n^*$ by generating different geometries for each REV presented. For the 2D-CD, the 3D-CFC, and the 3D-random-F-actin geometries, different $\epsilon_n$ and $\epsilon_n^*$ values were obtained by changing the numerical radius of the obstacles while keeping the same periodic domain numerical size. The corresponding physical size for the obstacles (either $R_{act}$ or $R_{rib}$) was kept constant. For the 3D-random-mixed geometry, different values of $\epsilon_n$ and $\epsilon_n^*$ were obtained by changing the number of obstacles while keeping the fiber:sphere number ratio to 2:1.

For the tortuosity closure problems (for $T_\gamma$ and $T_\alpha$), the REV geometries refer to the effective porous medium geometry for free-GFP movement[1]: the numerical obstacle fractions thus refer to the excluded volume fractions $\epsilon_n$ and $\epsilon_m$. For the permeability closure problem $H_\alpha$, the REV geometries refer to the physical nano-scale geometry: the numerical nano-obstacle fraction thus

[1] It is noteworthy that at the micro-scale, the volume fraction excluded by micro-obstacles for GFP diffusion is identical to the volume fraction they occupy, as GFP size is negligible compared to the typical micro-obstacle size

refers to the volume fraction occupied by the nano-obstacles $\epsilon_n^*$. Importantly, to plot not only $T_\alpha$ but also $H_\alpha$ as functions of $\epsilon_n$, it was important to link $\epsilon_n^*$ to $\epsilon_n$. This was done in SI Appendix 3.c for non-intersecting excluded volumes, an assumption that is only reasonable for 2D-CD and 3D-CFC configurations. For 3D-random configurations, we numerically computed the ratio between $\epsilon_n^*$ and $\epsilon_n$ by generating numerical geometries corresponding to the same physical configuration (same number of obstacles), but with different numerical obstacle sizes corresponding to either the physical nano-scale medium geometry (computation of $\epsilon_n^*$) or the effective nano-scale medium geometry for GFP mass-center (computation of $\epsilon_n$). The results are presented in Fig. S25.

For each of these geometries, the mesh was generated using Gmsh python API. To allow for the application of a periodic boundary condition on the REV, the mesh at the opposed faces of the square in 2D (and of the cube in 3D) were imposed identical: to each mesh element on a face corresponded a unique mesh element on the opposite face. The mesh was refined near the obstacles' surface to optimize computation. For all results, a mesh sensibility analysis was carried to ensure a negligible influence of the discretization on the computed solutions.
The meshes generated with Gmsh (.msh format) were then converted to a format compatible with FEniCS (.xdmf format) using the Meshio python package. The .xdmf files were read by FEniCS python package. The variational formulations of the closure problems were then derived and given as an input for the numerical finite element method using FEniCS. For the tortuosity closure problems ($T_\alpha$ and $T_\gamma$), which only have a pure Neumann boundary condition on the surface of the obstacles and a periodicity condition on the other domain boundaries, the weak formulation is

$$\int_\Omega (\nabla u \cdot \nabla v + cv + ud)\ dx = -\int_{\partial\Omega} (\boldsymbol{n} \cdot \boldsymbol{e_j}) v\ dS$$

Where $u$ is the trial scalar function, $v$ is the test scalar function, $c$ and $d$ are scalar constants, $\Omega$ is the 2D or 3D fluid domains, $\boldsymbol{n}$ is the unit vector normal to the fluid-obstacle interface, $\boldsymbol{e_j}$ is the arbitrary unit vector already present in the closure problem, and $\partial\Omega$ is the 1D or 2D interface between the fluid and obstacle domains. This problem was solved using the Gmres iterative solver. For the scalar version of the nano-scale permeability closure problem, a mixed variational formulation was used. The variational formulation of the problem and the associated preconditioner system are

$$\int_\Omega \nabla \boldsymbol{u} : \nabla \mathbf{v} + \mathrm{p}(\nabla \cdot \mathbf{v}) + \mathrm{q}(\nabla \cdot \mathbf{u})\, dx = \int_\Omega \boldsymbol{e_j} \cdot \boldsymbol{v}\ dx$$
$$\int_\Omega \nabla \boldsymbol{u} : \nabla \mathbf{v} + pq\ dx = \int_\Omega \boldsymbol{e_j} \cdot \boldsymbol{v}\ gx$$

Where $(\boldsymbol{u}, p)$ is the couple of trial functions, $(\boldsymbol{v}, q)$ are the test functions, $\boldsymbol{e_j}$ is the arbitrary unit vector already present in the closure problem, and $\Omega$ is the 2D or 3D fluid domain. When using FEniCS, Dirichlet boundary conditions do not appear in the weak formulation. Here, the no-slip boundary condition on the obstacles surfaces does not appear in the weak formulation, but it is still considered in the resolution of this weak formulation. This problem was solved using a TFQMR solver along with an AMG preconditioner.

To ensure our numerical implementation using FEniCS was correct, we compared our results for tortuosity and permeability computation with literature results. Three geometries were assessed: 2D-Centered Disk (same as the 2D-CD we used in our study), 2D-Centered Square, and 2D-Face Centered Squares. For tortuosity closure problem, the results from [33] were expressed in quantities equivalent to porosity $\epsilon$ and porosity times tortuosity $\epsilon \times T^{-1}$, so we presented our result using these quantities. The results are presented in Fig. S26 and show good agreement. The resolution of the closure problem in permeability was validated using the 2D results from [34] and using the 3D results from [35]. In 2D, we used the 2D Centered Disk geometry. In 3D, we used the 3D Centered Sphere geometry. The results from [35] were expressed in terms equivalent to $(1-\epsilon)^{1/3}\,/\,\epsilon$ and to the drag coefficient $C = \left(\frac{2}{9}\right) \times \left(\frac{R^2}{(1-\epsilon)}\right) \times K^{-1}$ where R is the sphere radius, so we presented our 3D results using these quantities. The 2D and 3D results are presented in Fig. S27 and S28, showing good agreement with literature results.

#### 4. Supplementary developments for model comparison with experiments

##### a. Cytosolic diffusivity dependence on osmolarity and correction

Due to the presence of numerous small solutes that act as viscogens, the cytosol is more viscous than water, hindering diffusion of macromolecules such as GFP. The cytosolic diffusivity of GFP $D_\alpha$ was left as a fitting parameter for comparison with experimental data for each osmotic condition separately in Fig. 5A-C. The fitted $D_\alpha$ values for each osmolarity allows to express the dependence of cytosolic viscosity on the osmotic condition present in Fig. 5D. As we wanted to understand this dependence and to compare the experimental data for all osmolarities with the full model in a single fitting procedure, it was necessary to model the dependence of cytosolic diffusivity on the osmotic condition to propose a correction for this dependence.

We assumed that the cytosol α-phase is homogeneous inside the cell. Though many small solutes may be subject to interactions with specific structures and be heterogeneously distributed in the cytosol, the solutes are so diverse that it is reasonable to assume the total viscogens concentration is nearly uniform in the α-phase. We also assumed that the cytosolic fluid (thus excluding nano- and micro- obstacles) may be described as a dilute solution. With these hypotheses, the diffusion of free-GFP in pure cytosol was described using the Stokes-Einstein relation

$$D_\alpha = \frac{k_b T}{6\pi \mu_\alpha R_{GFP}}$$

Where $k_b$, $T$, $\mu_\alpha$, and $R_{GFP} = 2.3nm$ are respectively the Boltzmann constant, the temperature, the cytosol dynamic viscosity, and GFP radius. The relative cytosolic to water diffusivity for free-GFP can thus be written as

$$\frac{D_\alpha}{D_{H_2O}} = \frac{\mu_{H_2O}}{\mu_\alpha}$$

Where $\mu_{H_2O}$ is the water dynamic viscosity and $D_{H_2O}$ is the free-GFP diffusivity in pure water. As the cytosol was assumed diluted, it is possible to use the ideal solution viscosity model [36]

$$\frac{\mu_\alpha}{\mu_{H_2O}} = 1 + kC_t$$

Where $k$ is a constant depending on the nature of the solutes and $C_t$ is the total cytosolic viscogens concentration (in molars). Along with Stokes-Einstein's relation, this yields

$$D_\alpha = D_{H_2O} \times \frac{1}{1 + kC_t}$$

Here knowledge of $kC_t$would allow us to estimate $D_\alpha$ However, estimating $kC_t$ accurately is difficult. We thus derived from this equation another expression that would be more useful in our study, by focusing on the dependence of $D_\alpha$ on osmolarity rather than trying to know the absolute $D_\alpha$ value. For a cell submitted to an arbitrary osmotic shock, we have after the hyper-osmotic shock

$$C_t = C_t^0 \times \frac{V_{cytosol}^0}{V_{cytosol}} \approx C_t^0 \times V_r^{-1}$$

Where for any variable $A$, we note $A^0$ its value in iso-osmotic condition, before the hyper-osmotic shock, with $V_{cytosol}$ the cell cytosolic volume and $V_r$ the cell volume ratio after/before the osmotic shock as presented in Fig. 2D. We defined $k' = kC_t^0$, and by using the above equations we obtained the equation for $D_\alpha/D_\alpha^0$ as

$$\frac{D_\alpha}{D_\alpha^0} = \frac{1 + k'}{1 + k'V_r^{-1}}$$

This relation links the cytosolic diffusivity at a given osmolarity $D_\alpha$ to the cytosolic diffusivity in iso-osmotic condition $D_\alpha^0$. We then wanted to obtain an estimate for $k'$. A first trivial estimate could be obtained by observing that $D_{eff} < D_\alpha^0 < D_{H_2O}$. From our FRAP results in iso-osmotic condition $D_{exp} \approx 20\ \mu m^2 s^{-1}$ (in HP regions, Fig 3C), and from the literature estimate for $D_{H_2O} = 87 \mu m^2 s^{-1}$ [18], [26], [37], [38], we obtained

$$0 \leq k' \leq 3.5$$

A more precise estimate was obtained using literature rotational mobility measurements [37], [39], [40]. In the cytoplasm, these measurements allow for probing the effects of the immediate particle environment on its diffusive mobility. For small enough molecules, rotational mobility measurements are thus believed to be mainly impacted by the cytosolic fluid phase viscosity rather than by cytoplasmic structures such as the nano- and micro-obstacles. Assuming the characteristic rotation time $\tau_\alpha^r$ for small solutes in the cytosol measured in these studies is proportional to the cytosol viscosity $\mu_\alpha$, we obtained

$$\frac{D_{H_2O}}{D_\alpha} = \frac{\mu_\alpha}{\mu_{H_2O}} = \frac{\tau_\alpha^r}{\tau_{H_2O}^r} \approx 1.5 - 2.5$$

With $\tau_{H_2O}^r$ the diffusive particle characteristic rotation time in pure water. This leads to

$$0.5 \leq k' \leq 1.5$$

While this estimate is still quite rough, the aforementioned equation for $D_\alpha/D_\alpha^0$ does not predict a strong variability of $D_\alpha/D_\alpha^0$ on $k'$ for $k' \in [0.5,1.5]$ (Fig. 5D). For instance, for the 600mOsm hyper-osmotic condition, if we assume $k' = 1 \pm 0.5$, we obtain $D_\alpha/D_\alpha^0 \in [0.806,0.882]$, which represents a relative error of only ±4.5%, which was considered acceptable for our study.

With the equation for $D_\alpha/D_\alpha^0$ this equation and the estimate $k' = 1 \pm 0.5$, we could correct our experimental FRAP results $D_{exp}$ and the model effective cytoplasmic diffusivity $D_{eff}$ for the cytosolic diffusivity osmotic variation. We then obtained the correction for cytosolic diffusivity variation with osmotic condition as

$$\frac{D_\alpha^0}{D_\alpha} = \frac{1 + k'V_r^{-1}}{1 + k'}$$

To obtain GFP experimental FRAP diffusivity corrected for cytosolic diffusivity variation with osmotic condition $D'_{exp}$ and GFP model effective cytoplasmic diffusivity corrected for cytosolic diffusivity variation with osmotic condition $D'_{eff}$, we simply multiplied $D_{exp}$ and $D_{eff}$ by $\frac{D_\alpha^0}{D_\alpha}$

$$D_{exp} \times \frac{D_\alpha^0}{D_\alpha} = D_{exp} \times \frac{1 + k'V_r^{-1}}{1 + k'}$$

$$D_{eff} \times \frac{D_\alpha^0}{D_\alpha} = \left(T_\gamma^{-1} \times T_\alpha^{-1} \times H_\alpha^{-1} \times D_\alpha\right) \times \frac{D_\alpha^0}{D_\alpha} = T_\gamma^{-1} \times T_\alpha^{-1} \times H_\alpha^{-1} \times D_\alpha^0$$

In iso-osmotic conditions we have $V_r = 1$ and thus $D'_{eff} = D_{eff}$ and $D'_{exp} = D_{exp}$. $D'_{eff}$ and $D'_{exp}$ are physically comparable, and $D'_{eff}$ only depends on $D_\alpha^0$ which does not depend on the osmotic condition. This allowed used to use $D_\alpha^0$as the only fitting parameter to fit our model with all our osmotic results in one go in Fig. 5E.

##### b. Bootstrapping for assessment of uncertainty in model-experiment fitting procedures

Bootstrapping from python package Scipy (scipy.stats.bootstrap) was used to obtain a standard deviation value of the fitted parameters $D_\alpha$ and $D_\alpha^0$ and of the RMSE for each comparison between the model and the osmotic experiments (Fig 5). For each fit plotted, the experimental osmotic data was resampled 1000 times cell wise (meaning that for an individual cell data $(\Phi_n,\ D_{exp}), \Phi_n$ was always taken into account as many times as $D_{exp}$). Resampling was conducted using always the same random number generation kernel to ensure repeatability. The standard deviation of the 1000 values obtained for $D_\alpha$, $D_\alpha^0$, and the RMSE was then presented in Fig. 5A-C, E. To obtain vertical standard deviations in Fig. 5D, uncertainty propagation was computed according to

$$\left(\boldsymbol{\sigma}_{\left(\frac{\boldsymbol{a}}{\boldsymbol{b}}\right)}\right)^2 \approx \left(\frac{\boldsymbol{a}}{\boldsymbol{b}}\right)^2 \left[\left(\frac{\boldsymbol{\sigma_a}}{\boldsymbol{a}}\right)^2 + \left(\frac{\boldsymbol{\sigma_b}}{\boldsymbol{b}}\right)^2\right]$$

##### c. Model application to particle with a radius $R < 10nm$

We applied our model to particles other than GFP. Importantly, we restricted the validity of our model to $R < 10nm,$ as for larger particles the validity of our model hypotheses (see SI Appendix 3.b) may not be considered as valid anymore: an important part of what we considered as nano-obstacles (such as ribosomes) for studying free-GFP diffusion may no longer be considered immobile on the timescales of particle diffusion, and consequent direct mechanical interactions (fibers and membrane deformations) would be expected between the diffusive macromolecule and the nano-obstacles. Also, we focused only on iso-osmotic condition, on an average cytoplasm region, with properties between those of HP and LP regions that we observed in our experiments.

First, we used our experimental data for the effective porous medium as perceived by GFP mass-center ($\Phi_n$ and $\Phi_m$ data from Fig. 1I) to deduce the porous medium as perceived by other macromolecules with $R < 10nm$. First, from the proposed model decomposition of the cytoplasm into three spatial scales, we know that the hyaloplasmic nano-obstacle excluded volume fraction $\epsilon_n$ and the cytoplasmic nano-obstacle excluded volume fraction $\Phi_n$ are linked by

$$\epsilon_n = \frac{\Phi_n}{1 - \Phi_m}$$

Then, from SI Appendix 3.c, by assuming the nano-obstacles have a shape closure to ribosomes (spheres with 12.5nm radius) we have that the hyaloplasmic nano-obstacle occupied volume fraction $\epsilon_n^*$ is

$$\epsilon_n^* = \frac{\Phi_n}{1.66\ (1 - \Phi_m)}$$

From our experimental data in Fig. 1I, we obtain depending on the cytoplasmic region considered

$$\epsilon_n^* \in [0.10, 0.18]$$

We chose $\epsilon_n^* = 0.12$ as representative of an average cytoplasm structure. Importantly, $\epsilon_n^*$ describes the intrinsic porous medium structure and is not dependent on the particle size, using this variable is thus relevant for all radii values in $R < 10nm$.

Then, we estimated the porous hindrances for different diffusive particle sizes. The micro-obstacle hindrance $T_\gamma$ was neglected here as for $\Phi_m = 0.08$ its impact is limited. Moreover, for particles with $R < 10nm$, $T_\gamma$ is nearly independent on particle size (excluded volume effects are negligible). Thus we solely focused on estimating $T_\alpha$ and $H_\alpha$. Accordingly with SI Appendix 3.d, $H_\alpha$ was estimated by

$$H_\alpha = 1 + \left(\frac{R}{\sqrt{K_\alpha}}\right) + \frac{1}{9}\left(\frac{R}{\sqrt{K_\alpha}}\right)^2$$

Where $K_\alpha$ is the intrinsic nano-scale medium permeability and thus does not depend on the diffusive particle considered. $T_\alpha$ was estimated by recomputing the nano-obstacle excluded volume fraction $\epsilon_n$ for each particle radius from $\epsilon_n^*$ with the relation (see SI Appendix 3.c)

$$\frac{\epsilon_n}{\epsilon_n^*} = \left(\frac{R + R_{rib}}{R_{rib}}\right)^3$$

Then, as for a given REV geometry the tortuosity $T_\alpha$ is only dependent on $\epsilon_n$ (and not directly on particle size), we reused and adapted results presented in Fig. 4E for 3D-CFC Ribosomes. The obtained theoretical function for $D_{eff}$ is plotted in Fig. S20.

To assess the relevance of these predictions it was crucial to compare it with experimental results. In the literature, different works observed different levels of dependence of the relative cytoplasmic to water diffusivity $D_{eff}/D_{H_2O}$ on the particle radius $R$. Several studies reported a dependance of $D_{eff}/D_{H_2O}$ on the particle radius $R$ that was stronger than Stokes-Einstein relation predictions [18], [41], [42], [43], [44].

To compare model predictions with experimental results, we thus used literature results from three articles that assessed the cytoplasmic effective diffusivity of several nanometric inert macromolecules of different radii R<10nm [18], [41], [43]. Indeed, comparing our results with these literature results might be impacted by several biases: the cell lines are different, and the abundance of nano-obstacles in indeed not studied in the original articles, hence we proposed to use $\epsilon_n^* = 0.12$ as an estimate for all results.

More importantly, we needed to find comparable model and experimental quantities. While the multiscale model directly informs us on $D_{eff}/D_\alpha$ (without any fitting procedure), literature experiments are expressed in relative cytoplasmic to water particle diffusivity $D_{exp}/D_{H_2O}$: the two cannot be compared directly. Hence, we derived the normalized model and experimental cytoplasmic diffusivities, as presented in Eq. 7. It is clear that $D_{exp}^*$ can be computed from the experimental diffusivity results expressed in $D_{exp}/D_{H_2O}$, and that $D_{eff}^*$ can be computed from our multiscale model without any fitting procedure.

Then it was also necessary to show that $D_{exp}^*$ and $D_{eff}^*$ are physically comparable. From the Stokes-Einstein relation, which we assumed valid in pure cytosol $\alpha$-phase, we obtained that the quantity $D_\alpha/D_{H_2O}$ does not depend on particle size for a given cytosolic viscosity (which may vary between different cell lines). From this, we observed that

$$D_{eff}^{*} = \frac{\left(\frac{D_{eff}}{D_{\alpha}}\right)_{R}}{\left(\frac{D_{eff}}{D_{\alpha}}\right)_{R_{ref}}} \times 1 = \frac{\left(\frac{D_{eff}}{D_{\alpha}}\right)_{R}}{\left(\frac{D_{eff}}{D_{\alpha}}\right)_{R_{ref}}} \times \frac{\left(\frac{D_{\alpha}}{D_{H_2O}}\right)_{R}}{\left(\frac{D_{\alpha}}{D_{H_2O}}\right)_{R_{ref}}} = \frac{\left(\frac{D_{eff}}{D_{H_2O}}\right)_{R}}{\left(\frac{D_{eff}}{D_{H_2O}}\right)_{R_{ref}}}$$

Which confirmed the relevance of comparing $D_{exp}^{*}$ and $D_{eff}^{*}$. Notably, this normalization also allowed for the comparison of experimental results from different cell lines, as differences in cytosolic viscosities do not impact $D_{exp}^{*}$ and $D_{eff}^{*}$. We set $R_{ref} = 3.5nm$ so that $\left(\frac{D_{exp}}{D_{H_2O}}\right)_{R_{ref}}$ could be estimated for the three literature datasets using linear interpolation of their results.

#### 5. Supporting Figures

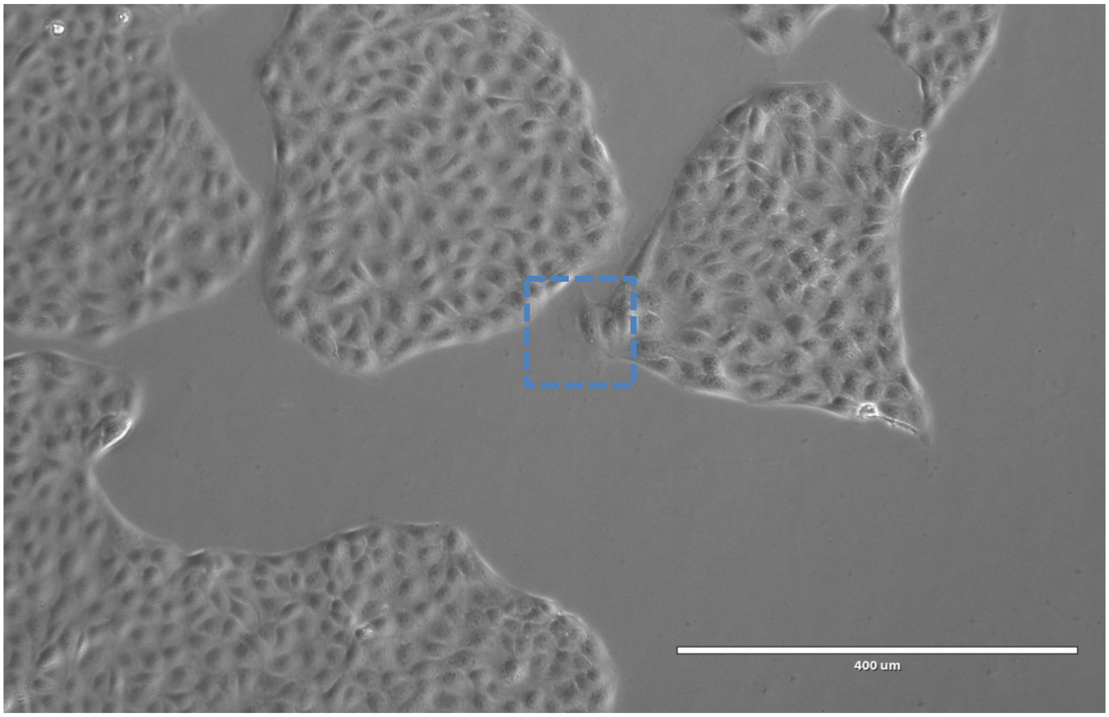


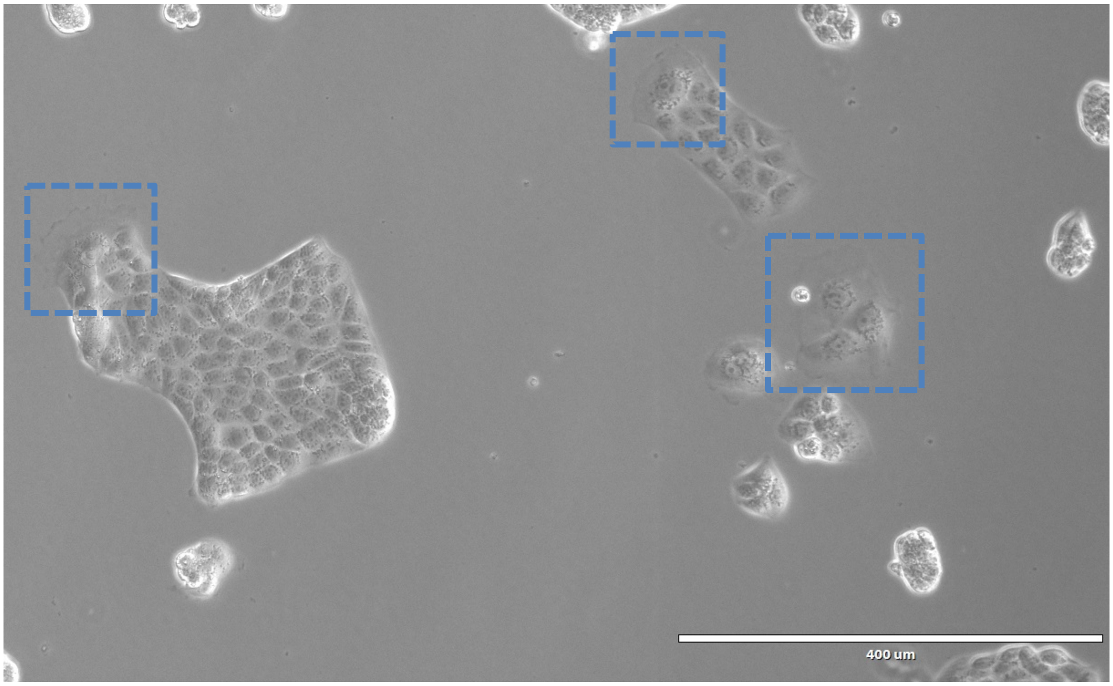


**Fig. S1.** Phase contrast images of MDCK cells in culture in a flask (top image) and on a fluorodish on the day of experiment (bottom image). Large and spread cells are present at the border of the cell islands (blue squares).

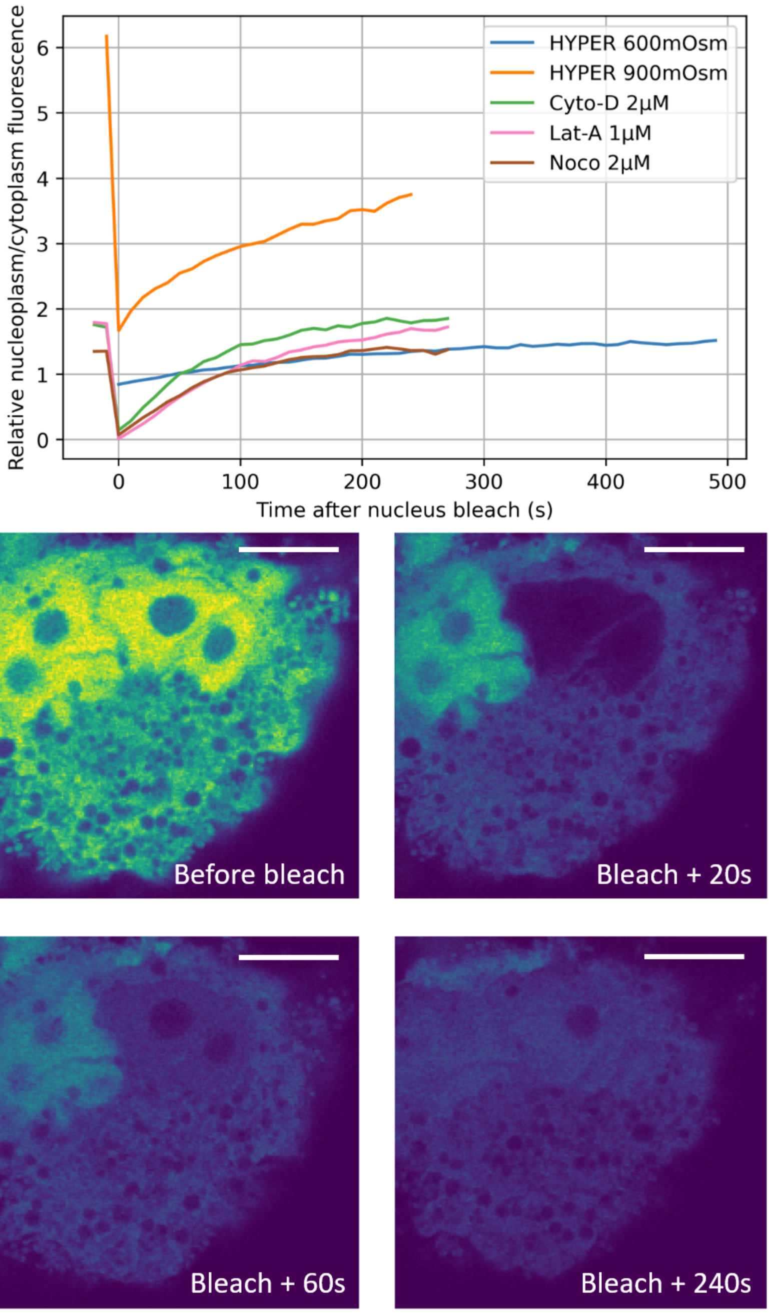


**Fig. S2.** We verified that Free-GFP could freely diffuse in and out of the nucleus by photobleaching nucleoplasm Free-GFP particle and observing a fluorescence recovery in a few minutes, for various osmotic conditions and drug treatments. Example images were obtained on a cell treated with Latrunculin-A 1µM.

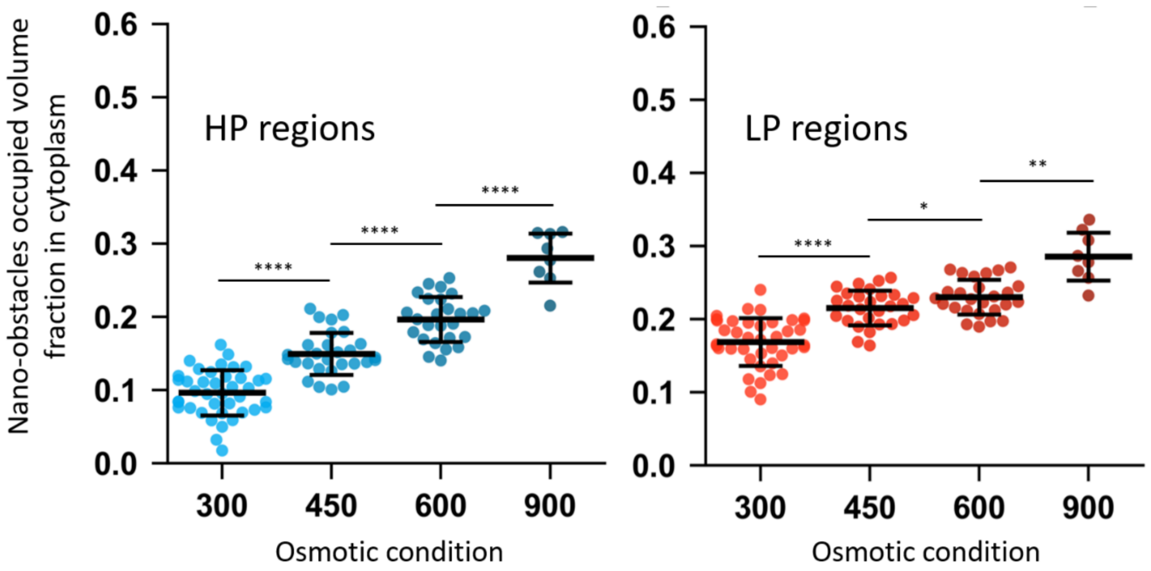


**Fig. S3.** Volume fraction occupied by nano-obstacles, estimated from Fig 1I using development presented in SI Appendix 3.c and assuming nano-obstacles are geometrically comparable to ribosomes, with non-intersecting excluded volumes. Micro-obstacles physically occupied volume fractions are identical to their excluded fractions presented in Fig. 2G. Horizontal bars are mean and ±STD. N cells=40 for 300mOsm, 28 for 450mOsm, 26 for 600mOsm and 8 for 900mOsm. Welch's t-test: *$p < 5e-2$ ; **$p < 1e-2$; ***$p < 1e-3$ ; ****$p < 1e-4$.

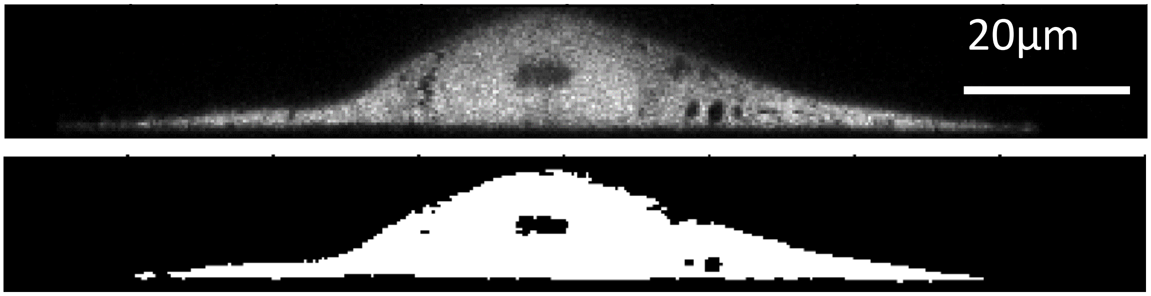


**Fig. S4.** Raw image and segmented image of a MDCK cell using GFP fluorescence signal, for inference of cell volume.

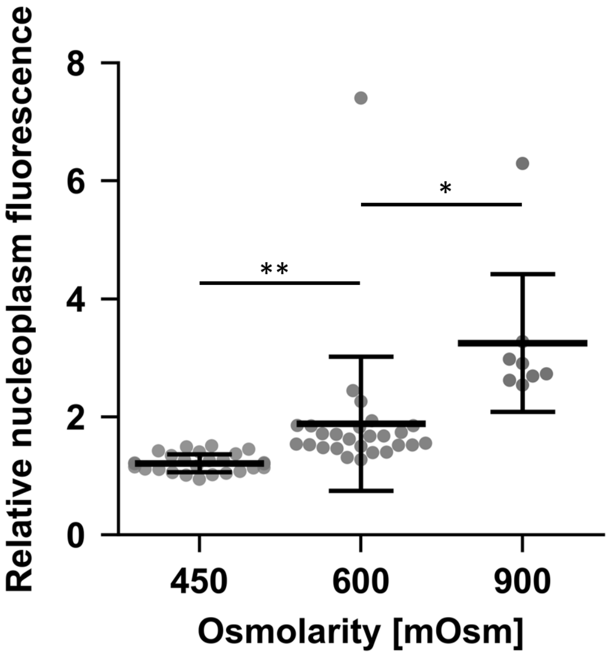


**Fig. S5.** Nucleoplasmic fluorescence reference level after hyper-osmotic shock relative its fluorescence level before osmotic-shock. Horizontal bars are mean and ±STD. N cells=28 for 450mOsm, 26 for 600mOsm and 8 for 900mOsm. Welch's t-test: *p < 5e − 2 ; **p < 1e − 2; ***p < 1e − 3 ; ****p < 1e − 4.

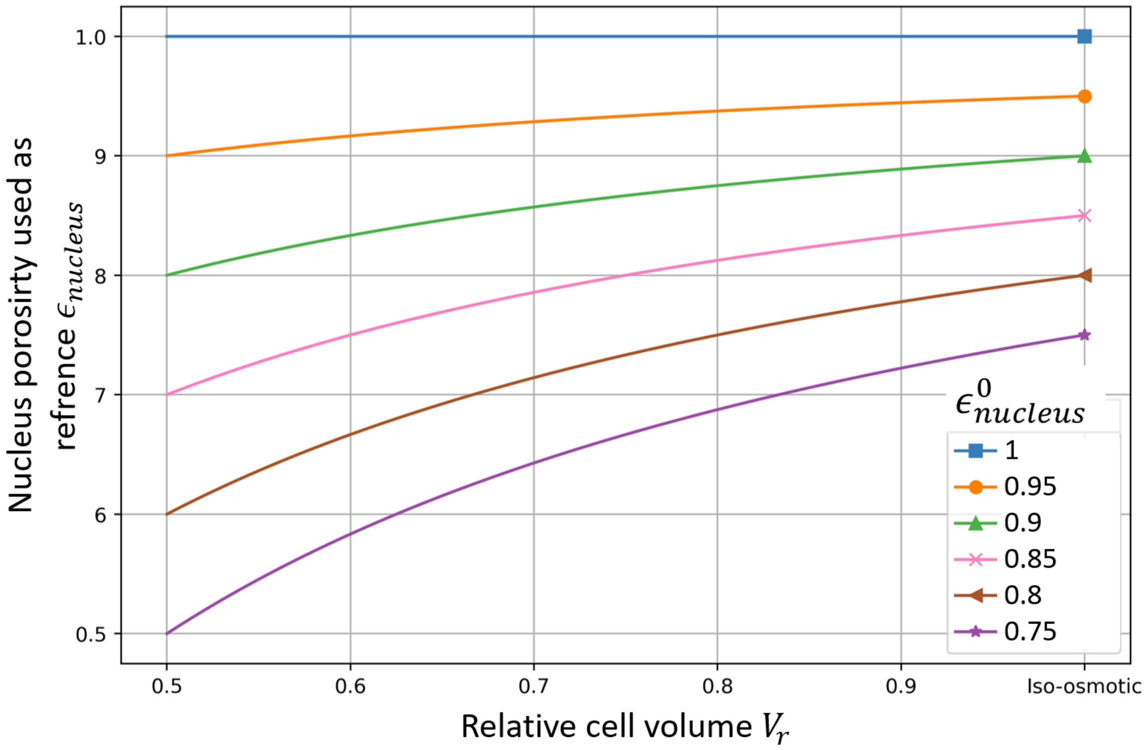


**Fig. S6.** Nucleoplasmic accessible volume fraction evolution with osmolarity for different iso-osmotic $\epsilon^0_{nucleus}$ values.

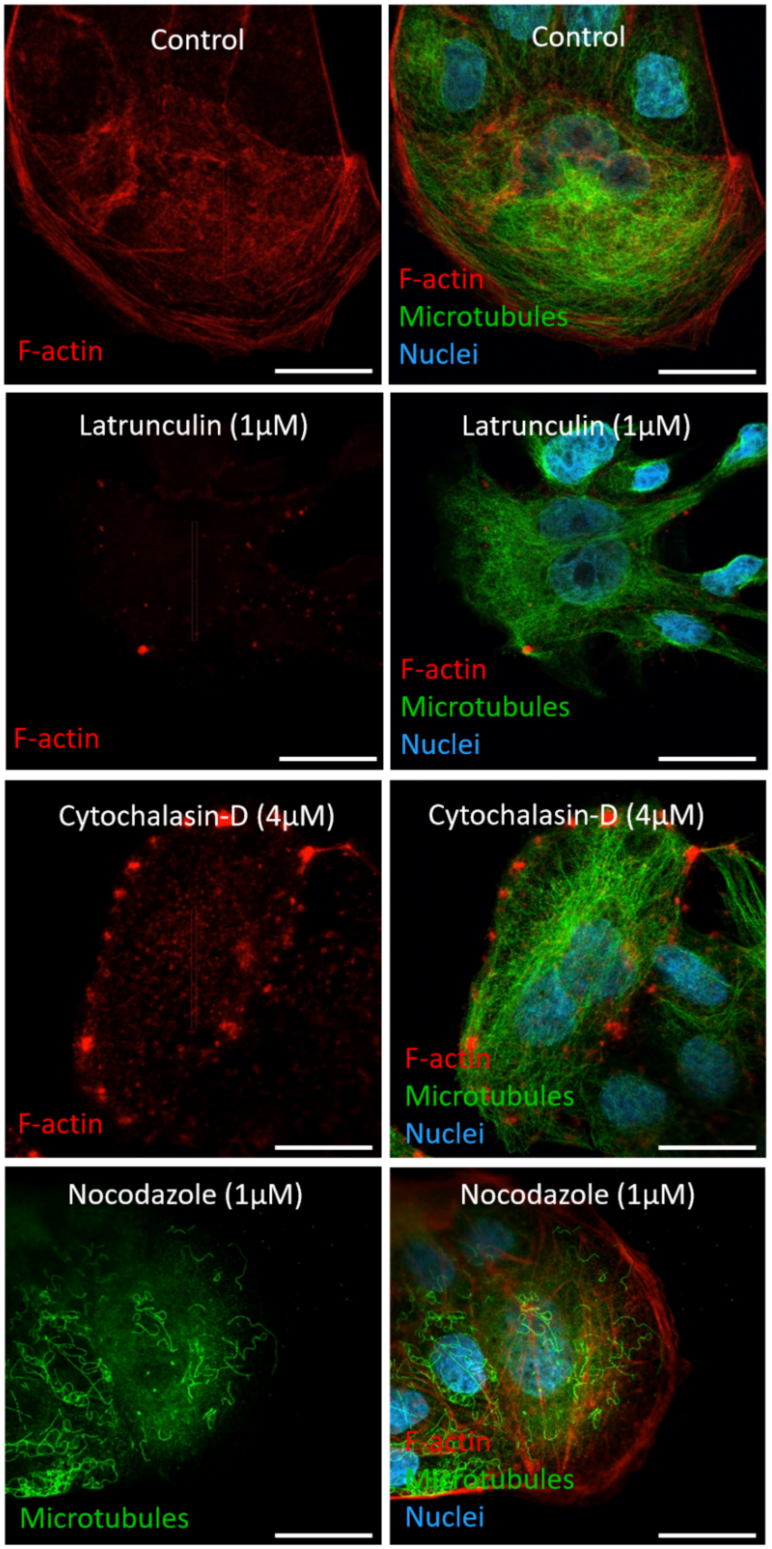


**Fig. S7.** Confocal fluorescence images of fixed cells with stained F-actin (red), microtubules (green), and nuclei (blue) for control cells (top), Latrunculin-A 1 μM (second line), Cytochalasin-D 4μM (third line), and Nocodazole 1μM (bottom). To allow comparison between different lines, for a given color the signal of the images from every line is normalized by the same value. Scale bar is 20μm.

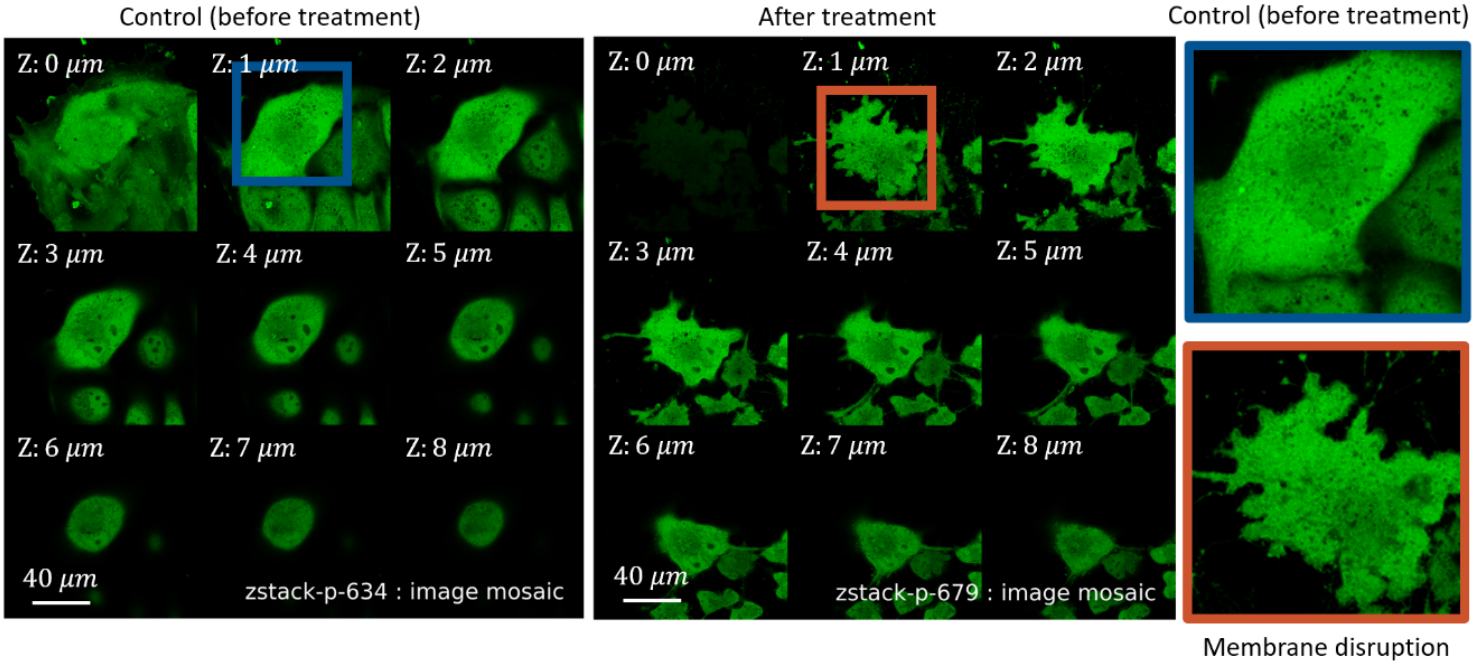


**Fig. S8.** Confocal fluorescence Z-stack images of an MDCK living cell before and after F-actin network disruption using Latrunculin-A treatment (1μM). The cell plasma membrane is strongly disrupted, possibly due to the degradation of cortical F-actin structures. Contrast adjustment is different before and after drug treatment.

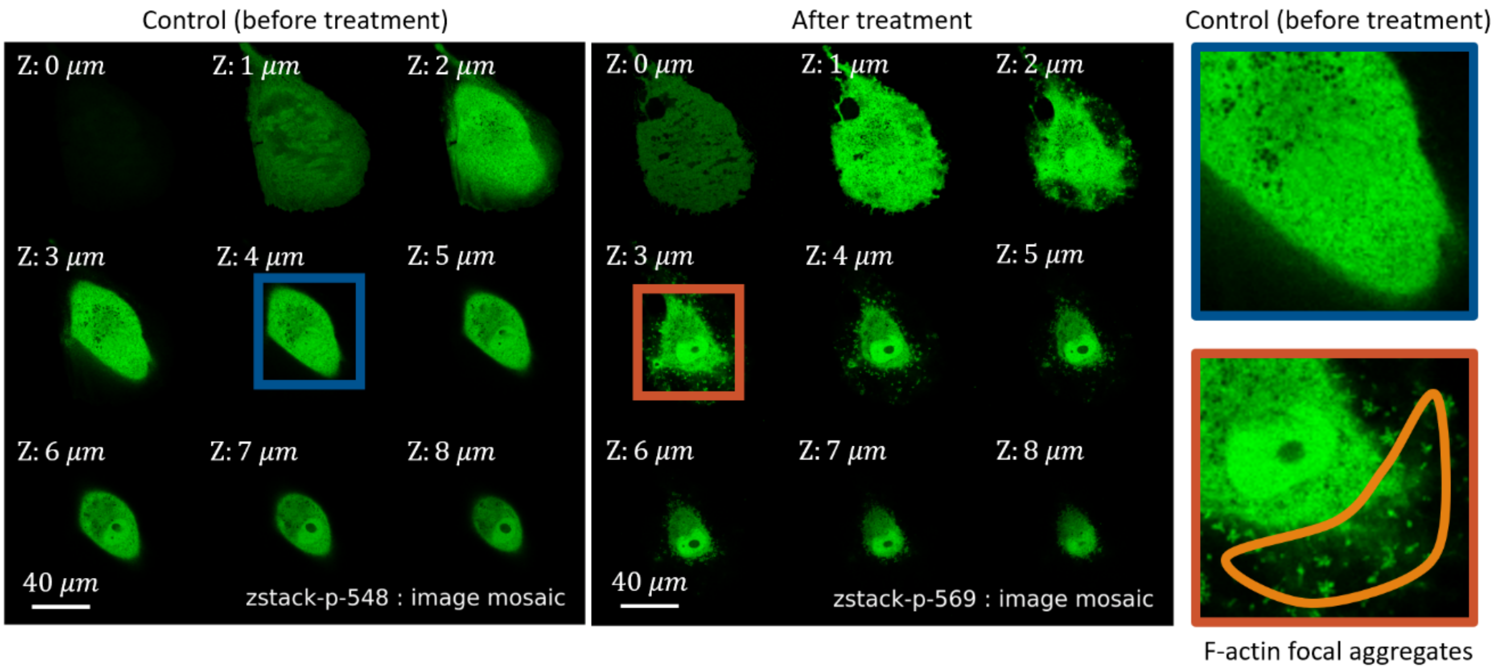


**Fig. S9.** Confocal fluorescence Z-stack images of an MDCK living cell before and after F-actin disruption using cytochalasin-D (2μM). F-actin forms aggregates. Contrast adjustment is different before and after drug treatment.

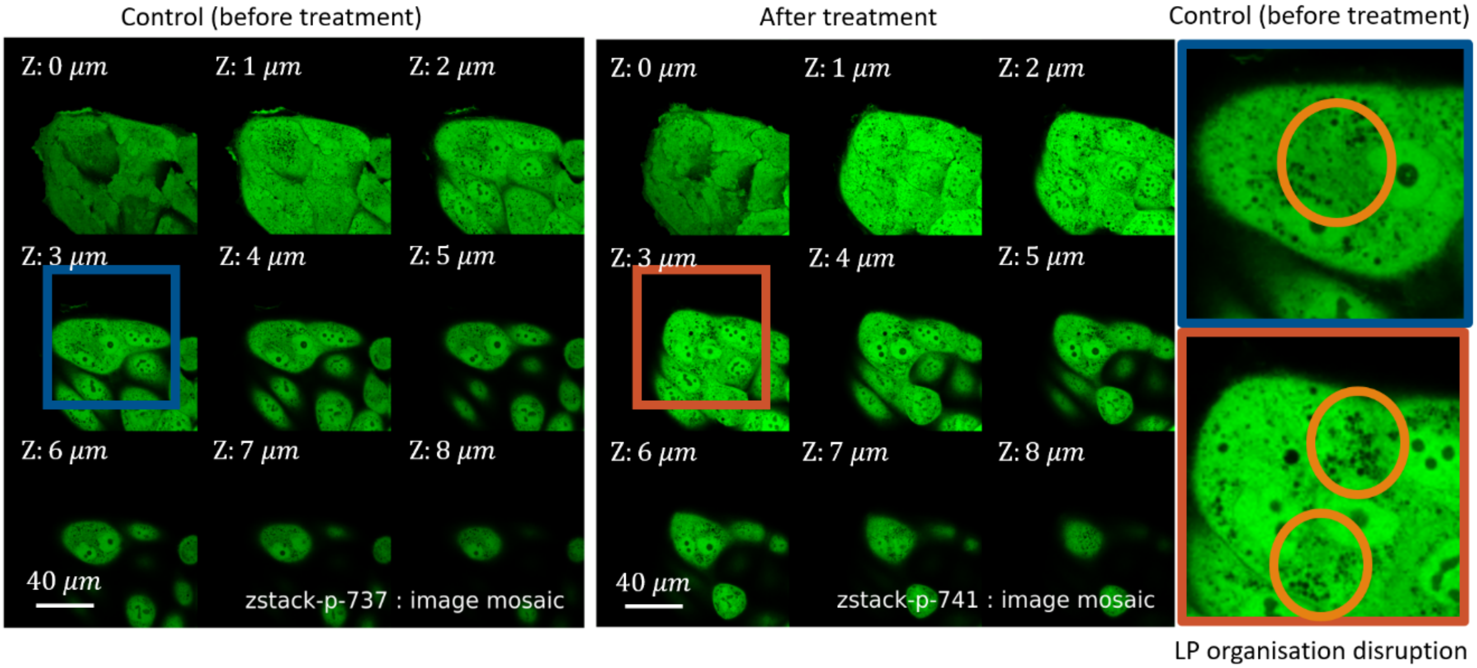


**Fig. S10.** Confocal fluorescence Z-stack images of an MDCK living cell before and after microtubule network disruption using Nocodazole treatment (2μM). The location of LP and HP regions is impacted after treatment. Contrast adjustment is different before and after drug treatment.

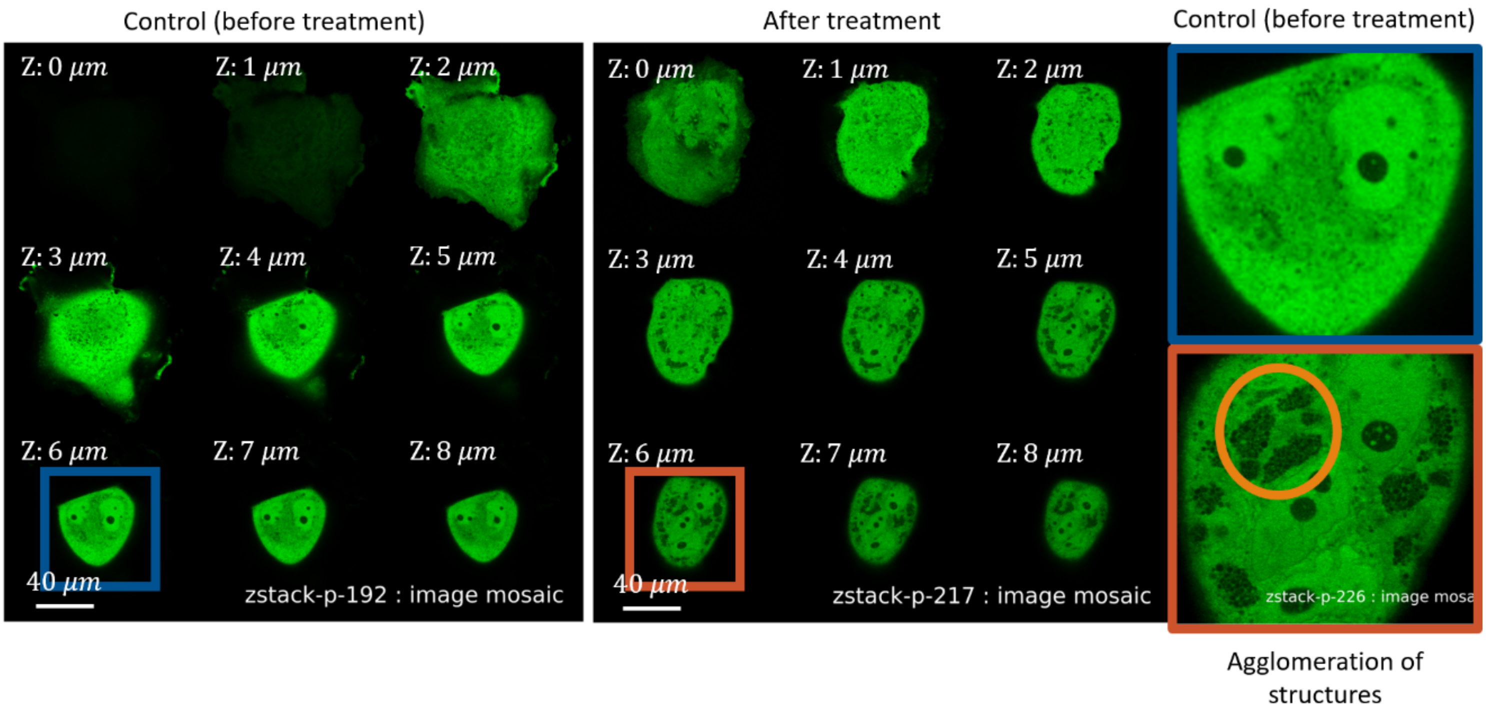


**Fig. S11.** Confocal fluorescence Z-stack images of an MDCK living cell before and after microtubule network disruption using Nocodazole treatment (10μM). Structure aggregates form at diverse locations in the cytoplasm, and the HP/LP regions can no longer be defined unambiguously. Contrast adjustment is different before and after drug treatment.

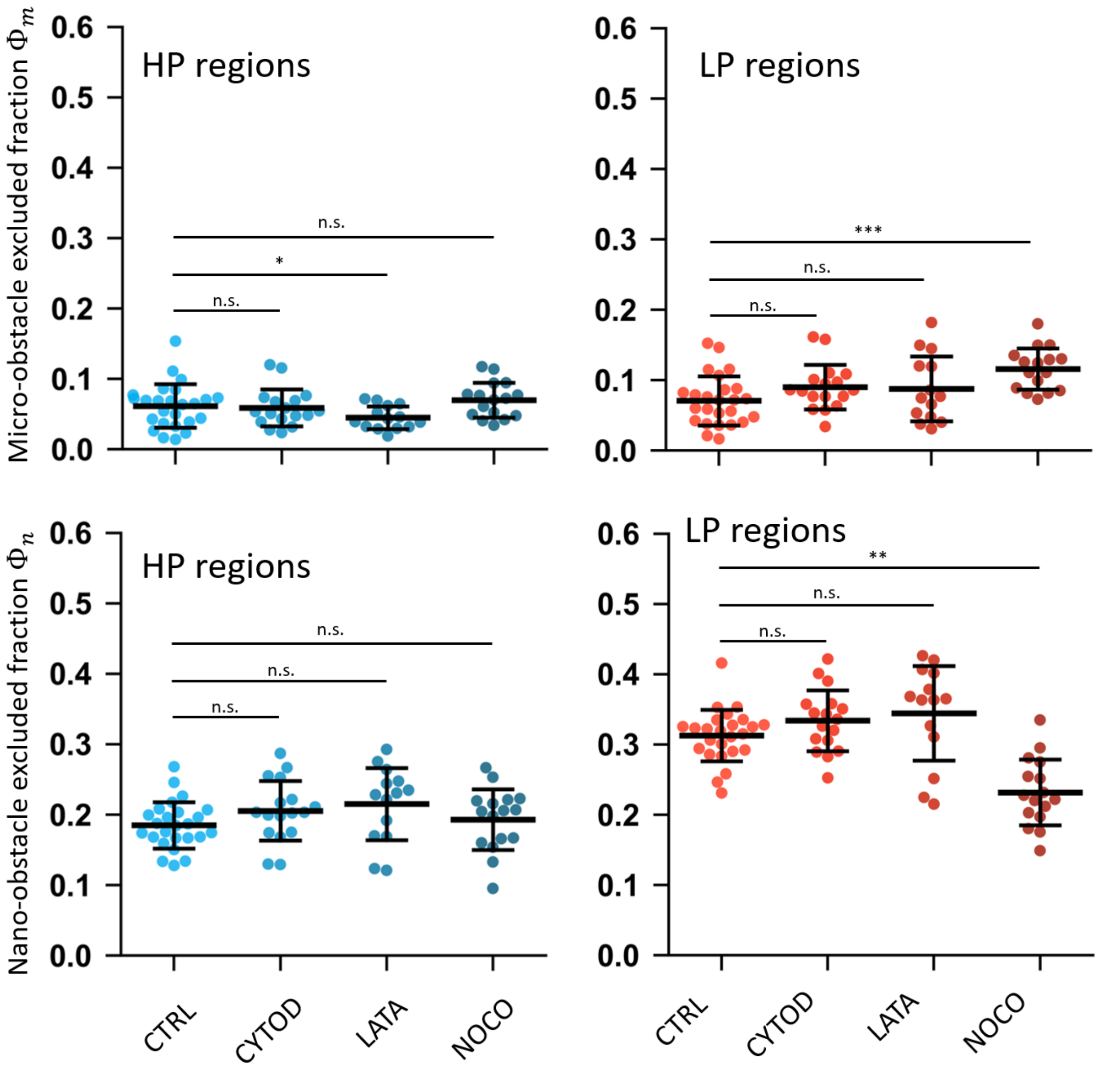


**Fig. S12.** Micro- and nano- obstacles excluded volume fractions in HP and LP regions for control cells, Cytochalasin-D (2 µM), Latrunculin-A (1 µM), and Nocodazole (2µM) drug treatments. N cells: 25 CTRL, 17 CYTOD, 14 LATA, 16 NOCO. Porosity reference shift with osmotic condition was not taken into account (as if $V_r = 1$ after drug treatment). Horizontal bars are mean and ±STD. Welch's t-test: *p < 5e − 2 ; **p < 1e − 2; ***p < 1e − 3 ; ****p < 1e − 4.

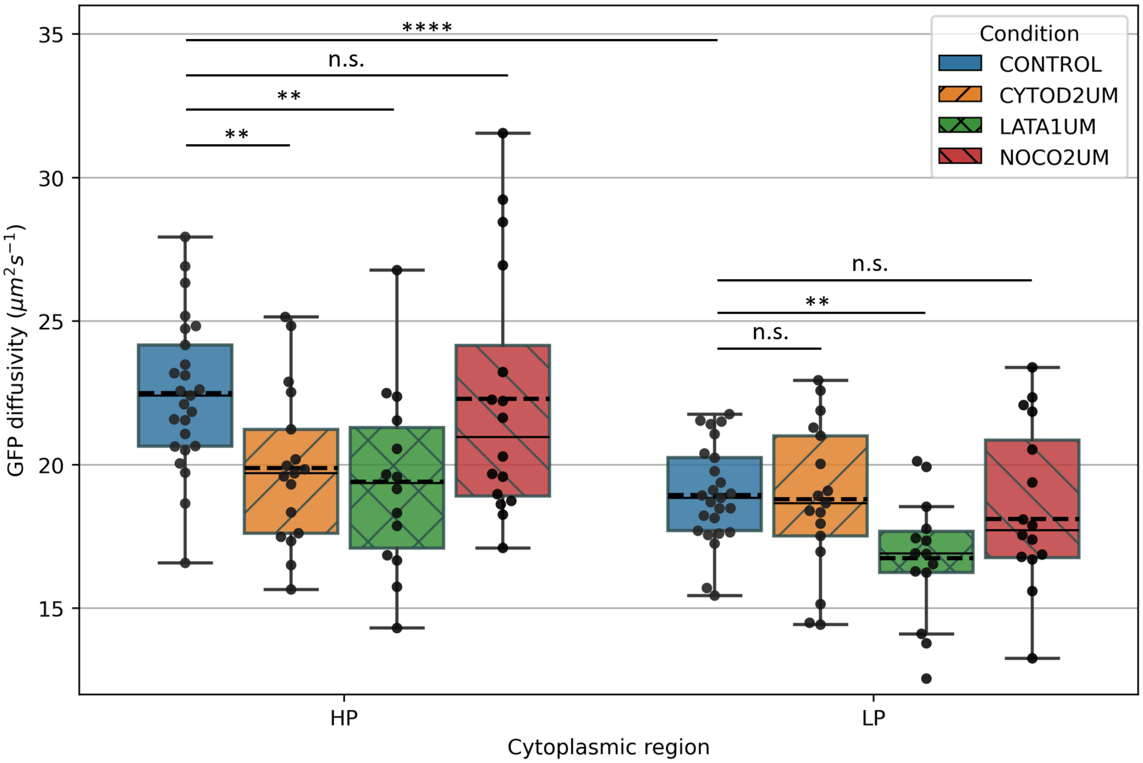


**Fig. S13.** GFP diffusivity measured by FRAP in HP and LP regions for control cells, Cytochalasin-D (2 μM), Latrunculin-A (1 μM), and Nocodazole (2μM) drug treatments. N cells: 25 CTRL, 17 CYTOD, 14 LATA, 16 NOCO. Horizontal bars are mean and ±STD. Welch's t-test: *p < 5e − 2 ; **p < 1e − 2; ***p < 1e − 3 ; ****p < 1e − 4.

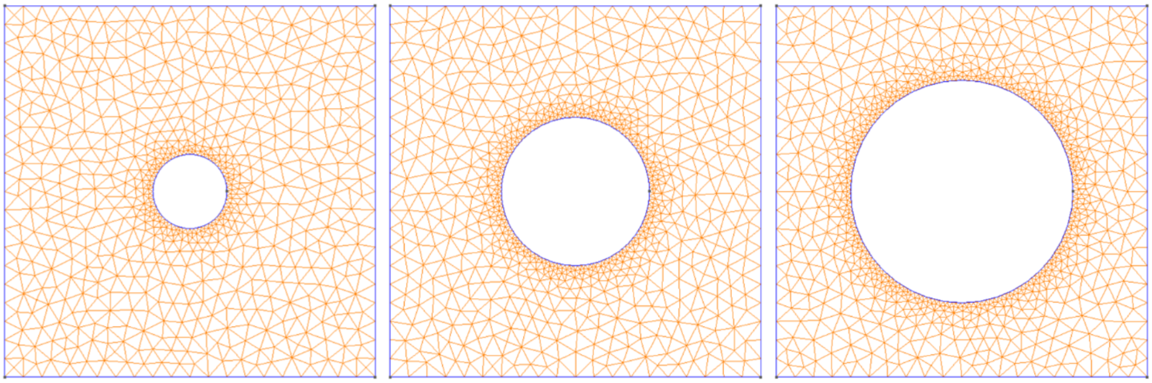

**Fig. S14.** Generated 2D mesh for the 2D-CD geometry for three disk radii: 0.1 (left), 0.2 (middle), 0.3 (right). Square size is 1.0. The 2D cytosol domain is in orange, and the 1D boundaries are in blue.

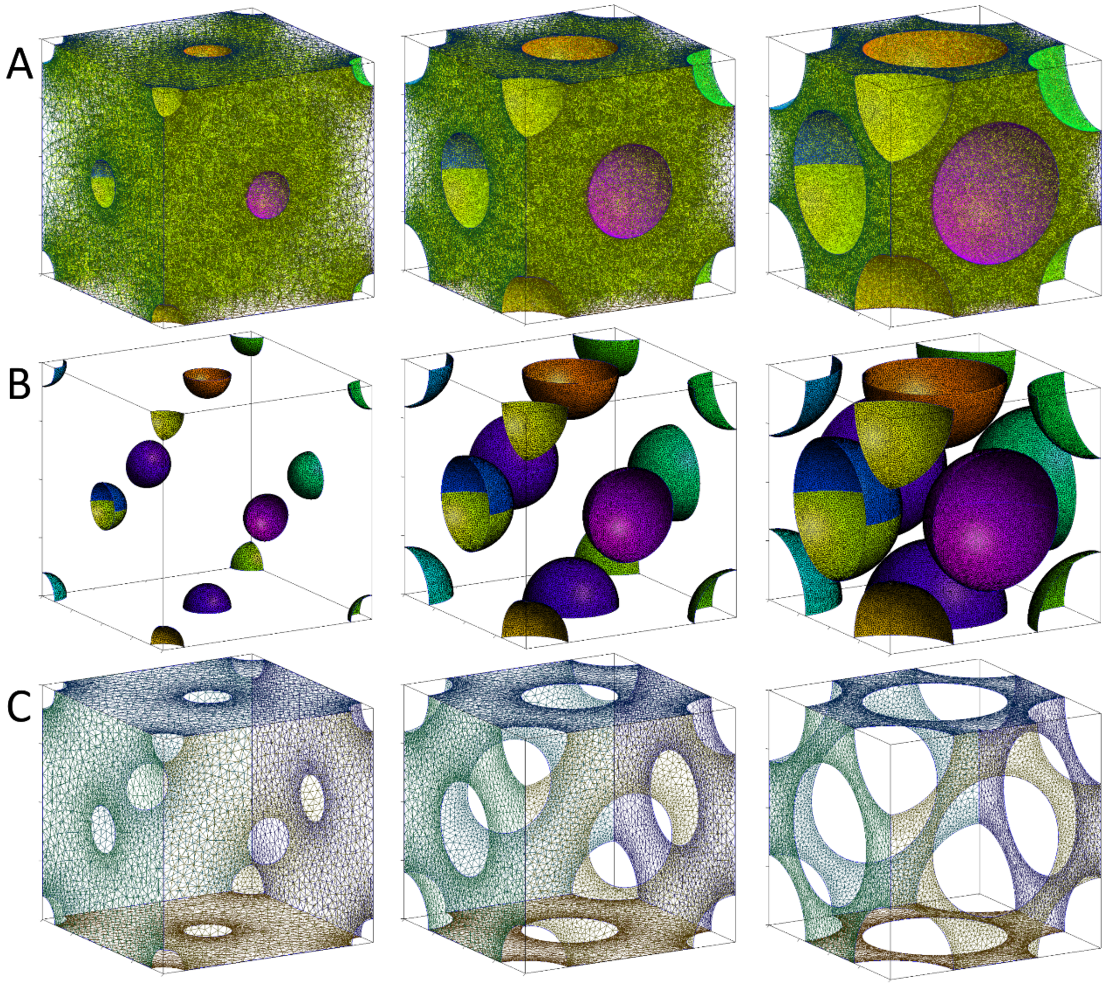


**Fig. S15.** Generated 3D mesh for the 3D-CFC geometry for three fiber radii: 0.1 (left), 0.2 (middle), 0.3 (right). The cube size is 1.0. The first line (A) corresponds to the fluid domain tetrahedral mesh (in green). The second line (B) corresponds to the obstacle surface triangular mesh (various colors). The third line (C) corresponds to the cube surface periodic triangular mesh (in black).

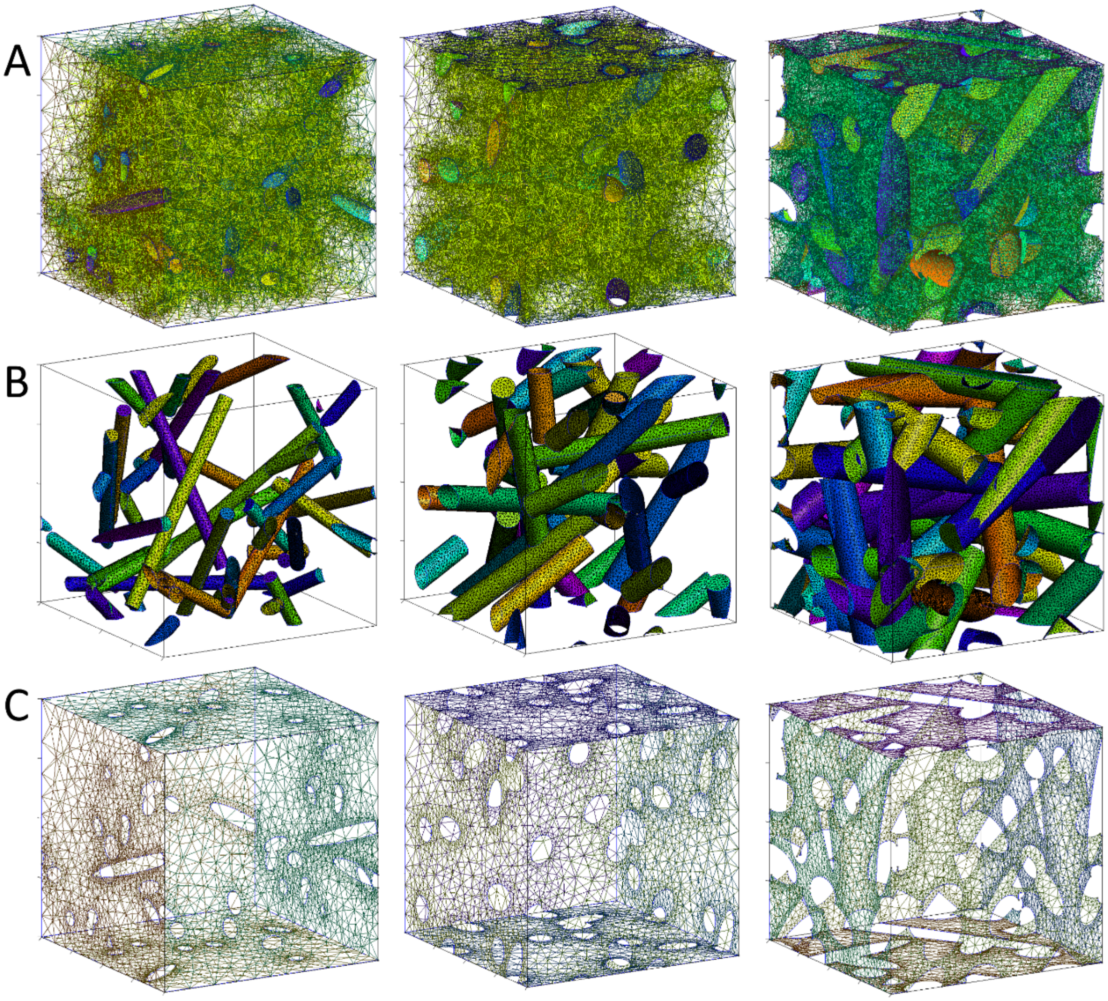


**Fig. S16.** Generated 3D mesh for the 3D-random-F-actin geometry for three fiber radii: 0.03 (left), 0.05 (middle), 0.07 (right). There are 20 fibers, their length is 0.95, and cube size is 1.0. The first line (A) corresponds to the fluid domain tetrahedral mesh (in green). The second line (B) corresponds to the obstacle surface triangular mesh (various colors). The third line (C) corresponds to the cube surface periodic triangular mesh (in black).

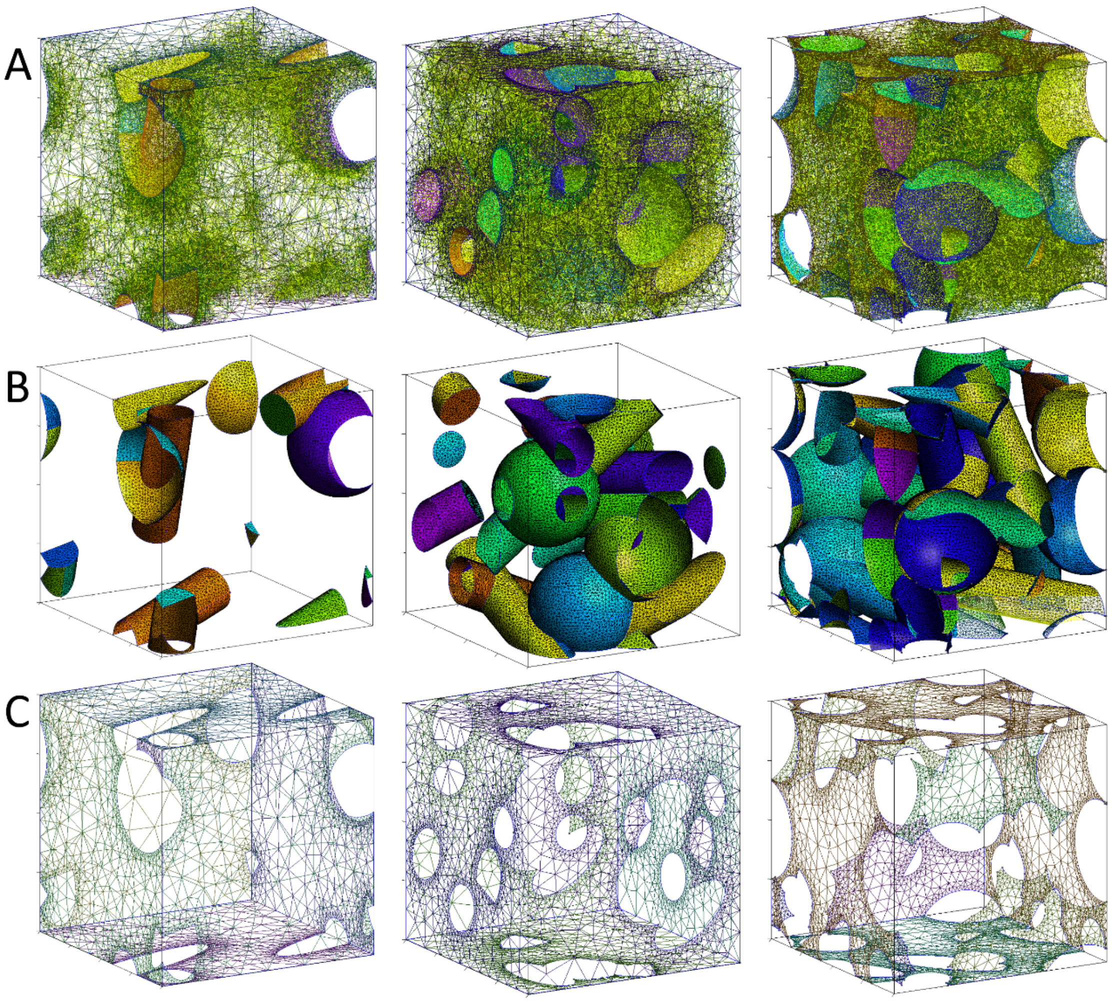


**Fig. S17.** Generated 3D mesh for the 3D-random-mixed geometry for different numbers of obstacles: 2 fibers and one sphere (left), six fibers and three spheres (middle), and ten fibers and five spheres (right). Fibers radius and length are respectively 0.06 and 0.95, sphere radius is 0.1845, and cube size is 1.0. The first line (A) corresponds to the fluid domain tetrahedral mesh (in green). The second line (B) corresponds to the obstacle surface triangular mesh (various colors). The third line (C) corresponds to the cube surface periodic triangular mesh (in black).

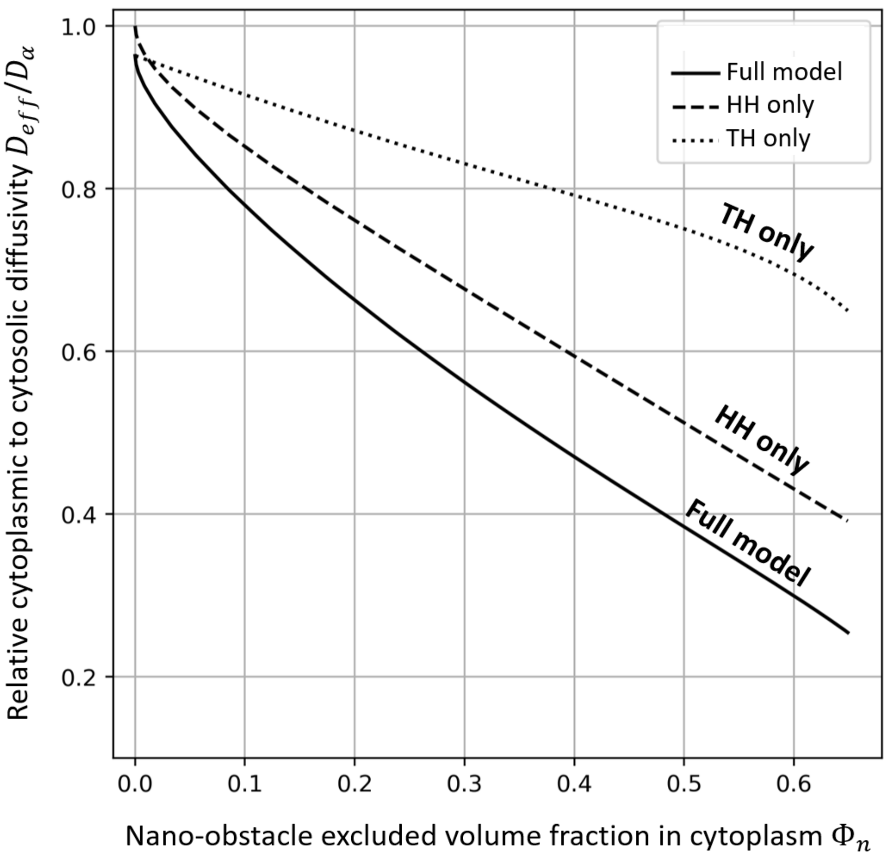


**Fig. S18.** Full model, porous Hydrodynamic Hindrance only, and Tortuous Hindrance only predictions for $D_{eff}/D_{\alpha}$ as a function of $\Phi_n$, for $\Phi_m = 0.08$, for 3D-CFC ribosome nano-scale REV and 3D-CFC micro-scale REV. These functions are used for fitting procedure with experimental results in Fig. 5A, B, C, E.

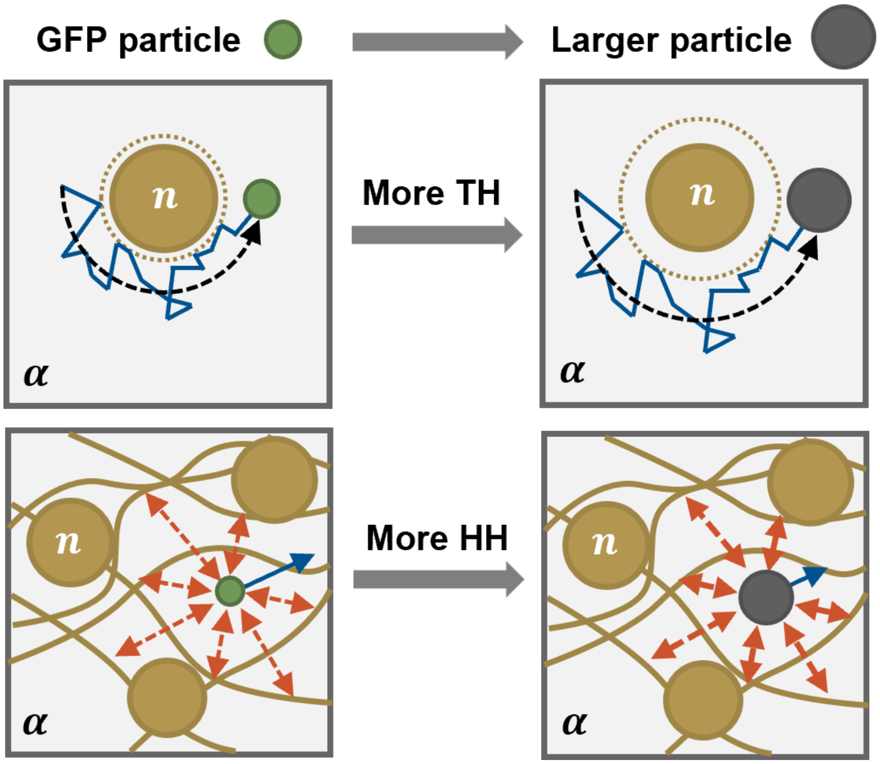


**Fig. S19.** Schematic of how a larger particle radius increases both the tortuous and the porous hydrodynamic diffusional hindrances.

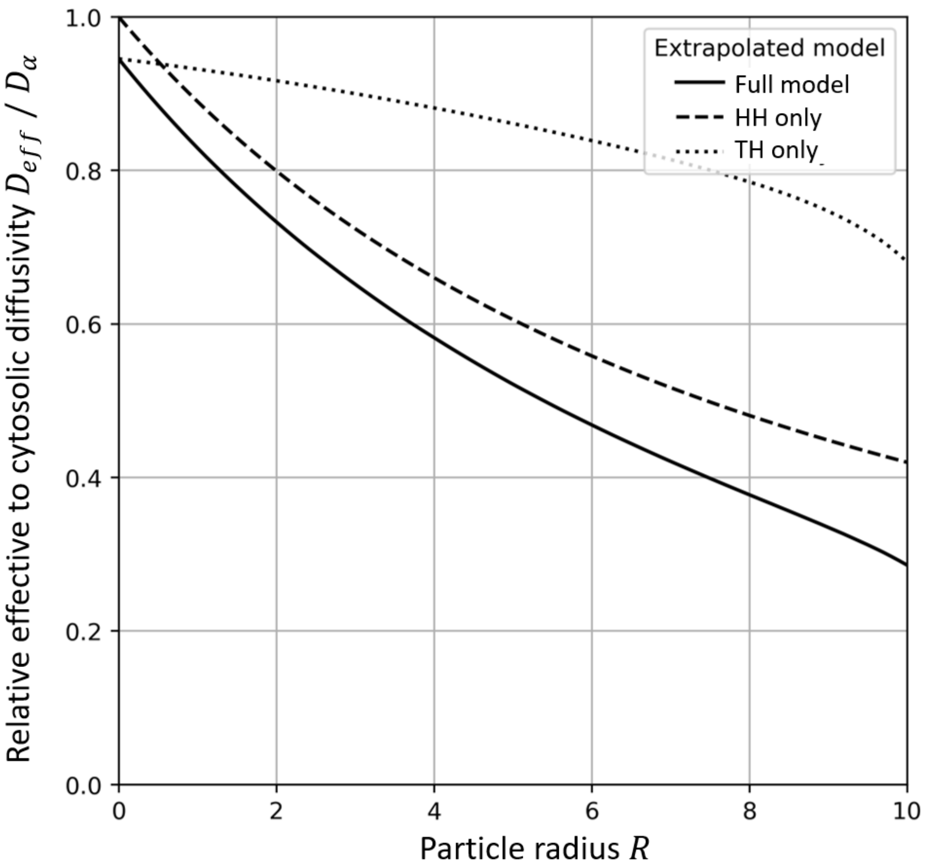


**Fig. S20.** Multiscale model predictions for dependence of relative effective cytoplasmic to cytosolic diffusivity with diffusive particle radius for $\Phi_m = 0.08$, and $\Phi_n = 0.12$.

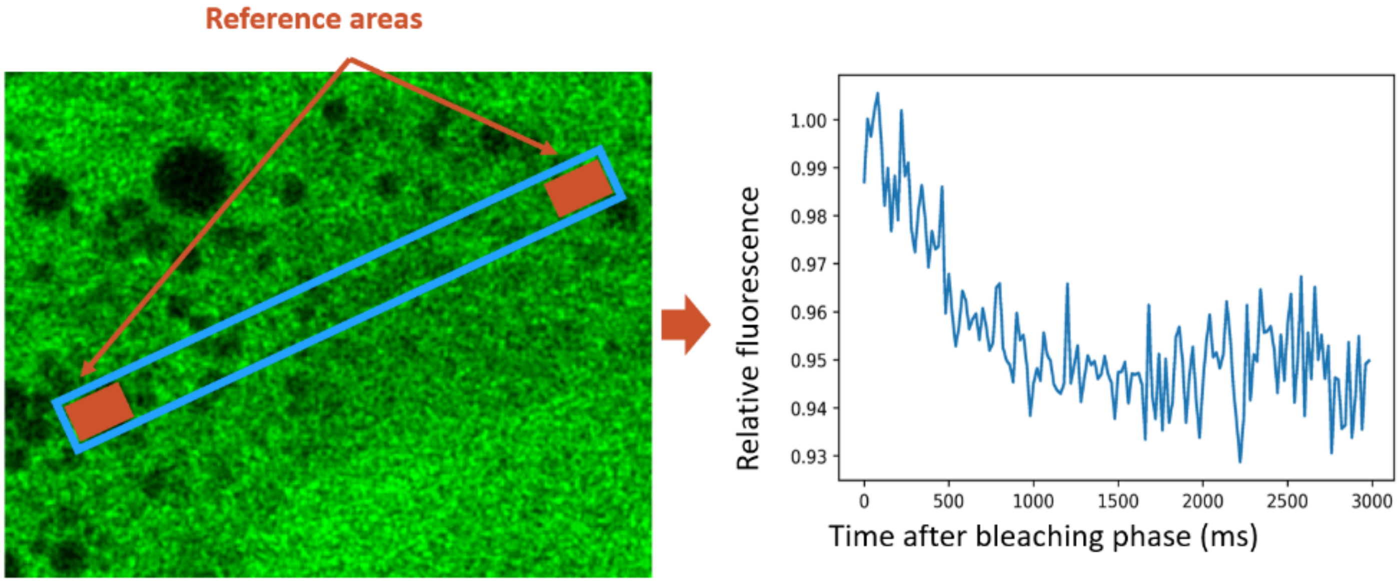


**Fig. S21.** The borders of the image rectangle are used as a reference area for correction of photobleaching during imaging. Left: confocal GFP fluorescence inside a MDCK cell. Top right: zoom on the rectangle imaged during the FRAP experiment. The rectangle borders are used as a reference area (in red). Bottom right: fluorescence signal during the post-bleach imaging phase in the reference area.

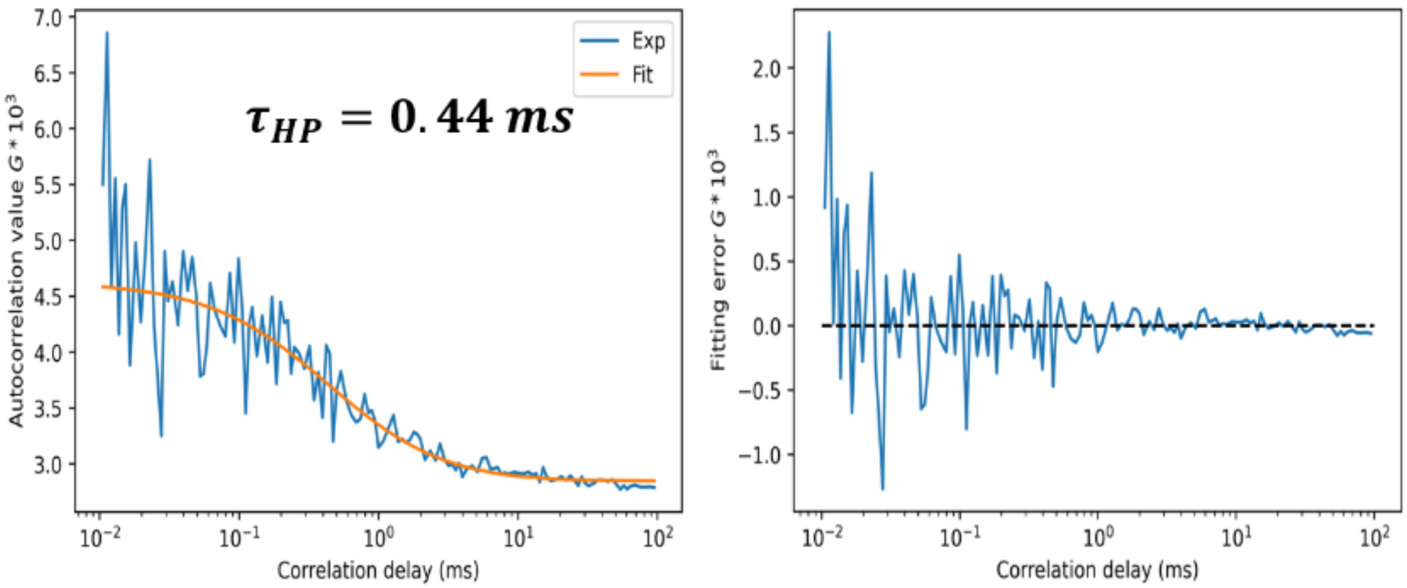


**Fig. S22.** Example of FCS simple diffusion model fitting. On the left, experimental averaged autocorrelation curve from a HP region of a cell is presented and fitted. On the right, the fitting error is plotted. Importantly, the fitting error is centered on zero for all timescales, indicating that the fit is physically relevant.

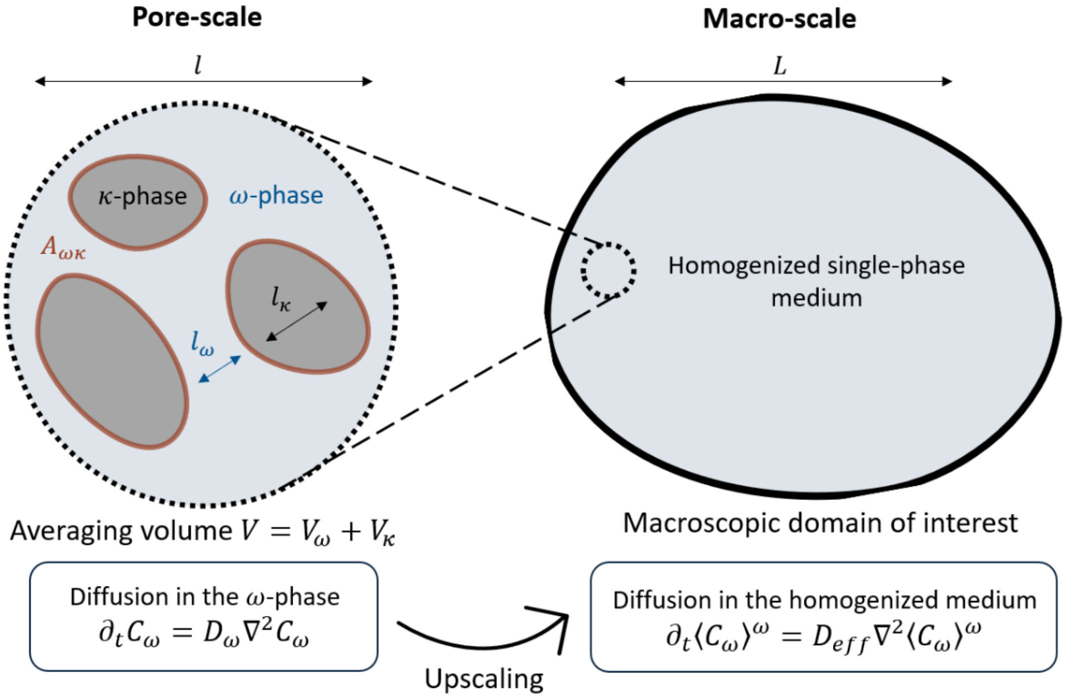


**Fig. S23.** Schematic of the upscaling for passive solutal diffusion in a homogeneous porous medium. $D_\omega$ is the local solute diffusivity in the fluid $\omega$-phase, $D_{eff}$ is the effective solute diffusivity at a much larger scale, in the homogenized medium. $\langle C_\omega \rangle^\omega$ is the intrinsic average of the solute concentration $C_\omega$ in the $\omega$-phase.

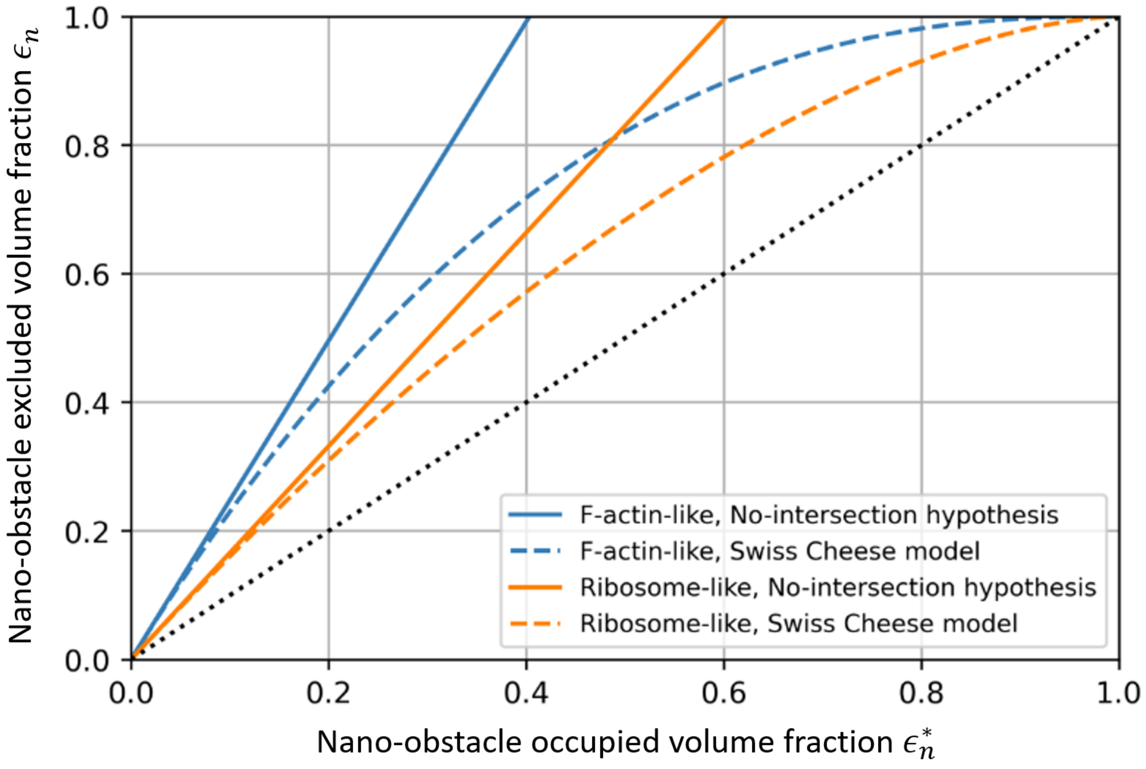


**Fig. S24.** Dependance of $\epsilon_n$ on $\epsilon_n^*$ for free-GFP, for F-actin like nano-obstacles and ribosome-like nano-obstacles. The black dotted line corresponds to of $\epsilon_n = \epsilon_n^*$.

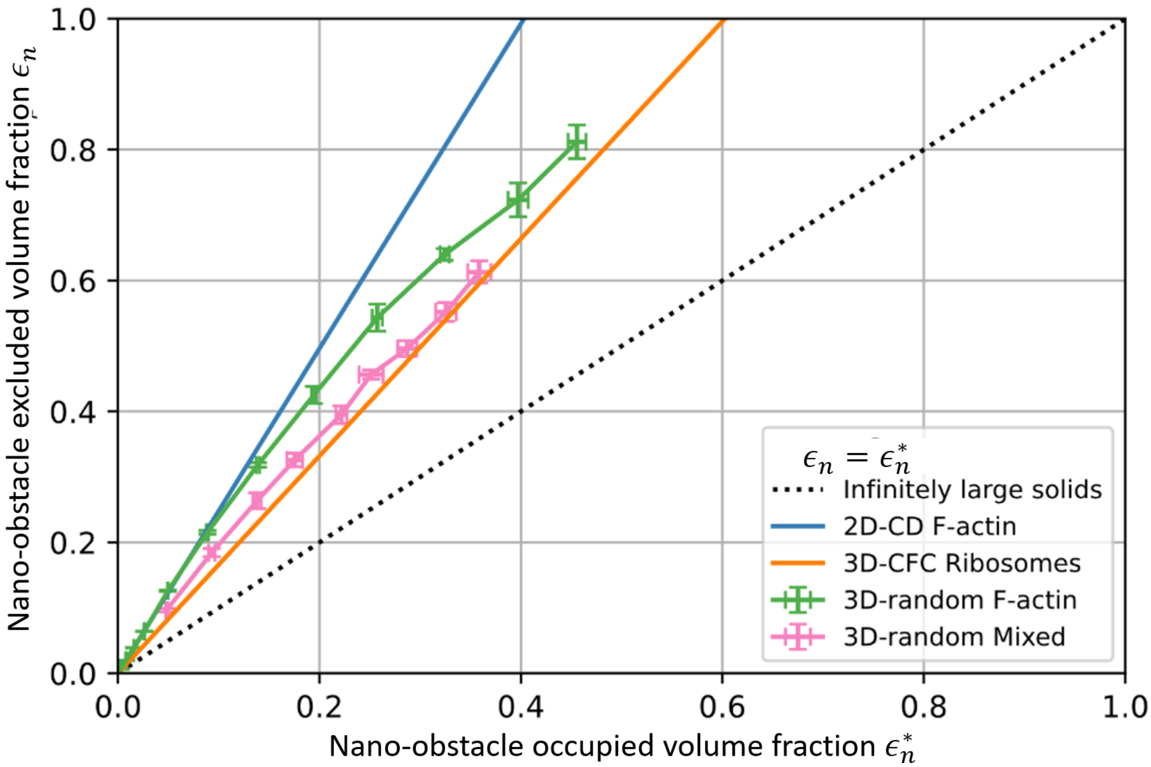


**Fig. S25.** Dependance of $\epsilon_n$ on $\epsilon_n^*$ for free-GFP, for 2D-CD-F-actin, 3D-CFC-Ribosomes, 3D-random-F-actin, and 3D-random-Mixed REV geometries. The black dotted line corresponds to of $\epsilon_n = \epsilon_n^*$.

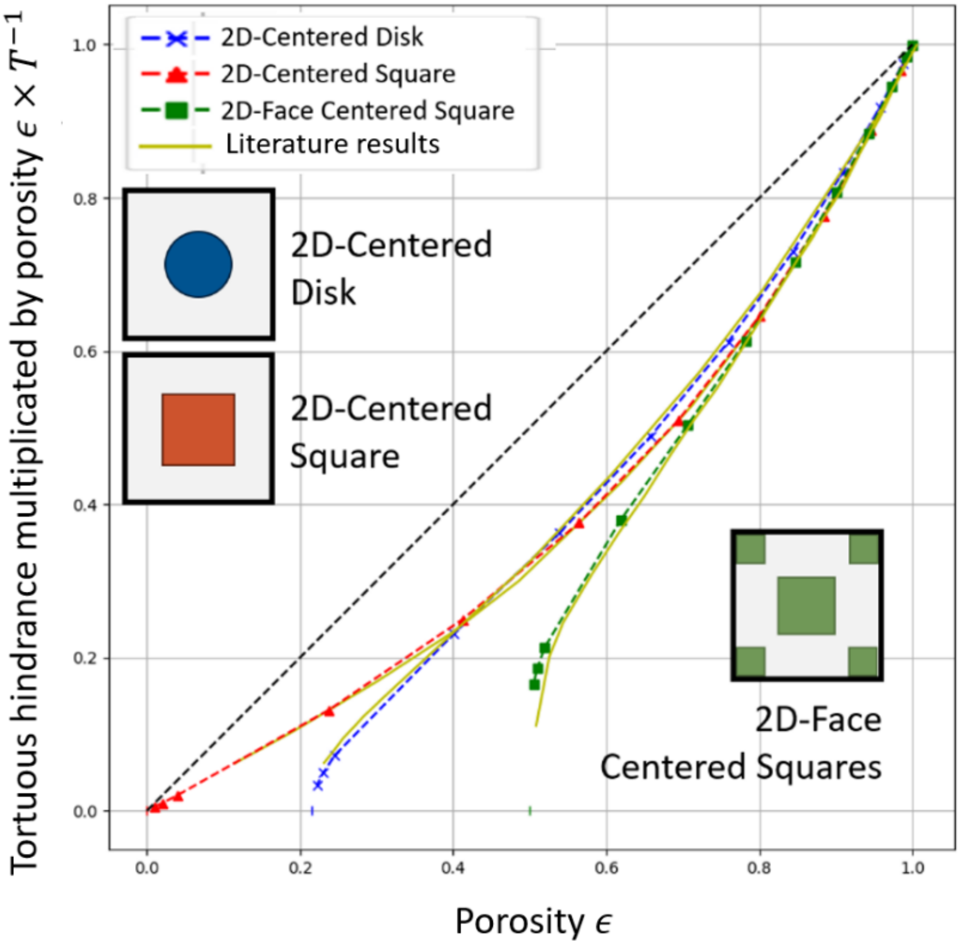


**Fig. S26.** Comparison between our results from our numerical resolution of the tortuosity closure problem with results from the literature [33]. Three REV geometries were tested: 2D Centered Disk, 2D Centered Square, and 2D Face Centered Squares. The curves from the literature (in yellow), obtained using Monte-Carlo methods, are also present for each geometry and closely match our results.

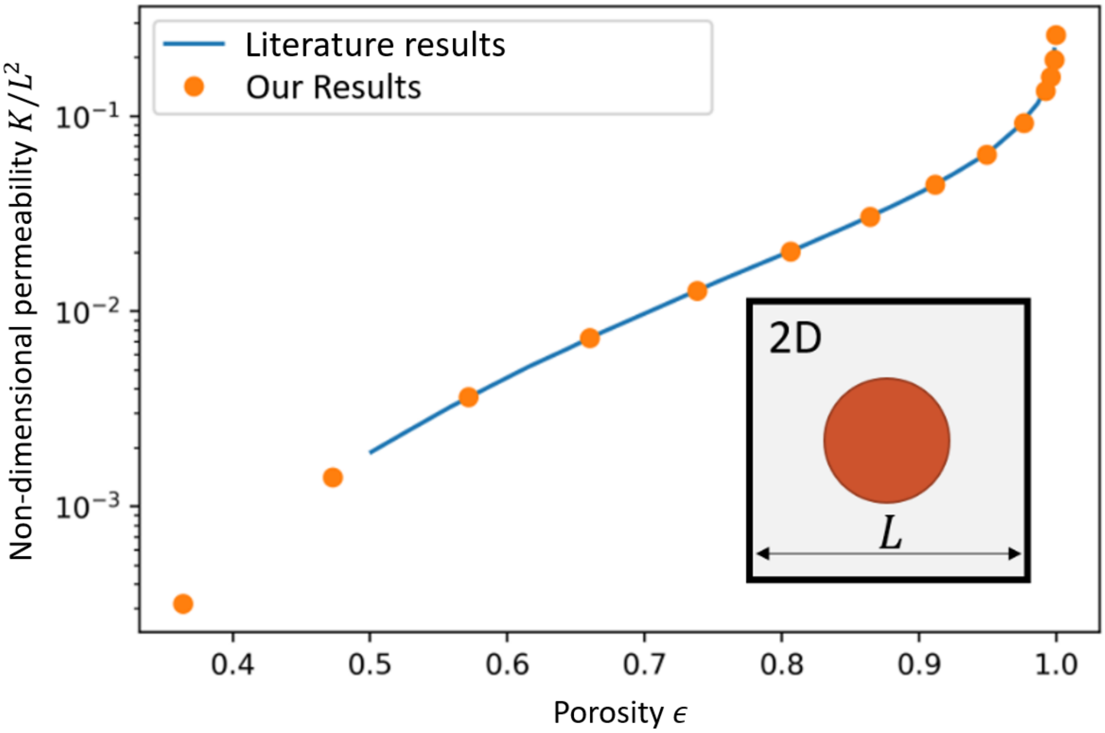


**Fig. S27.** Comparison between our results from our numerical resolution of the permeability closure problem for 2D-CD REV geometry with results from the literature [34].

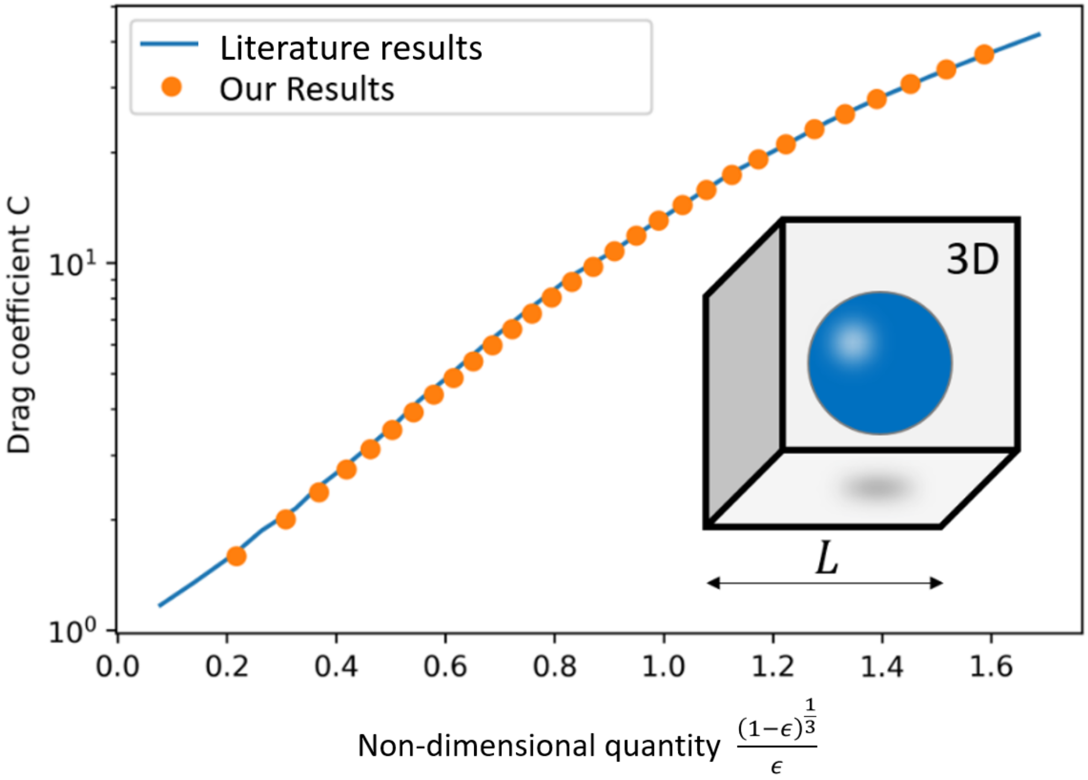


**Fig. S28.** Comparison between our results from our numerical resolution of the permeability closure problem for 3D Centered Sphere REV geometry with results from the literature [35].

| Fitting data and parameters | | | Fitted $D_\alpha$ or $D_\alpha^0$ $[\mu m^2 s^{-1}]$ | | | Fitting RMSE $[\mu m^2 s^{-1}]$ | | |
|---|---|---|---|---|---|---|---|---|
| Osmolarity | $\epsilon_{nucleus}^0$ | $k'$ | Full model | HH only | TH only | Full model | HH only | TH only |
| 300 | **0.80** | / | $30.0 \pm 0.73$ | $25.4 \pm 0.59$ | $21.3 \pm 0.49$ | $0.39 \pm 0.26$ | $0.87 \pm 0.51$ | $1.58 \pm 0.88$ |
| 450 | **0.80** | / | $27.6 \pm 0.73$ | $22.4 \pm 0.55$ | $17.5 \pm 0.40$ | $0.32 \pm 0.15$ | $0.65 \pm 0.28$ | $1.23 \pm 0.53$ |
| 600 | **0.80** | / | $24.2 \pm 0.75$ | $19.1 \pm 0.56$ | $14.3 \pm 0.40$ | $0.83 \pm 0.32$ | $0.97 \pm 0.37$ | $1.22 \pm 0.48$ |
| all | **0.80** | 1.0 | $30.0 \pm 0.50$ | $24.6 \pm 0.40$ | $19.5 \pm 0.31$ | $0.79 \pm 0.33$ | $1.20 \pm 0.46$ | $2.05 \pm 0.79$ |
| 300 | **0.90** | / | $26.1 \pm 0.64$ | $23.1 \pm 0.54$ | $20.5 \pm 0.47$ | $0.19 \pm 0.15$ | $0.75 \pm 0.44$ | $1.49 \pm 0.83$ |
| 450 | **0.90** | / | $23.3 \pm 0.63$ | $19.9 \pm 0.49$ | $16.7 \pm 0.38$ | $0.17 \pm 0.09$ | $0.57 \pm 0.25$ | $1.16 \pm 0.50$ |
| 600 | **0.90** | / | $20.3 \pm 0.62$ | $16.8 \pm 0.49$ | $13.6 \pm 0.38$ | $0.75 \pm 0.30$ | $0.92 \pm 0.35$ | $1.19 \pm 0.47$ |
| all | **0.90** | 1.0 | $25.7 \pm 0.43$ | $22.0 \pm 0.36$ | $18.6 \pm 0.29$ | $0.76 \pm 0.32$ | $1.25 \pm 0.50$ | $2.04 \pm 0.79$ |
| all | 0.85 | **0.5** | $27.0 \pm 0.45$ | $22.7 \pm 0.36$ | $18.5 \pm 0.29$ | $0.91 \pm 0.38$ | $1.42 \pm 0.56$ | $2.24 \pm 0.86$ |
| all | 0.85 | **1.5** | $28.2 \pm 0.47$ | $23.6 \pm 0.38$ | $19.3 \pm 0.31$ | $0.75 \pm 0.33$ | $1.14 \pm 0.45$ | $1.95 \pm 0.75$ |

**Table S1.** List of supplementary theory-experiment comparisons, for 3D-CFC ribosome nano-scale geometry, with associated fitting parameters and fitting RMS error. Osmolarity refers to which osmotic conditions datasets are $(\Phi_n, D_{exp})$ are used for fitting procedure. If "Osmolarity"="all", the 300, 450 and 600mOsm data are used, and the fitted diffusivity is $D_\alpha^0$. Else the fitted diffusivity is $D_\alpha$. $\boldsymbol{\epsilon_{nucleus}^0}$ refers to the nucleoplasm accessible volume at iso-osmotic condition, and $\boldsymbol{k'}$ refers to the constant used for cytosolic diffusivity variation with the osmotic condition. Standard deviations were estimated using bootstrapping accordingly with SI Appendix 4.b, with the difference that the number of resampling was reduced to 100 to speed up computations.

| Fitting data and parameters | | | Fitted $D_\alpha$ or $D_\alpha^0$ $[\mu m^2 s^{-1}]$ | | | Fitting RMSE $[\mu m^2 s^{-1}]$ | | |
|---|---|---|---|---|---|---|---|---|
| Osmolarity | $\epsilon_{nucleus}^0$ | $k'$ | Full model | HH only | TH only | Full model | HH only | TH only |
| 300 | 0.85 | / | $34.6 \pm 0.94$ | $27.3 \pm 0.66$ | $23.1 \pm 0.53$ | $0.48 \pm 0.33$ | $0.46 \pm 0.29$ | $1.10 \pm 0.62$ |
| 450 | 0.85 | / | $33.3 \pm 1.00$ | $24.3 \pm 0.62$ | $19.6 \pm 0.46$ | $0.23 \pm 0.14$ | $0.43 \pm 0.20$ | $0.90 \pm 0.39$ |
| 600 | 0.85 | / | $30.2 \pm 0.99$ | $20.9 \pm 0.35$ | $16.3 \pm 0.47$ | $0.61 \pm 0.26$ | $0.89 \pm 0.35$ | $1.08 \pm 0.41$ |
| all | 0.85 | 1.0 | $35.9 \pm 0.64$ | $26.7 \pm 0.44$ | $21.7 \pm 0.35$ | $0.76 \pm 0.41$ | $0.96 \pm 0.39$ | $1.59 \pm 0.62$ |
| 300 | **0.80** | / | $38.3 \pm 1.02$ | $29.0 \pm 0.69$ | $24.0 \pm 0.55$ | $0.27 \pm 0.22$ | $0.59 \pm 0.36$ | $1.18 \pm 0.67$ |
| 450 | **0.80** | / | $37.4 \pm 1.09$ | $26.2 \pm 0.66$ | $20.5 \pm 0.48$ | $0.08 \pm 0.09$ | $0.53 \pm 0.23$ | $0.95 \pm 0.41$ |
| 600 | **0.80** | / | $33.9 \pm 1.12$ | $22.5 \pm 0.67$ | $17.0 \pm 0.49$ | $0.68 \pm 0.28$ | $0.94 \pm 0.36$ | $1.10 \pm 0.42$ |
| all | **0.80** | 1.0 | $40.0 \pm 0.71$ | $28.6 \pm 0.47$ | $22.6 \pm 0.36$ | $0.84 \pm 0.45$ | $1.00 \pm 0.40$ | $1.60 \pm 0.61$ |
| 300 | **0.90** | / | $31.2 \pm 0.87$ | $25.6 \pm 0.62$ | $22.3 \pm 0.51$ | $0.71 \pm 0.47$ | $0.32 \pm 0.21$ | $1.00 \pm 0.57$ |
| 450 | **0.90** | / | $29.6 \pm 0.93$ | $22.6 \pm 0.59$ | $18.8 \pm 0.45$ | $0.40 \pm 0.22$ | $0.33 \pm 0.15$ | $0.83 \pm 0.36$ |
| 600 | **0.90** | / | $27.0 \pm 0.89$ | $19.5 \pm 0.58$ | $15.6 \pm 0.44$ | $0.54 \pm 0.24$ | $0.84 \pm 0.33$ | $1.05 \pm 0.41$ |
| all | **0.90** | 1.0 | $32.1 \pm 0.59$ | $24.9 \pm 0.41$ | $20.8 \pm 0.33$ | $0.77 \pm 0.39$ | $0.92 \pm 0.38$ | $1.56 \pm 0.62$ |
| all | 0.85 | **0.5** | $35.0 \pm 0.63$ | $26.0 \pm 0.43$ | $21.1 \pm 0.34$ | $0.62 \pm 0.28$ | $1.14 \pm 0.46$ | $1.79 \pm 0.70$ |
| all | 0.85 | **1.5** | $36.4 \pm 0.66$ | $27.1 \pm 0.44$ | $22.1 \pm 0.35$ | $0.92 \pm 0.51$ | $0.91 \pm 0.38$ | $1.50 \pm 0.58$ |

**Table S2.** List of supplementary theory-experiment comparisons for 2D-CD F-actin nano-scale geometry, with associated fitting parameters and fitting RMS error. Osmolarity refers to which osmotic conditions datasets are $(\Phi_n, D_{exp})$ are used for fitting procedure. If "Osmolarity"="all", the 300, 450 and 600mOsm data are used, and the fitted diffusivity is $D_\alpha^0$. Else the fitted diffusivity is $D_\alpha$. $\boldsymbol{\epsilon_{nucleus}^0}$ refers to the nucleoplasm accessible volume at iso-osmotic condition, and $\boldsymbol{k'}$ refers to the constant used for cytosolic diffusivity variation with the osmotic condition. Standard deviations were estimated using bootstrapping accordingly with SI Appendix 4.b, with the difference that the number of resampling was reduced to 100 to speed up computations.

**Movie S1 (separate file).** Free-GFP fluorescence confocal Z-stack of a MDCK cell representative of the ones used in our experiments.
